# Betulonic acid derivatives inhibiting coronavirus replication in cell culture via the nsp15 endoribonuclease

**DOI:** 10.1101/2020.12.10.418996

**Authors:** Besir Krasniqi, Annelies Stevaert, Benjamin Van Loy, Tien Nguyen, Joice Thomas, Julie Vandeput, Dirk Jochmans, Volker Thiel, Ronald Dijkman, Wim Dehaen, Arnout Voet, Lieve Naesens

**Affiliations:** Molecular Design and Synthesis, Department of Chemistry, KU Leuven, 3001 Leuven, Belgium; Laboratory of Virology and Chemotherapy, Rega Institute, KU Leuven, 3000 Leuven, Belgium; Biochemistry, Molecular and Structural Biology, Department of Chemistry, KU Leuven, 3001 Leuven, Belgium; Institute of Virology and Immunology (IVI), 3012 Bern and 3147 Mittelhäusern, Switzerland; Department of Infectious Diseases and Pathobiology, Vetsuisse Faculty, University of Bern, 3012 Bern, Switzerland; Institute for Infectious Diseases (IFIK), University of Bern, 3012 Bern, Switzerland

**Keywords:** coronavirus, antiviral, nsp15, endoribonuclease, betulonic acid

## Abstract

The lack of medication to suppress coronavirus infections is a main reason for the dramatic course of the COVID-19 pandemic. There is an urgent need to identify suitable coronavirus drug targets and corresponding lead molecules. Here we describe the discovery of a class of coronavirus inhibitors acting on nsp15, a hexameric protein component of the viral replication-transcription complexes, endowed with immune evasion-associated endoribonuclease activity. SAR exploration of these 1,2,3-triazolo fused betulonic acid derivatives yielded lead molecule **5h** as a strong inhibitor (antiviral EC_50_: 0.6 μM) of human coronavirus 229E replication. An nsp15 endoribonuclease active site mutant virus was markedly less sensitive to **5h**, and selected resistance to the compound mapped to mutations in the N-terminal part of nsp15, at an interface between two nsp15 monomers. The biological findings were substantiated by the nsp15 binding mode for **5h**, predicted by docking. Hence, besides delivering a distinct class of inhibitors, our study revealed a druggable pocket in the nsp15 hexamer with relevance for anti-coronavirus drug development.

## INTRODUCTION

The current SARS-CoV-2 pandemic is causing a major crisis in terms of human health and socio-economic losses. Within a period of ~20 years, SARS-CoV-2 is the third zoonotic coronavirus (CoV) to enter the human species, coming after SARS (Severe Acute Respiratory Syndrome) and MERS (Middle East Respiratory Syndrome).^1^ Besides, four human CoVs (i.e. HCoV-229E, -HKU1, -NL63, and -OC43) are endemic in the population and account each year for 15 to 30% of common colds.^2^ These can evolve into life-threatening lower respiratory tract infections in elderly, children and persons at risk.^3, 4^ Finally, the *Coronaviridae* family contains several species causing serious disease in pets and livestock.^5^

Most young persons infected with SARS-CoV-2 experience no or only mild symptoms of respiratory illness.^6^ In contrast, in persons with comorbidities or higher age, the viral replication phase is typically followed by a second phase that is characterized by hyperinflammation, acute respiratory distress syndrome and multi-organ failure.^7^ Hence, management of COVID-19 most likely requires antiviral drugs to suppress initial virus replication, plus anti-inflammatory medication, like corticosteroids, to treat severe cases.^8^ Several CoV proteins may be suitable drug targets,^9, 10^ but, at the moment, only two drug classes are far advanced in clinical testing, i.e. anti-spike antibodies^11^ and the nucleotide analogue remdesivir, which inhibits the CoV polymerase.^12-14^ Though less explored, the CoV nsp15 endoribonuclease (EndoU) is a highly attractive drug target, since it has no cellular counterpart, its catalytic site is conserved among CoVs, and it is amenable to structure-based design based on available protein structures.^15-19^ Nsp15 is one of the non-structural proteins (nsps) associated with the replication-transcription complexes (RTCs), the site where CoV RNA synthesis occurs.^5, 20^ Although the functions of nsp15 are not entirely understood, its EndoU function is known to regulate viral RNA synthesis, limit the recognition of viral dsRNA by cellular sensors and prevent the dsRNA-activated antiviral host cell response.^21-25^ The concept to inhibit nsp15 is thus unique, since it combines a direct antiviral effect with the potential to revert viral evasion from host cell immunity.

We here report identification of a class of coronavirus nsp15 inhibitors with 1,2,3-triazolo fused betulonic acid structure. We describe their synthesis, structure-activity relationship and the mechanistic findings, in particular resistance data, that corroborate nsp15 as their antiviral target. These biological data accord with the binding model that we obtained by compound docking in the hexameric nsp15 protein structure. The model also explains why the current lead is active against human coronavirus 229E, but not other coronaviruses like SARS-CoV-2. In short, our study validates the nsp15 protein, and particularly the interface where the lead compound binds, as a druggable and pertinent target for developing CoV inhibitors.

## RESULTS AND DISCUSSION

### Compound synthesis and structure-activity relationship

The 1,2,3-triazolo fused betulonic acid derivatives (**Scheme 1**) were designed and synthesized in the context of a pharmacological hit discovery program. Betulonic acid bears a pentacyclic triterpenoid core, present in a wide variety of agents with potential pharmacological use,^26-28^ like the HIV maturation inhibitor bevirimat.^29, 30^ We decided to fuse betulonic acid with a 1,2,3-triazole moiety, which has the unique property to both accept and donate hydrogen bonds.^31-34^ These derivatives were synthesized by our recently developed and convenient “triazolization” method to prepare 1,2,3-triazoles from primary amines and ketones.^35-38^ First, Jones oxidation was performed to convert betulin **1** into betulonic acid **2** (**Scheme 1**).^39^ Betulin **1**, a natural compound isolated from the bark of *Betula* species, is commercially available.^40-42^ Next, the triazolization method was applied to betulonic acid **2** as the ketone source, using primary amines **3** and 4-nitrophenyl azide **4**, and the previously reported reaction conditions.^13^ This yielded a series of sixteen 1,2,3-triazolo fused betulonic acids **5**, most of which were isolated at high yield (~80%; Table 1). Diverse primary amines **3** were attached to the 1,2,3-triazole ring to introduce a variety of aromatic or aliphatic moieties.

**Scheme 1.**
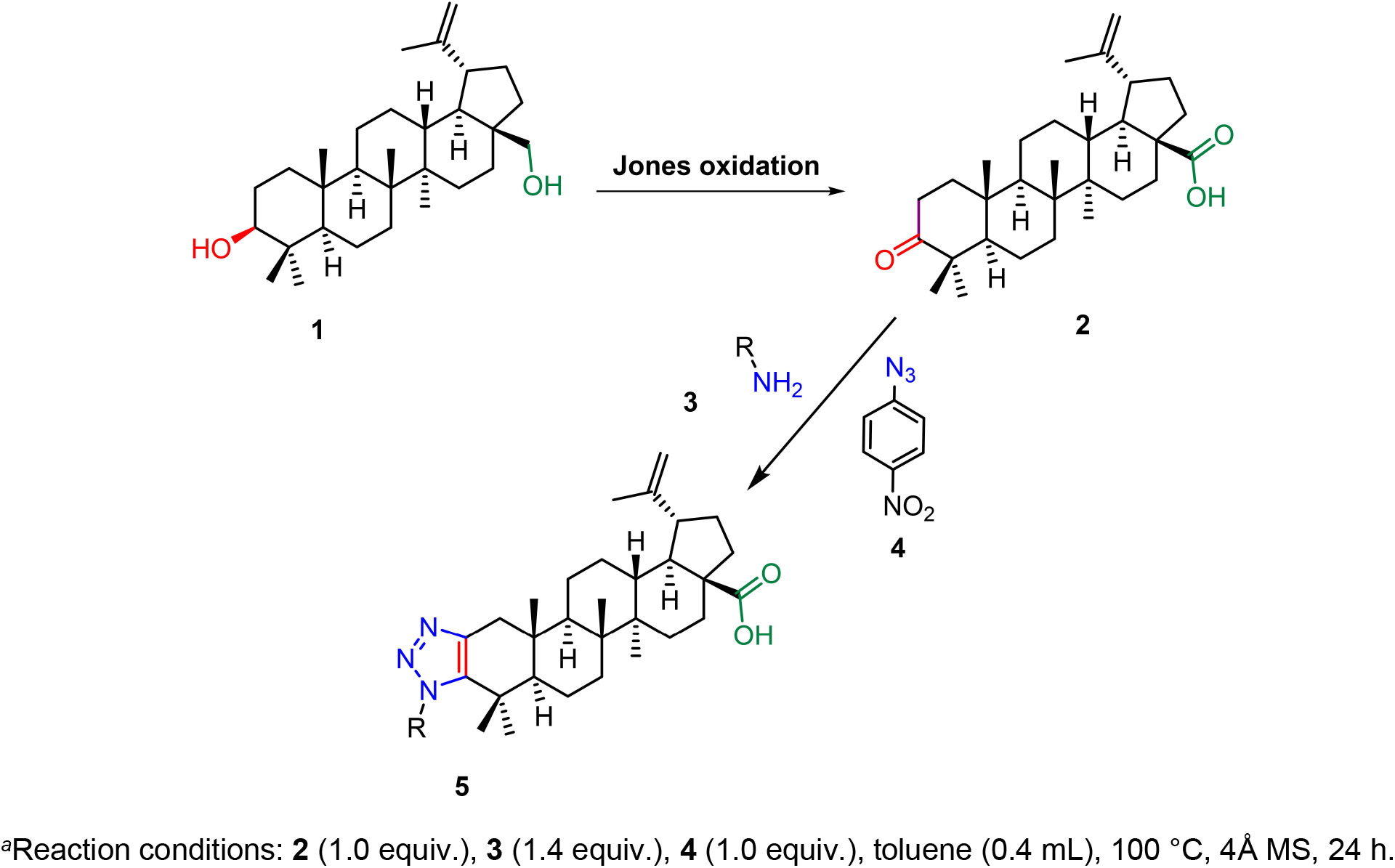
Synthesis of fused 1,2,3-triazole betulonic acid derivatives starting from betulin.^*a*^

**Table 1.**
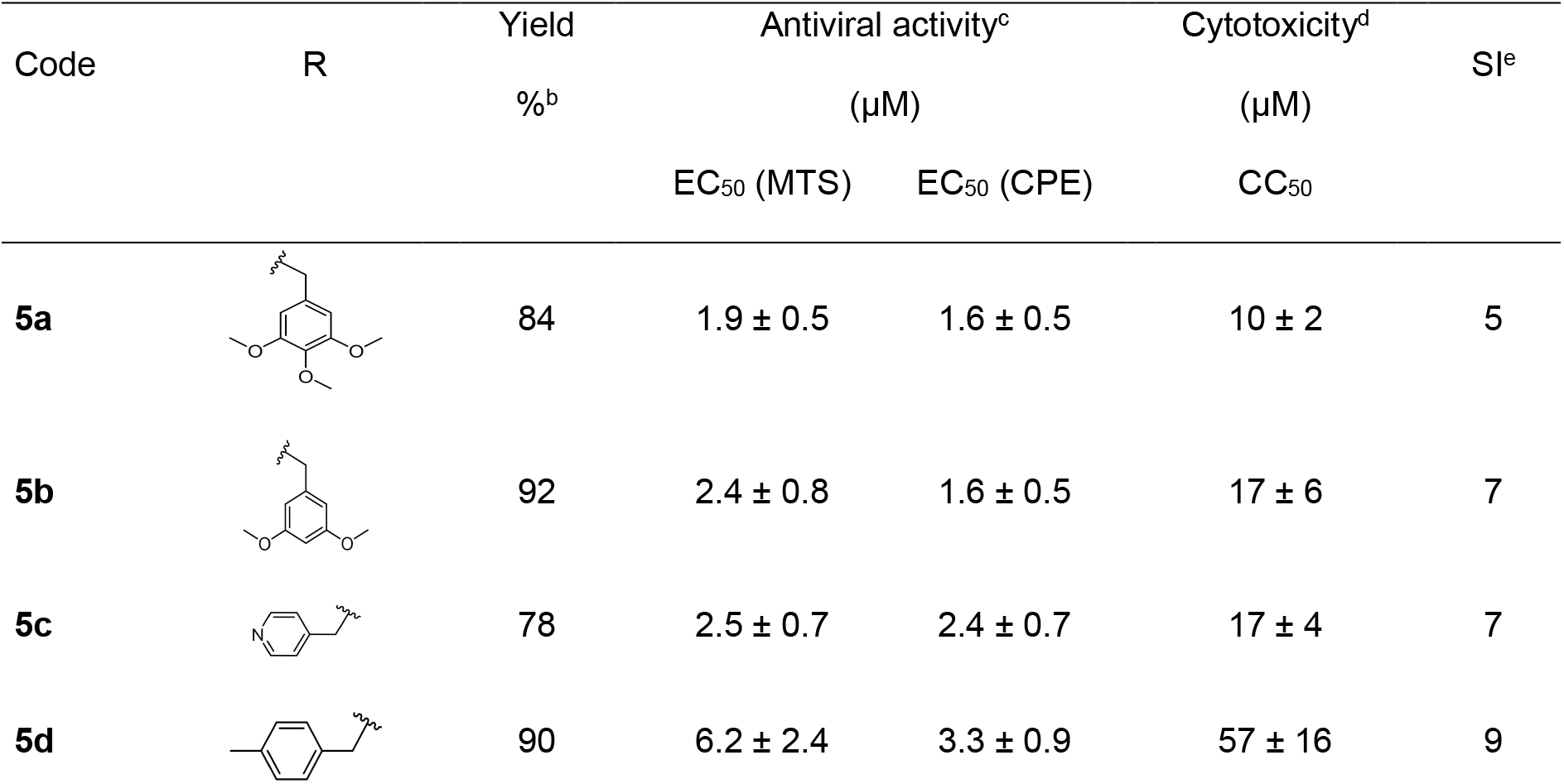
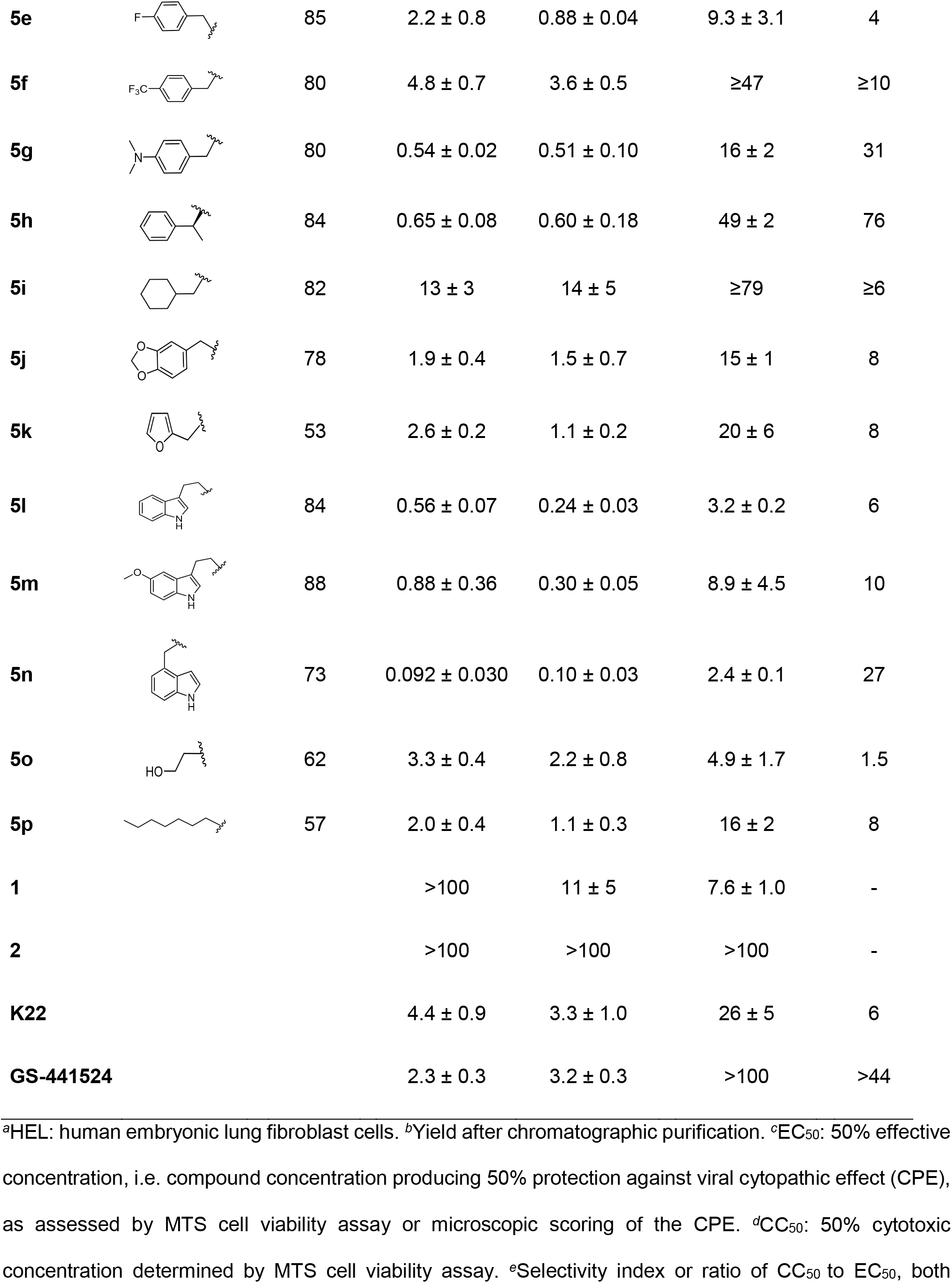
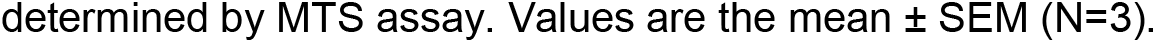
Anti-CoV activity and selectivity in human HEL^a^ cells infected with HCoV-229E.

| Code | R | Yield | Antiviral activity<sup>c</sup> |  | Cytotoxicity<sup>d</sup> | SI<sup>e</sup> |
| --- | --- | --- | --- | --- | --- | --- |
|  |  | %<sup>b</sup> | (μM) |  | (μM) |  |
|  |  |  | EC<sub>50</sub> (MTS) | EC<sub>50</sub> (CPE) | CC<sub>50</sub> |  |
| 5a |  | 84 | 1.9 ± 0.5 | 1.6 ± 0.5 | 10 ± 2 | 5 |
| 5b |  | 92 | 2.4 ± 0.8 | 1.6 ± 0.5 | 17 ± 6 | 7 |
| 5c |  | 78 | 2.5 ± 0.7 | 2.4 ± 0.7 | 17 ± 4 | 7 |
| 5d |  | 90 | 6.2 ± 2.4 | 3.3 ± 0.9 | 57 ± 16 | 9 |
| **5e** | | 85 | $2.2 \pm 0.8$ | $0.88 \pm 0.04$ | $9.3 \pm 3.1$ | 4 |
| **5f** | | 80 | $4.8 \pm 0.7$ | $3.6 \pm 0.5$ | $\geq 47$ | $\geq 10$ |
| **5g** | | 80 | $0.54 \pm 0.02$ | $0.51 \pm 0.10$ | $16 \pm 2$ | 31 |
| **5h** | | 84 | $0.65 \pm 0.08$ | $0.60 \pm 0.18$ | $49 \pm 2$ | 76 |
| **5i** | | 82 | $13 \pm 3$ | $14 \pm 5$ | $\geq 79$ | $\geq 6$ |
| **5j** | | 78 | $1.9 \pm 0.4$ | $1.5 \pm 0.7$ | $15 \pm 1$ | 8 |
| **5k** | | 53 | $2.6 \pm 0.2$ | $1.1 \pm 0.2$ | $20 \pm 6$ | 8 |
| **5l** | | 84 | $0.56 \pm 0.07$ | $0.24 \pm 0.03$ | $3.2 \pm 0.2$ | 6 |
| **5m** | | 88 | $0.88 \pm 0.36$ | $0.30 \pm 0.05$ | $8.9 \pm 4.5$ | 10 |
| **5n** | | 73 | $0.092 \pm 0.030$ | $0.10 \pm 0.03$ | $2.4 \pm 0.1$ | 27 |
| **5o** | | 62 | $3.3 \pm 0.4$ | $2.2 \pm 0.8$ | $4.9 \pm 1.7$ | 1.5 |
| **5p** | | 57 | $2.0 \pm 0.4$ | $1.1 \pm 0.3$ | $16 \pm 2$ | 8 |
| **1** | | | >100 | $11 \pm 5$ | $7.6 \pm 1.0$ | - |
| **2** |  |  | >100 | >100 | >100 | - |
| **K22** | | | $4.4 \pm 0.9$ | $3.3 \pm 1.0$ | $26 \pm 5$ | 6 |
| **GS-441524** | | | $2.3 \pm 0.3$ | $3.2 \pm 0.3$ | >100 | >44 |

<sup>a</sup>HEL: human embryonic lung fibroblast cells. <sup>b</sup>Yield after chromatographic purification. <sup>c</sup>EC<sub>50</sub>: 50% effective concentration, i.e. compound concentration producing 50% protection against viral cytopathic effect (CPE), as assessed by MTS cell viability assay or microscopic scoring of the CPE. <sup>d</sup>CC<sub>50</sub>: 50% cytotoxic concentration determined by MTS cell viability assay. <sup>e</sup>Selectivity index or ratio of CC<sub>50</sub> to EC<sub>50</sub>, both
determined by MTS assay. Values are the mean ± SEM (N=3).

We next evaluated the compounds in cell-based assays with diverse DNA- and RNA-viruses, and observed strong activity in human embryonic lung (HEL) fibroblast cells infected with HCoV-229E. We used a viral cytopathic effect (CPE) reduction assay, in which protection against CPE (expressed as the antiviral EC_50_ value) was monitored by the MTS cell viability assay and verified by microscopy. The MTS assay was also used to quantify compound cytotoxicity (expressed as the CC_50_ value) in mock-infected cells. Whereas the starting compounds betulin **1** and betulonic acid **2** were virtually inactive, all 1,2,3-triazolo fused betulonic acid derivatives proved to be highly effective CoV inhibitors (**Table 1**). The one exception was **5i** (EC_50_ value: 13 μM), which bears a non-aromatic cyclohexanemethyl substituent. Apparently, introducing this bulky group caused a considerable reduction in antiviral activity and selectivity. Several compounds in the series had EC_50_ values below 1 μM, which makes them superior to two known CoV inhibitors which we used as reference compounds, i.e. K22^43^ and GS-441524, the nucleoside form of remdesivir.^14^ Three analogues stood out for having superior selectivity, i.e. **5n**, **5g** and **5h**, having a selectivity index (ratio of CC_50_ to EC_50_) of 27, 31 and 76, respectively. The capacity of **5h** to fully suppress HCoV-229E replication at non-toxic concentrations is evident from the microscopic images in **Fig. 1A** and the dose-response curves in **Fig. 1B**. Also, **5h** fully prevented the formation of dsRNA intermediates of CoV RNA synthesis, as demonstrated by immunofluorescence staining of dsRNA in HCoV-229E-infected human bronchial epithelial 16HBE cells (**Fig. 1C**).

**Figure 1.**
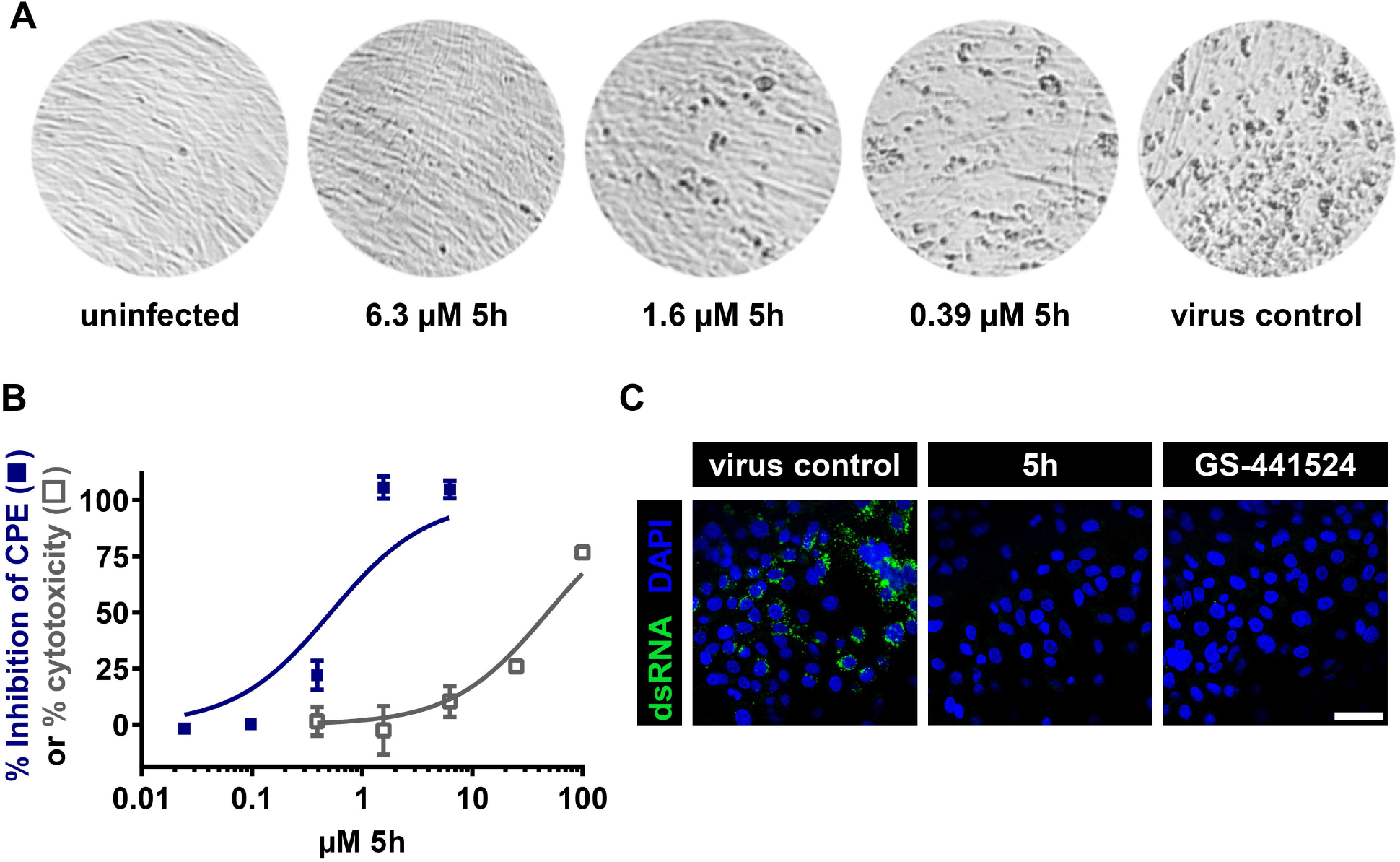
Activity of **5h** against HCoV-229E. (A) Representative microscopic images showing protection against virus-induced cytopathic effect (CPE) in human embryonic lung (HEL) cells. (B) Dose-response curves for inhibition of virus-induced CPE (◼) and for cytotoxicity (◻) in HEL cells, both determined by MTS cell viability assay. Data points are the mean ± SEM (N=3). (C) Immunofluorescence detection of viral dsRNA in HCoV-229E-infected human bronchial epithelial 16HBE cells at 24 h post-infection (p.i.). In green: dsRNA and in blue: nuclear DAPI staining. Compounds: 12 μM **5h** or 12 μM GS-441524. Scale bar: 50 μm.

To conduct a SAR exploration around lead compound **5h** (**Scheme 2** and **Table 2**), we first investigated the contribution of the α-methyl-phenylene moiety. Compound **5q**, in which this entire moiety is missing, had ~6-fold lower antiviral activity than **5h**. When only the α-methyl was missing (**5r**), the activity was not affected. Compound **5s**, which is the epimer at the 1,2,3-triazole substituent, displayed almost the same EC_50_ value as **5h**, indicating that isomerism does not alter the activity. Cytotoxicity was however slightly decreased, resulting in an even better selectivity index (≥ 90) than that of **5h**. In order to elucidate the role of the isopropenyl side chain, we reduced this moiety by hydrogenation, yielding compound **5t** which was 10- to 20-fold less active. Replacement of the carboxylic acid by a methyl group (**5u**) proved deleterious.

**Scheme 2.**
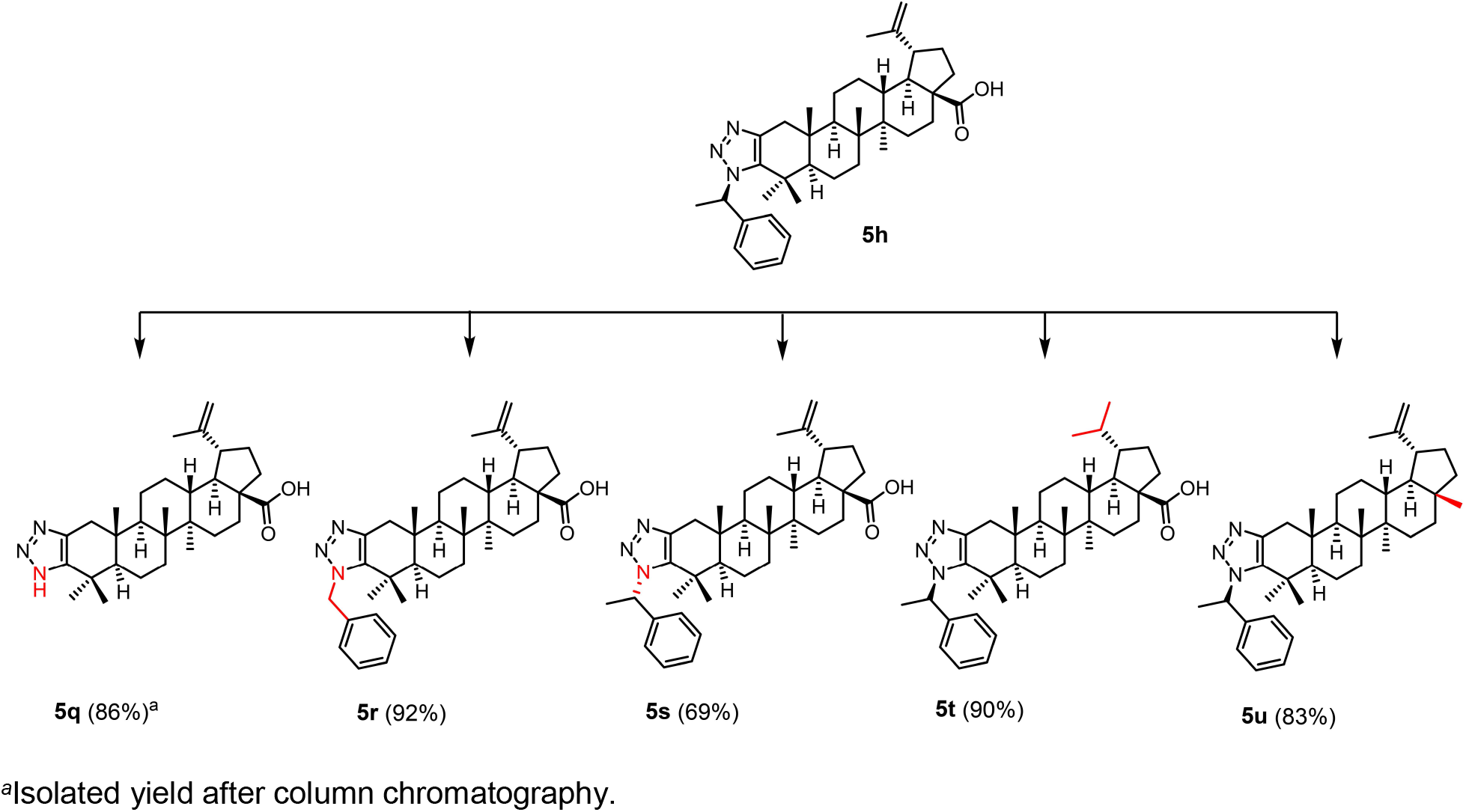
SAR study around compound **5h**.

**Table 2.** Activity of **5h** analogues in human HEL^a^ cells infected with HCoV-229E.

| Code | Antiviral activity<sup>b</sup> (μM) |  | Cytotoxicity<sup>c</sup> | SI<sup>d</sup> |
| --- | --- | --- | --- | --- |
|  |  |  | (μM) |  |
|  | EC<sub>50</sub> (MTS) | EC<sub>50</sub> (CPE) | CC<sub>50</sub> |  |
| **5q** | 4.3 ± 0.6 | 3.4 ± 0.4 | 8.3 ± 0.7 | 1.9 |
| **5r** | 0.85 ± 0.05 | 0.71 ± 0.04 | 12 ± 0 | 14 |
| **5s** | 1.1 ± 0.2 | 0.67 ± 0.02 | >100 | 91 |
| **5t** | 13 ± 5 | 6.1 ± 1.7 | ≥99 | 7.6 |
| **5u** | >100 | >100 | >100 | - |
| **5h** | 0.65 ± 0.08 | 0.60 ± 0.18 | 49 ± 2 | 76 |

*a,b,c,d* See Legend to Table 1.

We next evaluated **5h** in cell culture assays with a panel of other CoVs. The compound had no inhibitory effect on mouse hepatitis virus A59 (MHV-A59) and feline infectious peritonitis virus (FIPV), in CPE reduction assays in, respectively, murine fibroblast L2 cells and Crandell-Rees Feline Kidney cells (data not shown). HCoV-229E and FIPV belong to the alpha genus, while MHV-A59 belongs to the beta genus comprising also the highly pathogenic species SARS-CoV-1, MERS-CoV and SARS-CoV-2.^44, 45^ In VeroE6-eGFP cells infected with SARS-CoV-2, **5h** and **5t** were inactive [see reference^46^ for assay description]. Hence, though nicely active against HCoV-229E, **5h** appeared, unfortunately, to be confined to this CoV species. Besides, when tested against a broad panel of DNA and RNA viruses, the 1,2,3-triazolo fused betulonic acid derivatives proved inactive against HIV, herpes simplex virus, vaccinia virus, adenovirus, vesicular stomatitis virus, Coxsackie B4 virus, respiratory syncytial virus, parainfluenza-3 virus, reovirus-1, Sindbis virus, Punta Toro virus, yellow fever virus and influenza virus (data not shown).

### Mechanistic studies establishing nsp15 as the target of 5h

Given the robust activity of the betulonic acid derivatives against HCoV-229E, we used this virus to reveal their mechanism of action and appreciate how their anti-CoV activity spectrum may be expanded. A time-of-addition experiment indicated that **5h** acts post-entry at an early stage in viral RNA synthesis, since the molecule started to have reduced activity when added at 6 h p.i. (Fig. 2). For comparison, the action point of the entry inhibitor bafilomycin, an inhibitor of endosomal acidification, was situated before 2 h p.i. K22 was still effective when added as late as 8 h p.i. This CoV inhibitor targets nsp6-dependent anchorage of the viral replication-transcription complexes (RTCs) to host cell-derived double-membrane vesicles.^43^

**Figure 2.**
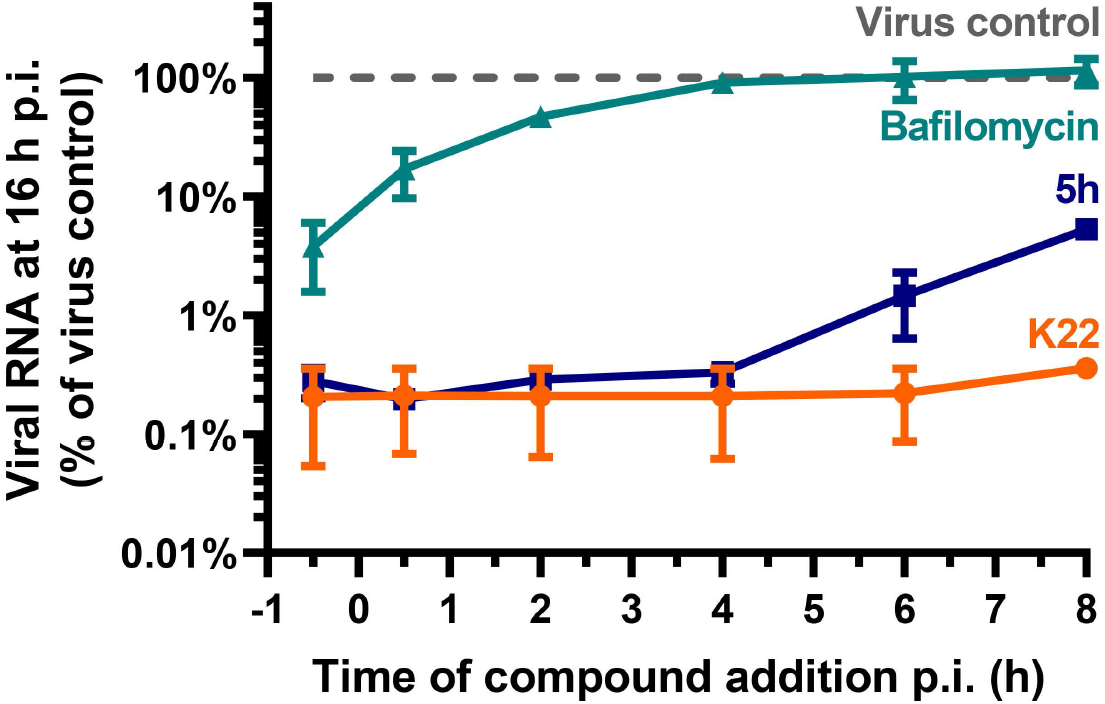
5h acts post-entry at an early stage in viral RNA synthesis. Compound addition was delayed until different time points after infecting HEL cells with HCoV-229E, and viral RNA was quantified at 16 h p.i. Compound concentrations: bafilomycin 6.3 nM; **5h** and K22: 15 μM. The Y-axis shows the viral RNA copy number relative to the virus control (mean ± SEM of two independent experiments).

Next, we performed two independent virus passaging experiments to select **5h**-resistant viruses and identify the viral target. After three cell culture passages under increasing concentrations (up to 40 μM) of **5h**, HCoV-229E acquired resistance. Whole virus genome sequencing revealed that this was attributed to two substitutions in nsp15, K60R (first selection) and T66I (second selection), located in the N-terminal part of this protein. For both mutants, **5h** exhibited an antiviral EC_99_ value (= concentration producing 100-fold reduction in virus yield) of >40 μM, which is at least 14-fold higher than the EC_99_ value of 2.9 μM measured for wild-type virus (**Fig. 3A**). Both mutant viruses remained fully sensitive to GS-441524. The conclusion that **5h** targets nsp15 was corroborated by determining its activity against a reverse-engineered EndoU-deficient H250A-mutant HCoV-229E, which lacks the catalytic His250 residue in the EndoU active site.^22^ **5h** proved dramatically less active against this mutant (**Fig. 3B**), producing a maximal reduction in virus yield of 23-fold compared to 1479-fold for wild-type (WT) virus. Again, GS-441524 proved equally active against nsp15-mutant and wild-type virus. To conclude, we established **5h** as an inhibitor of nsp15, and showed that its activity is linked to residues Lys60 and Thr66 in the N-terminal part, plus His250 in the EndoU catalytic site of nsp15. This inhibition of nsp15 accords with the time-of-addition profile of **5h** (see above), showing that the compound interferes with an early stage in viral RNA synthesis.

**Figure 3.**
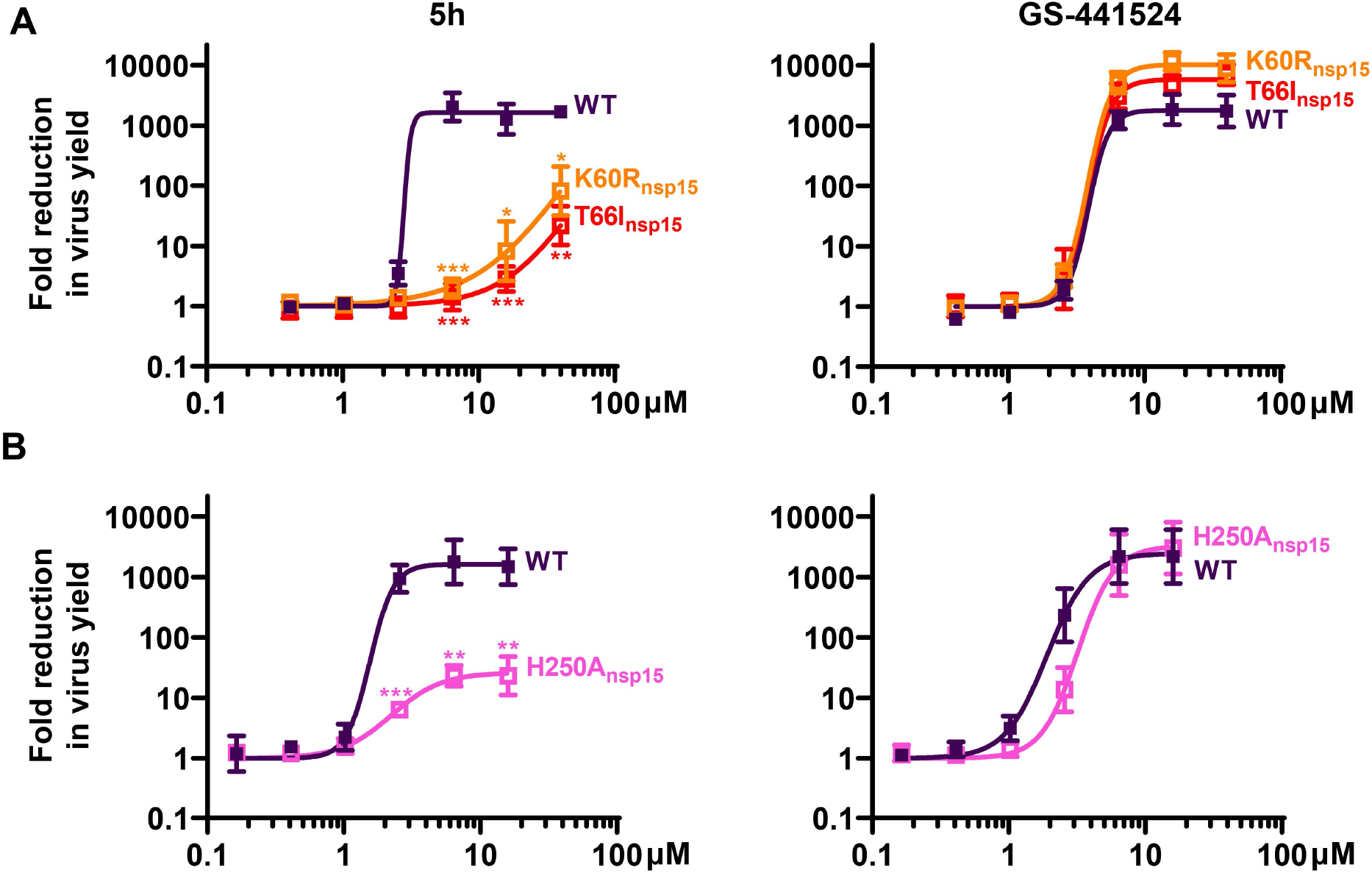
Mutations in nsp15 confer resistance of HCoV-229E to **5h** (left panels), but not to GS-441524 (right panels). The graphs show the effect of the compounds on virus yield. (A) HEL cells infected with **5h**-resistant mutants obtained by virus passaging under **5h** and carrying substitution K60R (first selection) or T66I (second selection) in nsp15. (B) 16HBE cells infected with EndoU-deficient mutant virus (H250A_nsp15_), obtained by reverse genetics.^22^ Data points are the mean ± SEM (N=3). An unpaired t-test (GraphPad Prism 8.4.3) was used to compare the mutant viruses to WT, and the resulting two-tailed p-values were adjusted for multiple comparisons using Holm-Sidak (α = 0.05). *, P < 0.05; **, P < 0.01; ***, P < 0.001.

### Binding model of 5h in hexameric nsp15 protein

To predict the possible binding site of **5h**, the compound was docked into the X-ray structure of hexameric nsp15 protein from HCoV-229E (PDB code: 4RS4). This hexameric structure formed by two trimers is the functional form of nsp15.^15^ First, a few substitutions (see Experimental section for all details) were introduced to render the protein sequence identical to that of the HCoV-229E virus used in the biological assays. By using a pocket-detection protocol implemented on the Site Finder module of Molecular Operating Environment (MOE)^19^ software, we identified a druggable binding pocket at the nsp15 dimer interface, surrounded by the catalytic residue His250 in the EndoU active site of one nsp15 monomer and residues Lys60 and Thr66 in the N-terminal domain of the other monomer (**Fig. 4A**). Next, ligand **5h** was placed inside the pocket and docked by using both MOE and GOLD softwares and the common top scoring binding mode was further analyzed.^20^ This docked result indicates that the carboxylic acid of **5h** forms hydrogen bonds with the backbone of residues Cys294 and Thr295 (**Fig. 4B**). The importance of this interaction is supported by the observation that **5u**, the **5h** analogue bearing a methyl instead of carboxylic acid group, lacks antiviral activity. Furthermore, the 1,2,3-triazole group of **5h** engages in hydrogen-bonding interactions with Thr245, explaining why the parent compounds **1** and **2** are not active against HCoV-229E. At the other side of the pocket, the aromatic ring of **5h** makes hydrophobic contacts with Val63 and Leu65. This may explain why nearby mutations K60R and T66I yield resistance to the compound, since these substitutions may negatively affect the interactions of Val63 and Leu65, or disturb the conformation of the loop flanked by both residues.

**Figure 4.**
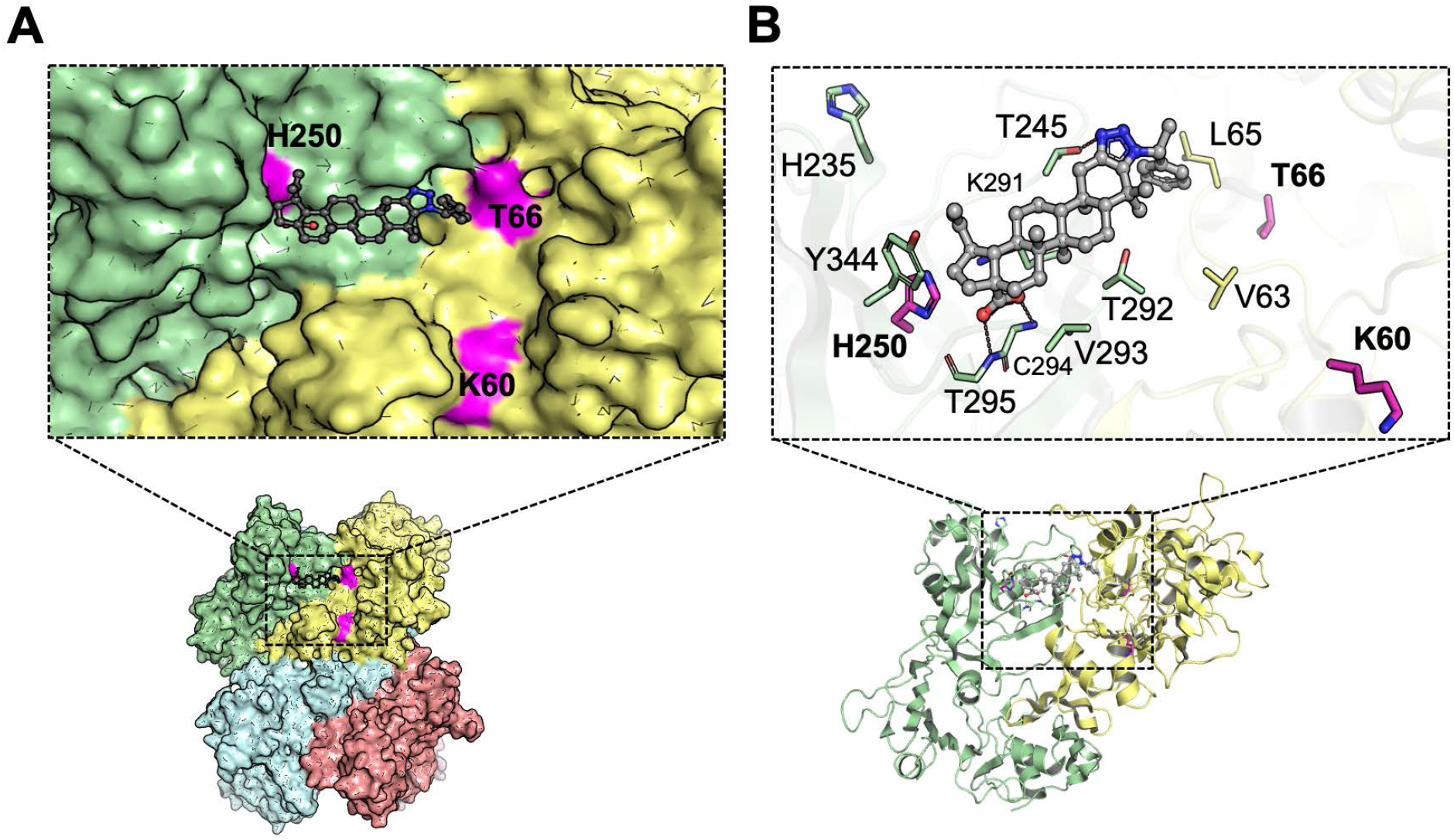
Binding mode of **5h** in HCoV-229E nsp15 hexameric protein (PDB 4RS4), as predicted by docking. (A) The hydrophobic pocket lies adjacent to the EndoU catalytic centre (catalytic triad consisting of His235, His250 and Lys291) and at an nsp15 dimer interface (monomers depicted in differently coloured surface). The pocket is surrounded by His250, Lys60 and Thr66, explaining why **5h** is inactive against HCoV-229E viruses carrying mutations at these sites. (B) **5h** occupies the pocket by making hydrophobic interactions with Val293 and side chain fragments of Lys291 and Thr292. The molecule further engages in hydrogen-bonding interactions with Cys294 and Thr295 via the carboxylic acid and with Thr245 via the 1,2,3-triazole. Additional hydrophobic interactions with Val63, Leu65 and Thr292 are made via the aromatic ring-substituted 1,2,3-triazole moiety.

Analysis of the nsp15 sequence similarity between HCoV-229E and SARS-CoV-2 showed that a few residues in the pocket are not conserved. This explains why docking **5h** into the SARS-CoV-2 nsp15 hexamer was unable to identify a similar pose within the top solutions. The largest influence seems attributed to the residue at position 244/245, since substituting Thr245 (present in HCoV-229E) by Gln244 (the corresponding residue in SARS-CoV-2) abrogates a hydrogen-bond interaction with the 1,2,3-triazole group of **5h** (**Fig. 5**). Additionally, the loop between Val/Ile63 and Leu/Pro65 has a slightly different orientation in these two CoV nsp15 proteins. Both factors may explain why **5h** is active against HCoV-229E, but not SARS-CoV-2. Still, most of the pocket residues are conserved, underscoring the relevance of this nsp15 interface pocket for drug design. The role of this protein region in forming inter-monomer interactions is evident from reports that nsp15 exists as a monomer when key interactions in this region (i.e. Arg61-Glu266 in SARS-CoV nsp15 and Tyr58-Glu263 in MERS-CoV nsp15) are eliminated by mutation.^19, 47^ When nsp15 is unable to hexamerize, the EndoU catalytic site undergoes important structural changes that abolish RNA binding and enzymatic activity.^48^

**Figure 5.**
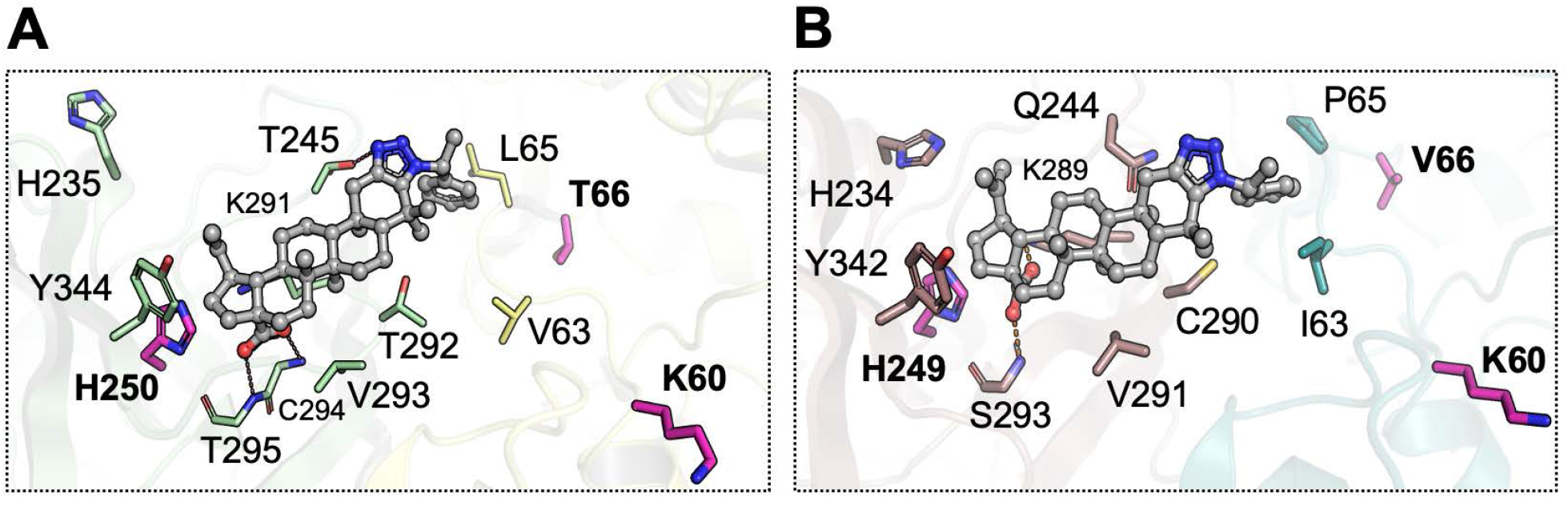
Comparison of the hydrophobic pocket, occupied by **5h**, in the nsp15 proteins of HCoV-229E (left; PDB 4RS4) and SARS-CoV-2 (right; PDB 7K1O). The carboxylic acid of **5h** forms hydrogen bonds with both nsp15 binding pockets. On the other hand, the 1,2,3-triazole group engages hydrogen-bonding interactions with the HCoV-229E nsp15 protein but is incompatible with the SARS-CoV-2 pocket.

To conclude, the nsp15 binding mode of **5h**, predicted by docking, nicely accords with the biological findings. Namely, the binding model rationalizes the requirement of the 1,2,3-triazolo function and carboxylate substituent; **5h** resistance of the nsp15-K60R, -T66I and -H250A mutant viruses; and lack of activity against SARS-CoV-2.

## CONCLUSIONS

To conclude, we present the first prototype of CoV nsp15 inhibitors, having a 1,2,3-triazolo fused betulonic acid structure. The SAR analysis, resistance data and docking model provide strong evidence that lead molecule **5h** binds to an inter-monomer nsp15 interface lying adjacent to the EndoU catalytic core. This provides an excellent basis to modify the substituents or betulonic acid scaffold, and expand the activity spectrum beyond HCoV-229E. Since **5h** appears to interact with the catalytic His250 residue in the EndoU domain of nsp15, the molecule plausibly interferes with the role of EndoU in regulating viral dsRNA synthesis.^24^ To complement the findings in this report, obtained in non-immune cells, we are currently evaluating the antiviral and immunomodulatory effects of **5h** in HCoV-229E-infected human macrophages. This may validate the intriguing concept that nsp15 inhibition could have a dual outcome, by inhibiting CoV replication and promoting host cell antiviral immunity.

## EXPERIMENTAL SECTION

### Chemistry

^1^H and ^13^C NMR spectra were measured on commercial instruments (Bruker Avance 300 MHz, Bruker AMX 400 MHz and 600 MHz). Chemical shifts (δ) are reported in parts per million (ppm) referenced to tetramethylsilane (^1^H), or the internal (NMR) solvent signal (^13^C). Melting points were determined using a Reichert Thermovar apparatus. For column chromatography, 70-230 mesh silica 60 (E. M. Merck) was used as the stationary phase. Chemicals received from commercial sources were used without further purification. Reaction dry solvents (toluene, DMF, THF) were used as received from commercial sources. TLC was carried out on Kieselgel 60 F254 plates (Merck).

Exact mass was acquired on a quadrupole orthogonal acceleration time-of-flight mass spectrometer (Synapt G2 HDMS, Waters, Milford, MA). Samples were infused at 3 μL/min and spectra were obtained in positive (or: negative) ionization mode with a resolution of 15000 (FWHM) using leucine enkephalin as lock mass.

All Liquid Chromatography (LC-MS) data were acquired on Agilent 1100 HPLC with quaternary pump, autosampler, UV*-*DAD detector and a thermostatic column (25 °C) module coupled to Agilent 6110 single-quadrupole electrospray ionization mass spectrometry (capillary voltage = +3500V or −3000V), detector was set to 210 nm for detection. Injection volume was 10 or 20 μL of a dilution of 100 μg/mL (sample in mobile phase). The column was a Grace Prevail RP-C18 3μm 150mm × 2.1mm. Data collection and analysis was done with Agilent LC/MSD Chemstation software. All tested compounds showed a purity >95.0%.

#### 3-Oxo-lup-20(29)-en-28-oic acid (betulonic acid, 2)

To a solution of betulin (50.0 g, 113 mmol; purchased from Eburon Organics BVBA) in acetone (1500 mL, use ultra-sonic bath to dissolve) was added freshly prepared Jones reagent [Na_2_Cr_2_O_7_, (66.5 g, 226 mmol) and H_2_SO_4_ (60 mL) in water (500 mL)] during 1 h in an ice bath. The reaction mixture was allowed to warm to room temperature and stirring was continued for 6 h fellow with TLC. First, MeOH was added and then water to the reaction mixture. Precipitate was filtered off and washed with water (500 mL). The crude product was dried in a vacuum oven, dissolved to Et_2_O (600 mL) and washed with water (300 mL), 7.5 % hydrochloric acid (200 mL), water (200 mL), saturated aqueous NaHCO_3_ solution (200 mL) and water (200 mL). The crude reaction mixture was purified by column chromatography (silica gel) whereas eluent was used a mixture of heptane and ethyl acetate 70:30 to afford betulonic acid 23 g 45% yield. Spectroscopic data for betulonic acid was consistent with previously reported data for this compound.^25^

#### 3-Oxo-lupan-28-oic acid(dihydro-betulonic acid)

Betulonic acid (180 mg, 0.396 mmol, 1.0 equiv.) was dissolved in a mixture of MeOH/THF (2/6 mL). 10% Pd(OH)_2_ (30 mg) was added under N_2_ atmosphere. This atmosphere was replaced by H_2_ atmosphere. The reaction was stirred under H_2_ atmosphere for 78 h, then filtered through celite and washed with CHCl_3_ to afford a white solid. The residue obtained was purified by silica gel column chromatography using 100% chloroform as eluent to afford dihydro-betulonic acid (quantitative yield) as a white amorphous powder. The spectra proved the identity of the compound by comparing the data with the literature.^26^

#### General Procedure

To a dried screw-capped reaction tube equipped with a magnetic stirring bar was added betulonic acid, amine, 4-nitrophenyl azide, 4 Å molecular sieves (50 mg). The mixture was dissolved in the proper solvent (toluene, DMF) and stirred at 100 °C for 12-72 hours. The reaction was monitored using TLC with the plate first developed with DCM then different ratios petroleum ether/ethyl acetate 7:3, 6:4 were used depending on the substrates. For visualization of TLC plates was used 5% H_2_SO_4_ in ethanol, for more sensitive detection was used cerium-ammonium-molybdate after heating to 150–200 °C. The crude reaction mixture was then directly purified by column chromatography (silica gel), at first with CH_2_Cl_2_ as an eluent to remove all 4-nitroaniline formed during the reaction, followed by using a mixture of petroleum ether and ethyl acetate as eluent to afford the betulonic acid 1,2,3-triazole derivatives.

#### 1’-(3,4,5-Trimethoxybenzyl)-1*H*’-lup-2-eno-[2,3-*d*]-[1,2,3]-triazole-28-oic acid (5a)

Betulonic acid (100 mg, 1 equiv., 0.220 mmol), 3,4,5-trimethoxybenzylamine (56 mg, 1.3 equiv., 0.286 mmol), 4-nitrophenyl azide (36 mg, 1 equiv., 0.220 mmol), 4 Å molecular sieves (50 mg) and toluene (0.5 mL). Reaction time is overnight. The product was purified by flash column chromatography (first washed with CH_2_Cl_2_ followed by petroleum ether : EtOAc 9:1 → 6:4) affording **5a** (122 mg, 84% yield) as off white crystals. m.p. 152 °C. ^1^**H NMR** (400 MHz, CDCl_3_) δ 6.25 (s, 2H), 5.57 (s, 2H), 4.76 (s, 1H), 4.64 (s, 1H), 3.81 (s, 3H), 3.77 (s, 6H), 3.04 (m, 1H), 2.96 (d, *J* = 15.4 Hz, 1H), 2.32 – 2.15 (m, 3H), 2.05 – 1.94 (m, 2H), 1.77 – 1.64 (m, 5H), 1.59 – 1.40 (m, 11H), 1.20 (s, 7H), 1.03 (s, 3H), 1.00 (s, 3H), 0.97 (s, 3H), 0.77 (s, 3H). ^13^**C NMR** (101 MHz, CDCl_3_) δ 180.9, 153.5, 150.2, 141.9, 138.0, 137.4, 132.1, 109.8, 103.5, 60.8, 56.3, 56.1, 54.6, 52.8, 49.3, 49.1, 46.8, 42.4, 40.5, 38.9, 38.5, 37.0, 33.7, 33.3, 32.0, 30.5, 29.8, 29.7, 28.7, 25.5, 21.3, 21.3, 19.4, 18.9, 16.0, 15.7, 14.6. **HRMS** (ESI^+^): m/z calculated for C_40_H_57_N_3_O_5_H [M+H]^+^: 660.4370, found 660.4384.

#### 1’-(3,5-Dimethoxybenzyl)-1*H*’-lup-2-eno-[2,3-*d*]-[1,2,3]-triazole-28-oic acid (5b)

Betulonic acid (100 mg, 1 equiv., 0.220 mmol), 3,5-dimethoxybenzylamine (48 mg, 1.3 equiv., 0.286 mmol), 4-nitrophenyl azide (36 mg, 1 equiv., 0.220 mmol), 4 Å molecular sieves (50 mg) and toluene (0.5 mL). Reaction time is overnight. The product was purified by flash column chromatography (first washed with CH_2_Cl_2_ followed by petroleum ether : EtOAc 9:1 → 6:4) affording **5b** (126 mg, 92% yield) as off white crystals. m.p. 159 °C. ^1^**H NMR** (400 MHz, CDCl_3_) δ 6.34 (s, 1H), 6.16 (s, 2H), 5.57 (s, 2H), 4.76 (s, 1H), 4.64 (s, 1H), 3.71 (s, 6H), 3.08 – 2.99 (m, 1H), 2.95 (d, *J* = 15.4 Hz, 1H), 2.32 – 2.14 (m, 3H), 2.05 – 1.94 (m, 2H), 1.79 – 1.62 (m, 5H), 1.60 – 1.33 (m, 11H), 1.29 – 1.10 (m, 7H), 1.03 (s, 3H), 1.00 (s, 3H), 0.98 (s, 3H), 0.77 (s, 3H). ^13^**C NMR** (101 MHz, CDCl_3_) δ 181.6, 161.1, 150.2, 141.8, 138.9, 138.0, 109.8, 104.4, 99.6, 56.4, 55.3, 54.5, 52.8, 49.2, 49.2, 46.9, 42.4, 40.5, 38.9, 38.5, 38.3, 37.0, 33.7, 33.3, 32.0, 30.6, 29.8, 28.7, 25.5, 21.3, 21.2, 19.4, 18.9, 16.0, 15.7, 14.6. **HRMS** (ESI^+^): m/z calculated for C_39_H_55_N_3_O_4_H [M+H]^+^: 630.4265, found 630.4274.

#### 1’-(Pyridin-4-ylmethyl)-1*H*’-lup-2-eno-[2,3-*d*]-[1,2,3]-triazole-28-oic acid (5c)

Betulonic acid (100 mg, 1 equiv., 0.220 mmol), 4-(aminomethyl)pyridine (31 mg, 1.3 equiv., 0.286 mmol), 4-nitrophenyl azide (36 mg, 1 equiv., 0.220 mmol), 4 Å molecular sieves (50 mg) and toluene (0.5 mL). Reaction time is overnight. The product was purified by flash column chromatography (first washed with CH_2_Cl_2_ followed by petroleum ether : EtOAc 9:1 → 6:4) affording **5c** (97 mg, 78% yield) as off white crystals. m.p 172 °C. ^1^**H NMR** (400 MHz, CDCl_3_) δ 8.56 (d, *J* = 5.6 Hz, 2H), 6.93 (d, *J* = 5.6 Hz, 2H), 5.66 (s, 2H), 4.76 (s, 1H), 4.64 (s, 1H), 3.06 (m, 1H), 2.97 (d, *J* = 15.4 Hz, 1H), 2.36 – 2.15 (m, 3H), 2.06 – 1.95 (m, 2H), 1.83 – 1.64 (m, 5H), 1.60 – 1.37 (m, 11H), 1.32 – 1.21 (m, 7H), 1.16 (s, 3H), 1.01 (m, 3H), 1.00 (m, 3H), 0.77 (s, 3H). ^13^**C NMR** (101 MHz, CDCl_3_) δ 180.6, 150.4, 149.8, 146.1, 142.3, 138.3, 121.2, 109.7, 56.3, 54.4, 51.6, 49.2, 49.2, 46.9, 42.4, 40.5, 39.0, 38.4, 38.2, 37.0, 33.6, 33.2, 32.1, 30.6, 29.8, 29.6, 28.8, 25.4, 21.5, 21.4, 19.4, 18.8, 16.0, 15.7, 14.6. **HRMS** (ESI^+^): m/z calculated for C_36_H_50_N_4_O_2_H [M+H]^+^: 571.4006, found 571.4013.

#### 1’-(4-Methylbenzyl)-1*H*’-lup-2-eno-[2,3-*d*]-[1,2,3]-triazole-28-oic acid (5d)

Betulonic acid (100 mg, 1 equiv., 0.220 mmol), 4-methylbenzylamine (35 mg, 1.3 equiv., 0.286 mmol), 4-nitrophenyl azide (36 mg, 1 equiv., 0.220 mmol), 4 Å molecular sieves (50 mg) and toluene (0.5 mL). Reaction time is overnight. The product was purified by flash column chromatography (first washed with CH_2_Cl_2_ followed by petroleum ether : EtOAc 9:1 → 6:4) affording **5d** (115 mg, 90% yield) as off white crystals. m.p. 310 °C. ^1^**H NMR** (600 MHz, CDCl_3_) δ 7.09 (d, *J* = 7.9 Hz, 2H), 6.92 (d, *J* = 7.9 Hz, 2H), 5.59 (s, br, 2H), 4.75 (s, 1H), 4.63 (s, 1H), 3.03 (m, 1H), 2.95 (d, *J* = 15.4 Hz, 1H), 2.32 – 2.23 (m, 5H), 2.17 (d, *J* = 15.4 Hz, 1H), 2.05 – 1.94 (m, 2H), 1.80 – 1.62 (m, 5H), 1.50 (m 11H), 1.22 – 1.09 (m, 7H), 1.03 (s, 3H), 0.99 (s, 3H), 0.97 (s, 3H), 0.77 (s, 3H). ^13^**C NMR** (101 MHz, CDCl_3_) δ 181.8, 150.2, 141.7, 138.0, 137.4, 133.3, 129.3, 126.3, 109.8, 56.4, 54.5, 52.6, 49.2, 49.1, 46.8, 42.4, 40.8, 40.5, 38.9, 38.4, 38.2, 37.0, 33.7, 33.3, 32.0, 30.5, 29.7, 28.7, 28.4, 25.4, 23.8, 21.3, 21.3, 21.0, 20.8, 19.4, 18.9, 17.5, 17.2, 16.0, 15.6, 14.6. **HRMS** (ESI^+^): m/z calculated for C_38_H_53_N_3_O_2_H [M+H]^+^: 584.4210, found 584.4217.

#### 1’-(4-Fluorobenzyl)-1*H*’-lup-2-eno-[2,3-*d*]-[1,2,3]-triazole-28-oic acid (5e)

Betulonic acid (100 mg, 1 equiv., 0.220 mmol), 4-fluorobenzylamine (36 mg, 1.3 equiv., 0.286 mmol), 4-nitrophenyl azide (36 mg, 1 equiv., 0.220 mmol), 4 Å molecular sieves (50 mg) and toluene (0.5 mL). Reaction time is overnight. The product was purified by flash column chromatography (first washed with CH_2_Cl_2_ followed by petroleum ether : EtOAc 9:1 → 6:4) affording **5e** (110 mg, 85% yield) as off white crystals. m.p. 309 °C. ^1^**H NMR** (400 MHz, CDCl_3_) δ 7.00 (m, 4H), 5.60 (s, 2H), 4.76 (s, 1H), 4.64 (s, 1H), 3.08 – 2.99 (m, 1H), 2.95 (d, *J* = 15.3 Hz, 1H), 2.33 – 2.14 (m, 3H), 2.06 – 1.93 (m, 2H), 1.82 – 1.62 (m, 5H), 1.61 – 1.31 (m, 11H), 1.30 – 1.10 (m, 7H), 1.08 (s, 3H), 1.02 (s, 3H), 1.00 (s, 3H), 0.77 (s, 3H). ^13^**C NMR** (101 MHz, CDCl_3_) δ 181.1, 163.5, 161.0, 150.2, 142.0, 137.9, 132.2, 132.2, 128.2, 128.1, 115.8, 115.5, 109.8, 56.3, 54.5, 52.1, 49.2, 49.2, 46.9, 42.4, 40.5, 38.9, 38.5, 38.3, 37.0, 33.7, 33.3, 32.0, 30.5, 29.8, 28.7, 25.4, 21.3, 21.3, 19.4, 18.8, 16.0, 15.7, 14.6. **HRMS** (ESI^+^): m/z calculated for C_37_H_50_FN_3_O_2_H [M+H]^+^: 588.3959, found 588.3969.

#### 1’-(4-Trifluoromethylbenzyl)-1*H*’-lup-2-eno-[2,3-*d*]-[1,2,3]-triazole-28-oic acid (5f)

Betulonic acid (100 mg, 1 equiv., 0.220 mmol), 4-(trifluoromethyl)benzylamine (50 mg, 1.3 equiv., 0.286 mmol), 4-nitrophenyl azide (36 mg, 1 equiv., 0.220 mmol), 4 Å molecular sieves (50 mg) and toluene (0.5 mL). Reaction time is overnight. The product was purified by flash column chromatography (first washed with CH_2_Cl_2_ followed by petroleum ether : EtOAc 9:1 → 6:4) affording **5f** (112 mg, 80% yield) as off white crystals. m.p. 315 °C. ^1^**H NMR** (400 MHz, CDCl_3_) δ 7.56 (d, *J* = 8.0 Hz, 2H), 7.11 (d, *J* = 8.0 Hz, 2H), 5.69 (s, 2H), 4.76 (s, 1H), 4.64 (s, 1H), 3.02 (m, 2H), 2.34 – 2.16 (m, 3H), 2.07 – 1.94 (m, 2H), 1.82 – 1.64 (m, 5H), 1.59 – 1.38 (m, 11H), 1.18 (m, 7H), 1.02 (s, 3H), 1.00 (s, 3H), 0.97 (s, 3H), 0.78 (s, 3H). ^13^**C NMR** (101 MHz, CDCl_3_) δ 181.4, 150.2, 142.1, 140.5, 138.2, 126.6, 125.7, 125.7, 109.8, 56.4, 54.5, 52.2, 49.3, 49.1, 46.9, 42.4, 40.8, 40.5, 38.9, 38.5, 38.2, 37.0, 33.8, 33.6, 33.3, 32.0, 30.5, 29.8, 29.7, 28.8, 28.4, 25.4, 23.8, 21.4, 21.3, 20.8, 20.5, 19.4, 18.8, 17.5, 17.3, 16.1, 15.7, 14.6, 7.9. **HRMS** (ESI^+^): m/z calculated for C_38_H_50_F_3_N_3_O_2_H [M+H]^+^: 638.3927, found 638.3939.

#### 1’-(4-Dimethylaminobenzyl)-1*H*’-lup-2-eno-[2,3-*d*]-[1,2,3]-triazole-28-oic acid (5g)

Betulonic acid (100 mg, 1 equiv., 0.220 mmol), 4-(dimethylamino)benzylamine (43 mg, 1.3 equiv., 0.286 mmol), 4-nitrophenyl azide (36 mg, 1 equiv., 0.220 mmol), 4 Å molecular sieves (50 mg) and toluene (0.5 mL). Reaction time is overnight. The product was purified by flash column chromatography (first washed with CH_2_Cl_2_ followed by petroleum ether : EtOAc 9:1 → 6:4) affording **5g** (112 mg, 80% yield) as off white crystals. m.p. 190 °C. ^1^**H NMR** (400 MHz, CDCl_3_) δ 6.97 (d, *J* = 8.3 Hz, 2H), 6.64 (d, *J* = 8.3 Hz, 2H), 5.52 (s, 2H), 4.76 (s, 1H), 4.64 (s, 1H), 3.10 – 2.84 (m, 9H), 2.33 – 2.11 (m, 3H), 1.99 (m, 2H), 1.81 – 1.62 (m, 5H), 1.56 – 1.34 (m, 11H), 1.25 – 1.15 (m, 7H), 1.06 (s, 3H), 0.99 (s, 3H), 0.97 (s, 3H), 0.76 (s, 3H). ^13^**C NMR** (101 MHz, CDCl_3_) δ 181.4, 150.2, 150.1, 141.6, 137.7, 127.6, 123.9, 112.5, 109.8, 56.4, 54.6, 52.6, 49.2, 49.1, 46.8, 42.4, 40.5, 40.5, 38.9, 38.9, 38.4, 38.3, 37.0, 33.7, 33.3, 32.1, 30.5, 29.8, 28.7, 25.5, 23.8, 21.3, 21.3, 20.8, 19.4, 18.9, 16.0, 15.6, 14.6. **HRMS** (ESI^+^): m/z calculated for C_39_H_56_N_4_O_2_H [M+H]^+^: 613.4475, found 613.4480.

#### 1’-(1-Phenylethyl)-1*H*’-lup-2-eno-[2,3-*d*]-[1,2,3]-triazole-28-oic acid (5h)

Betulonic acid (100 mg, 1 equiv., 0.220 mmol), (S)-(−)-α-methylbenzylamine (35 mg, 1.3 equiv, 0.286 mmol), 4-nitrophenyl azide (36 mg, 1 equiv., 0.220 mmol), 4 Å molecular sieves (50 mg) and toluene (0.5 mL). Reaction time is overnight. The product was purified by flash column chromatography (first washed with CH_2_Cl_2_ followed by petroleum ether : EtOAc 9:1 → 6:4) affording **5h** (107 mg, 84% yield) as off white crystals. m.p. 327 °C. ^1^**H NMR** (400 MHz, CDCl_3_) δ 7.30 – 7.21 (m, 3H), 7.13 (d, *J* = 7.2 Hz, 2H), 5.73 (m, 1H), 4.76 (s, 1H), 4.64 (s, 1H), 3.00 (m, 2H), 2.33 – 2.12 (m, 3H), 2.07 – 1.92 (m, 5H), 1.81 – 1.62 (m, 5H), 1.58 – 1.37 (m, 11H), 1.14 (m, 7H), 1.00 (s, 6H), 0.96 (s, 3H), 0.72 (s, 3H). ^13^**C NMR** (101 MHz, CDCl_3_) δ 180.6, 150.2, 141.8, 141.1, 137.6, 128.6, 127.5, 126.1, 109.8, 59.1, 56.3, 54.8, 49.3, 49.1, 46.8, 42.4, 40.5, 38.8, 38.4, 37.0, 33.6, 33.3, 32.0, 30.5, 29.7, 29.7, 28.6, 25.4, 23.3, 21.4, 21.3, 19.4, 19.0, 15.9, 15.6, 14.6. **HRMS** (ESI^+^): m/z calculated for C_38_H_53_N_3_O_2_H [M+H]^+^: 584.4210, found 584.4218.

#### 1’-(Cyclohexylmethyl)-1*H*’-lup-2-eno-[2,3-*d*]-[1,2,3]-triazole-28-oic acid (5i)

Betulonic acid (100 mg, 1 equiv., 0.220 mmol), cyclohexanemethylamine (33 mg, 1.3 equiv., 0.286 mmol), 4-nitrophenyl azide (36 mg, 1 equiv., 0.220 mmol), 4 Å molecular sieves (50 mg) and toluene (0.5 mL). Reaction time is overnight. The product was purified by flash column chromatography (first washed with CH_2_Cl_2_ followed by petroleum ether : EtOAc 9:1 → 6:4) affording **5i** (103 mg, 82% yield) as off white crystals. m.p. 338 °C. ^1^**H NMR** (600 MHz, CDCl_3_) δ 4.76 (s, 1H), 4.64 (s, 1H), 4.09 (m, 2H), 3.04 (m, 1H), 2.92 (m, 1H), 2.33 – 2.21 (m, 2H), 2.13 (d, *J* = 15.3 Hz, 1H), 2.07 – 1.95 (m, 2H), 1.79 – 1.64 (m, 10H), 1.61 – 1.34 (m, 11H), 1.32 – 1.19 (m, 10H), 1.18 – 1.08 (m, 5H), 1.00 (s, 3H), 0.99 (s, 3H), 0.77 (s, 3H). ^13^**C NMR** (101 MHz, CDCl_3_) δ 181.7, 150.2, 140.9, 137.8, 109.8, 56.4, 55.6, 54.7, 49.2, 49.2, 46.9, 42.4, 40.5, 38.8, 38.6, 38.5, 38.2, 37.0, 33.7, 33.3, 32.1, 31.0, 30.6, 29.8, 28.9, 26.3, 25.7, 25.4, 21.5, 21.3, 19.4, 18.9, 16.0, 15.7, 14.6. **HRMS** (ESI^+^): m/z calculated for C_37_H_57_N_3_O_2_H [M+H]^+^: 576.4523, found 576.4529.

#### 1’-(Benzo[*d*][1,3]dioxol-5-ylmethyl)-1*H*’-lup-2-eno-[2,3-*d*]-[1,2,3]-triazole-28-oic acid (5j)

Betulonic acid (100 mg, 1 equiv., 0.220 mmol), piperonylamine (43 mg, 1.3 equiv., 0.286 mmol), nitrophenyl azide (36 mg, 1 equiv., 0.220 mmol), 4 Å molecular sieves (50 mg) and toluene (0.5 mL). Reaction time is overnight. The product was purified by flash column chromatography (first washed with CH_2_Cl_2_ followed by petroleum ether: EtOAc 9:1 → 6:4) affording **5j** (105 mg, 78% yield) as off white crystals. m.p. 313 °C. ^1^**H NMR** (300 MHz, CDCl_3_) δ 6.72 (d, *J* = 7.9 Hz, 1H), 6.59 – 6.48 (m, 2H), 5.93 (d, *J* = 0.9 Hz, 2H), 5.53 (s, 2H), 4.76 (s, 1H), 4.64 (s, 1H), 3.10 – 2.89 (m, 2H), 2.35 – 2.11 (m, 3H), 1.99 (m, 2H), 1.83 – 1.62 (m, 5H), 1.61 – 1.33 (m, 11H), 1.33 – 1.08 (m, 7H), 1.05 (s, 3H), 1.00 (s, 3H), 0.97 (s, 3H), 0.77 (s, 3H). ^13^**C NMR** (101 MHz, CDCl_3_) δ 181.3, 150.2, 148.1, 147.2, 141.8, 137.9, 130.1, 119.9, 109.8, 108.3, 107.2, 101.1, 56.4, 54.5, 52.6, 50.8, 49.2, 49.2, 46.9, 42.4, 40.5, 38.9, 38.5, 38.3, 37.0, 33.7, 33.3, 32.0, 30.5, 29.8, 28.7, 25.4, 21.3, 21.3, 19.4, 18.9, 16.0, 15.7, 14.6. **HRMS** (ESI^+^): m/z calculated for C_38_H_51_N_3_O_4_H [M+H]^+^: 614.3952, found 614.3951.

#### 1’-(Furan-2-ylmethyl)-1*H*’-lup-2-eno-[2,3-*d*]-[1,2,3]-triazole-28-oic acid (5k)

Betulonic acid (100 mg, 1 equiv., 0.220 mmol), furfurylamine (28 mg, 1.3 equiv., 0.286 mmol), 4-nitrophenyl azide (36 mg, 1 equiv., 0.220 mmol), 4 Å molecular sieves (50 mg) and toluene (0.5 mL). Reaction time is overnight. The product was purified by flash column chromatography (first washed with CH_2_Cl_2_ followed by petroleum ether : EtOAc 9:1 → 6:4) affording **5k** (66 mg, 53% yield) as a brown crystals. m.p. 227 °C. ^1^**H NMR** (400 MHz, CDCl_3_) δ 7.35 (s, 1H), 6.34 – 6.31 (m, 1H), 6.23 (d, *J* = 3.1 Hz, 1H), 5.55 (s, 2H), 4.76 (s, 1H), 4.64 (s, 1H), 3.07 – 2.99 (m, 1H), 2.93 (d, *J* = 15.3 Hz, 1H), 2.30 – 2.11 (m, 3H), 1.99 (d, *J* = 7.5 Hz, 2H), 1.78 – 1.66 (m, 5H), 1.60 – 1.40 (m, 11H), 1.28 (t, *J* = 10.2 Hz, 7H), 1.15 (s, 3H), 1.00 (s, 3H), 0.98 (s, 3H), 0.77 (s, 3H). ^13^**C NMR** (101 MHz, CDCl_3_) δ 179.9, 150.2, 148.8, 142.5, 141.6, 137.7, 110.7, 109.9, 109.0, 56.3, 54.6, 49.3, 49.2, 46.8, 46.4, 42.4, 40.6, 39.0, 38.4, 38.3, 37.0, 33.6, 33.3, 32.0, 30.5, 29.8, 29.7, 28.6, 19.4, 18.9, 16.1, 15.7, 14.6. **HRMS** (ESI^+^): m/z calculated for C_35_H_49_N_3_O_3_H [M+H]^+^: 560.3846, found 560.3857.

#### 1’-((1*H*-Indol-3-yl)methyl)-1*H*’-lup-2-eno-[2,3-*d*]-[1,2,3]-triazole-28-oic acid (5l)

Betulonic acid (100 mg, 1 equiv., 0.220 mmol), tryptamine (46 mg, 1.3 equiv., 0.286 mmol), 4-nitrophenyl azide (36 mg, 1 equiv., 0.220 mmol), 4 Å molecular sieves (50 mg) and toluene (0.5 mL). Reaction time is overnight. The product was purified by flash column chromatography (first washed with CH_2_Cl_2_ followed by petroleum ether : EtOAc 9:1 → 6:4) affording **5l** (115 mg, 84% yield) as off white crystals. m.p. 196 °C. ^1^**H NMR** (400 MHz, DMSO) δ 10.87 (s, 1H), 7.41 (d, *J* = 7.8 Hz, 1H), 7.33 (d, *J* = 8.1 Hz, 1H), 7.10 – 7.02 (m, 2H), 6.96 (m, 1H), 4.72 (s, 1H), 4.72 (s, 1H), 4.59 (s, 1H), 4.54 (s, 2H), 3.17 (s, 1H), 3.03 – 2.92 (m, 1H), 2.70 (d, *J* = 15.2 Hz, 1H), 2.29 (m, 1H), 2.14 – 2.06 (m, 2H), 1.81 (d, *J* = 6.9 Hz, 2H), 1.63 (d, *J* = 29.9 Hz, 5H), 1.57 – 1.24 (m, 11H), 1.25 – 1.01 (m, 7H), 0.96 (s, 3H), 0.93 (s, 3H), 0.90 (s, 3H), 0.63 (s, 3H). ^13^**C NMR** (101 MHz, DMSO) δ 177.7, 150.7, 140.3, 137.8, 136.5, 127.4, 123.7, 121.4, 118.9, 118.3, 111.8, 110.7, 110.1, 55.9, 54.4, 50.1, 49.0, 48.9, 47.0, 42.5, 38.8, 38.1, 36.7, 33.5, 33.3, 32.0, 31.1, 30.5, 29.7, 28.6, 26.9, 25.5, 21.4, 21.1, 19.4, 18.8, 16.1, 15.7, 14.8. **HRMS** (ESI^+^): m/z calculated for C_40_H_54_N_4_O_2_H [M+H]^+^: 623.4319, found 623.4317.

#### 1’-(2-(1*H*-Indol-3-yl)ethyl)-1*H*’-lup-2-eno-[2,3-*d*]-[1,2,3]-triazole-28-oic acid (5m)

Betulonic acid (100 mg, 1 equiv., 0.220 mmol), 5-methoxytryptamine (54 mg, 1.3 equiv., 0.286 mmol), 4-nitrophenyl azide (36 mg, 1 equiv., 0.220 mmol), 4 Å molecular sieves (50 mg) and toluene (0.5 mL). Reaction time is overnight. The product was purified by flash column chromatography (first washed with CH_2_Cl_2_ followed by petroleum ether : EtOAc 9:1 → 6:4) affording **5m** (126 mg, 88% yield) as off white crystals. m.p. 240 °C. ^1^**H NMR** (400 MHz, DMSO) δ 10.69 (s, 1H), 7.19 (d, *J* = 8.7 Hz, 1H), 7.05 (s, 1H), 6.76 (s, 1H), 6.67 (d, *J* = 8.7 Hz, 1H), 4.72 (s, 1H), 4.59 (s, 1H), 4.57 – 4.45 (m, 2H), 3.71 (s, 2H), 3.03 – 2.92 (m, 1H), 2.69 (d, *J* = 15.3 Hz, 1H), 2.28 (m, 1H), 2.09 (m, 2H), 1.81 (d, *J* = 6.9 Hz, 2H), 1.73 – 1.55 (m, 5H), 1.32 (m, 11H), 1.19 – 1.02 (m, 7H), 0.96 (s, 3H), 0.89 (s, 3H), 0.86 (s, 3H), 0.58 (s, 3H). ^13^**C NMR** (101 MHz, DMSO) δ 177.7, 153.5, 150.7, 140.3, 137.9, 131.5, 127.8, 124.3, 112.4, 111.7, 110.7, 110.1, 99.9, 55.9, 55.5, 54.4, 50.2, 48.9, 47.0, 42.5, 38.8, 38.1, 36.7, 33.5, 33.3, 32.0, 30.5, 29.7, 28.5, 26.9, 25.5, 22.5, 21.4, 20.9, 19.4, 18.8, 16.0, 15.7, 14.8. **HRMS** (ESI^+^): m/z calculated for C_41_H_56_N_4_O_3_H [M+H]^+^: 653.4424, found 653.4418.

#### 1’-((1*H*-Indol-4-yl)methyl)-1*H*’-lup-2-eno-[2,3-*d*]-[1,2,3]-triazole-28-oic acid (5n)

Betulonic acid (100 mg, 1 equiv., 0.220 mmol), 4-(aminomethyl) indole (42 mg, 1.3 equiv., 0.286 mmol), 4-nitrophenyl azide (36 mg, 1 equiv., 0.220 mmol), 4 Å molecular sieves (50 mg) and toluene (0.5 mL). Reaction time is overnight. The product was purified by flash column chromatography (first washed with CH_2_Cl_2_ followed by petroleum ether : EtOAc 9:1 → 6:4) affording **5n** (98 mg, 73% yield) as off white crystals. m.p. 260 °C. ^1^**H NMR** (400 MHz, CDCl_3_) δ 8.32 (s, 1H), 7.31 (d, *J* = 8.2 Hz, 1H), 7.22 (m, 1H), 7.07 (m, 1H), 6.52 – 6.40 (m, 2H), 5.93 (m, 2H), 4.76 (s, 1H), 4.65 (s, 1H), 3.07 – 2.93 (m, 2H), 2.31 – 2.16 (m, 3H), 2.05 – 1.93 (m, 2H), 1.77 – 1.64 (m, 5H), 1.59 – 1.35 (m, 11H), 1.34 – 1.15 (m, 7H), 1.12 (d, *J* = 14.6 Hz, 3H), 0.99 (s, 3H), 0.96 (s, 3H), 0.78 (s, 3H). ^13^**C NMR** (101 MHz, CDCl_3_) δ 180.4, 150.2, 141.9, 138.3, 135.7, 128.2, 125.2, 124.5, 122.1, 117.5, 110.6, 109.8, 100.0, 56.3, 54.6, 51.4, 49.2, 49.2, 46.9, 42.4, 40.5, 38.9, 37.0, 33.8, 33.3, 32.0, 30.5, 29.7, 29.7, 28.4, 25.5, 22.7, 21.3, 20.9, 19.4, 18.9, 16.0, 15.6, 14.6. **HRMS** (ESI^+^): m/z calculated for C_39_H_52_N_4_O_2_H [M+H]^+^: 609.4162, found 609.4174.

#### 1’-(2-Hydroxyethyl)-1*H*’-lup-2-eno-[2,3-*d*]-[1,2,3]-triazole-28-oic acid (5o)

Betulonic acid (100 mg, 1 equiv., 0.220 mmol), ethanolamine (17 mg, 1.3 equiv., 0.286 mmol), 4-nitrophenyl azide (36 mg, 1 equiv., 0.220 mmol), 4 Å molecular sieves (50 mg) and toluene (0.5 mL). Reaction time is overnight. The product was purified by flash column chromatography (first washed with CH_2_Cl_2_ followed by CH_2_Cl_2_ : MeOH 95:5) affording **5o** (70 mg, 62% yield) as off white crystals. m.p. 236 °C. ^1^**H NMR** (400 MHz, DMSO) δ 4.72 (s, 1H), 4.59 (s, 1H), 4.36 (m, 2H), 3.86 (m, 2H), 2.98 (m, 1H), 2.70 (d, *J* = 15.2 Hz, 1H), 2.30 (m, 1H), 2.11 (m, 2H), 1.82 (d, *J* = 6.9 Hz, 2H), 1.64 (s,br, 5H), 1.58 – 1.33 (m, 11H), 1.33 – 1.22 (m, 7H), 1.17 (s, 3H), 0.98 (s, 3H), 0.93 (s, 3H), 0.71 (s, 3H). ^13^**C NMR** (101 MHz, DMSO) δ 177.7, 150.7, 140.2, 138.0, 110.1, 79.6, 60.7, 55.9, 54.5, 51.4, 48.9, 47.0, 42.5, 38.9, 38.1, 36.7, 33.6, 33.3, 32.0, 30.5, 29.8, 28.7, 25.5, 21.4, 19.4, 18.8, 16.2, 15.8, 14.8. **HRMS** (ESI^+^): m/z calculated for C_32_H_49_N_3_O_3_H [M+H]^+^: 524.3846, found 524.3853.

#### 1’-Heptyl-1*H*’-lup-2-eno-[2,3-*d*]-[1,2,3]-triazole-28-oic acid (5p)

Betulonic acid (100 mg, 1 equiv., 0.220 mmol), 1-heptylamine (33 mg, 1.3 equiv, 0.286 mmol), 4-nitrophenyl azide (36 mg, 1 equiv, 0.220 mmol), 4 Å molecular sieves (50 mg) and toluene (0.5 mL). Reaction time is overnight. The product was purified by flash column chromatography (first washed with CH_2_Cl_2_ followed by petroleum ether : EtOAc 9:1 → 6:4) affording **5p** (72 mg, 57% yield) as off white crystals. m.p. 273 °C. ^1^**H NMR** (600 MHz, CDCl_3_) δ 4.76 (s, 1H), 4.64 (s, 1H), 4.33 – 4.23 (m, 2H), 3.04 (m, 1H), 2.91 (d, *J* = 15.3 Hz, 1H), 2.33 – 2.22 (m, 2H), 2.13 (d, *J* = 15.3 Hz, 1H), 2.00 – 1.96 (m, 2H), 1.79 – 1.64 (m, 5H), 1.61 – 1.35 (m, 13H), 1.35 – 1.19 (m, 14H), 1.17 (s, 3H), 1.00 (s, 3H), 0.99 (s, 3H), 0.88 (m, 3H), 0.77 (s, 3H). ^13^**C NMR** (101 MHz, CDCl_3_) δ 181.7, 150.2, 141.0, 137.3, 109.8, 56.4, 54.6, 49.6, 49.2, 49.2, 46.9, 42.4, 40.5, 38.9, 38.5, 38.2, 37.0, 33.6, 33.3, 32.1, 31.6, 30.8, 30.6, 29.8, 29.7, 28.8, 28.6, 26.9, 25.4, 22.5, 21.3, 19.4, 18.9, 16.0, 15.7, 14.6, 14.0. **HRMS** (ESI^+^): m/z calculated for C_37_H_59_N_3_O_2_H [M+H]^+^: 578.4679, found 578.4687.

#### 1*H*’-Lup-2-eno-[2,3-*d*]-[1,2,3]-triazole-28-oic acid (5q)

Betulonic acid (100 mg, 1 equiv., 0.220 mmol), ammonium acetate (85 mg, 5 equiv., 1.100 mmol), 4-nitrophenyl azide (51 mg, 1.4 equiv., 0.308 mmol), 4 Å molecular sieves (50 mg) and DMF (0.8 mL). Reaction time is overnight. The product was purified by flash column chromatography (first washed with CH_2_Cl_2_ followed by CH_2_Cl_2_ : MeOH 95:5) affording **5q** (90 mg, 86% yield) as off white crystals. m.p. 158 °C. Spectroscopic data for compound **5q** was consistent with previously reported data for this compound.^27^ ^1^**H NMR** (400 MHz, CDCl_3_) δ 4.77 (s, 1H), 4.64 (s, 1H), 3.03 (d, *J* = 10.1 Hz, 1H), 2.90 (d, *J* = 15.5 Hz, 1H), 2.37 – 2.22 (m, 2H), 2.12 (d, *J* = 15.5 Hz, 1H), 2.00 (dd, *J* = 19.9, 11.7 Hz, 2H), 1.83 – 1.66 (m, 5H), 1.57 (dd, *J* = 38.5, 24.6 Hz, 12H), 1.36 – 1.24 (m, 7H), 1.21 (d, *J* = 7.0 Hz, 4H), 1.01 (s, 3H), 1.00 (s, 3H) 0.78 (s, 3H**). ^13^C NMR** (101 MHz, CDCl_3_) δ 181.0, 150.3, 150.1, 140.5, 109.8, 56.3, 53.4, 49.2, 49.0, 46.9, 42.5, 40.7, 39.0, 38.4, 37.3, 37.0, 33.3, 33.3, 32.1, 31.0, 30.6, 29.8, 25.5, 23.7, 21.4, 19.4, 19.1, 16.2, 15.6, 14.6. **HRMS** (ESI^+^): m/z calculated for C_30_H_45_N_3_O_2_H [M+H]^+^: 480.3584, found 480.3585.

#### 1’-Benzyl-1*H*’-lup-2-eno-[2,3-*d*]-[1,2,3]-triazole-28-oic acid (5r)

Betulonic acid (100 mg, 1 equiv., 0.220 mmol), benzylamine (31 mg, 1.3 equiv., 0.286 mmol), 4-nitrophenyl azide (36 mg, 1 equiv., 0.220 mmol), 4 Å molecular sieves (50 mg) and toluene (0.5 mL). Reaction time is overnight. The product was purified by flash column chromatography (first washed with CH_2_Cl_2_ followed by petroleum ether : EtOAc 9:1 → 6:4) affording **5r** (115 mg, 92% yield) as off white crystals. m.p. 290 °C. ^1^**H NMR** (400 MHz, CDCl_3_) δ 7.33 – 7.24 (m, 3H), 7.02 (d, *J* = 7.2 Hz, 2H), 5.64 (s, 2H), 4.76 (s, 1H), 4.64 (s, 1H), 3.09 – 3.00 (m, 1H), 2.96 (d, *J* = 15.4 Hz, 1H), 2.33 – 2.13 (m, 3H), 2.06 – 1.93 (m, 2H), 1.82 – 1.63 (m, 5H), 1.61 – 1.37 (m, 11H), 1.27 – 1.15 (m, 7H), 1.04 (s, 3H), 0.99 (s, 3H), 0.97 (s, 3H), 0.77 (s, 3H). ^13^**C NMR** (101 MHz, CDCl_3_) δ 181.5, 150.2, 141.8, 138.0, 136.4, 128.7, 127.7, 126.3, 109.8, 56.4, 54.5, 52.8, 49.2, 49.1, 46.9, 42.4, 40.5, 38.9, 38.5, 38.3, 37.0, 33.7, 33.3, 32.0, 30.5, 29.7, 28.7, 25.4, 23.8, 21.3, 21.3, 19.4, 18.9, 16.0, 15.7, 14.6. **HRMS** (ESI^+^): m/z calculated for C_37_H_51_N_3_O_2_H [M+H]^+^: 570.4053, found 570.4064.

#### 1’-((*S*)-1-Phenylethyl)-1*H*’-lup-2-eno-[2,3-*d*]-[1,2,3]-triazole-28-oic acid (5s)

Betulonic acid (100 mg, 1 equiv., 0.220 mmol), (R)-(+)-α-methylbenzylamine (35 mg, 1.3 equiv., 0.286 mmol), 4-nitrophenyl azide (36 mg, 1 equiv., 0.220 mmol), 4 Å molecular sieves (50 mg) and toluene (0.5 mL). Reaction time is overnight. The product was purified by flash column chromatography (first washed with CH_2_Cl_2_ followed by petroleum ether : EtOAc 9:1 → 6:4) affording **5s** (88 mg, 69% yield) as off white crystals. m.p. 327 °C. ^1^**H NMR** (600 MHz, CDCl_3_) δ 7.30 – 7.20 (m, 3H), 7.15 – 7.10 (m, 2H), 5.73 (m, 1H), 4.75 (s, br, 1H), 4.64 (s,br, 1H), 3.02 (m, 1H), 2.95 (d, *J* = 15.4 Hz, 1H), 2.32 – 2.22 (m, 2H), 2.17 (d, *J* = 15.4 Hz, 1H), 2.05 – 1.95 (m, 5H), 1.80 – 1.64 (m, 5H), 1.64 – 1.39 (m, 11H), 1.38 – 1.25 (m, 7H), 1.00 (s, 3H), 0.99 (s, 3H), 0.96 (s, 3H) 0.72 (s, 3H). ^13^**C NMR** (101 MHz, CDCl_3_) δ 181.5, 150.2, 141.7, 141.1, 137.6, 128.6, 127.5, 126.1, 109.8, 59.1, 56.4, 54.8, 49.3, 49.1, 46.8, 42.4, 40.5, 38.8, 38.4, 38.3, 37.0, 33.6, 33.3, 32.0, 30.5, 29.7, 28.6, 25.4, 23.3, 21.4, 21.3, 19.4, 19.0, 15.9, 15.6, 14.6. **HRMS** (ESI^+^): m/z calculated for C_38_H_53_N_3_O_2_H [M+H]^+^: 584.4210, found 584.4214.

#### 1’-((S)-1-Phenylethyl)-1*H*’-lupano-[2,3-*d*]-[1,2,3]-triazole-28-oic acid (5t)

Dihydrobetulonic acid (100 mg, 1 equiv., 0.219 mmol), (S)-(−)-α-methylbenzylamine (34 mg, 1.3 equiv., 0.285 mmol), 4-nitrophenyl azide (36 mg, 1 equiv., 0.219 mmol), 4 Å molecular sieves (50 mg) and toluene (0.5 mL). Reaction time is overnight. The product was purified by flash column chromatography (first washed with CH_2_Cl_2_ followed by petroleum ether : EtOAc 9:1 → 6:4) affording **5t** (115 mg, 90% yield) as off white crystals. m.p. 173 °C. ^1^**H NMR** (400 MHz, CDCl_3_) δ 7.33 – 7.27 (m, 2H), 7.22 (m, 3H), 5.73 (m, 1H), 2.98 (d, *J* = 15.2 Hz, 1H), 2.32 – 2.22 (m, 3H), 2.17 (d, *J* = 15.3 Hz, 1H), 2.03 (m, 3H), 1.96 – 1.63 (m, 5H), 1.45 (m, 11H), 1.29 – 1.22 (m, 7H), 1.10 (s, 3H), 0.97 (m, 6H), 0.87 (s, 3H), 0.82 (s, 3H), 0.77 (s, 3H). ^13^**C NMR** (101 MHz, CDCl_3_) δ 181.5, 141.7, 141.0, 137.5, 128.6, 127.5, 126.2, 59.2, 56.8, 54.7, 49.1, 48.7, 44.1, 42.6, 40.6, 38.8, 38.3, 38.3, 37.4, 33.8, 33.4, 31.9, 29.8, 29.7, 28.8, 26.8, 23.7, 23.0, 22.7, 21.3, 21.3, 18.9, 16.2, 15.7, 14.7, 14.5. **HRMS** (ESI^+^): m/z calculated for C_38_H_55_N_3_O_2_H [M+H]^+^: 586.4366, found 586.4370.

#### 1’-((*S*)-1-Phenylethyl)-28-methyl-1*H*’-lup-2-eno-[2,3-*d*]-[1,2,3]-triazole (5u)

Lupenone (100 mg, 1 equiv., 0.235 mmol; provided by Milan Urban), (R)-(+)-α-methylbenzylamine (37 mg, 1.3 equiv., 0.306 mmol), 4-nitrophenyl azide (39 mg, 1 equiv., 0.235 mmol), 4 Å molecular sieves (50 mg) and toluene (0.5 mL). Reaction time is overnight. The product was purified by flash column chromatography (first washed with CH_2_Cl_2_ followed by petroleum ether : EtOAc 9:1 → 6:4) affording **5u** (107 mg, 83% yield) as off white solid. m.p. 283 °C. ^1^**H NMR** (600 MHz, CDCl_3_) δ 7.29 (t, *J* = 7.7 Hz, 2H), 7.23 (t, *J* = 7.3 Hz, 1H), 7.15 (d, *J* = 7.8 Hz, 2H), 5.75 (m, 1H), 4.71 (s, 1H), 4.60 (s, 1H), 2.95 (d, *J* = 15.3 Hz, 1H), 2.39 (m, 1H), 2.17 (d, *J* = 15.4 Hz, 1H), 2.03 (d, *J* = 7.0 Hz, 3H), 1.93 (m, 1H), 1.74 – 1.68 (m, 5H), 1.68 – 1.40 (m, 11H), 1.40 – 1.27 (m, 9H), 1.06 (s, 3H), 0.98 (s, 3H), 0.96 (s, 3H), 0.80 (s, 3H), 0.73 (s, 3H). ^13^**C NMR** (151 MHz, CDCl_3_) δ 150.7, 141.9, 141.2, 137.6, 128.6, 127.5, 126.1, 109.5, 77.2, 77.0, 76.8, 59.1, 54.9, 49.2, 48.2, 47.9, 43.0, 42.8, 40.7, 39.9, 38.8, 38.5, 38.2, 35.5, 33.6, 33.3, 29.8, 28.6, 27.5, 25.1, 23.4, 21.5, 19.3, 19.0, 18.0, 15.9, 15.6, 14.5. **HRMS** (ESI^+^): m/z calculated for C_38_H_55_N_3_H [M+H]^+^: 554.4468, found 554.4462.

### Biology

#### Anti-coronavirus evaluation in cell culture

HCoV-229E was purchased from ATCC (VR-740) and expanded in human embryonic lung fibroblast cells (HEL; ATCC^®^ CCL-137). The titers of virus stocks were determined in HEL cells and expressed as TCID_50_ (50% tissue culture infective dose).^49^ The cytopathic effect (CPE) reduction assay was performed in 96-well plates containing confluent HEL cell cultures, as previously described.^50^ Serial compound dilutions were added together with HCoV-229E at an MOI of 100. In parallel, the compounds were added to a mock-infected plate to assess cytotoxicity. Besides the test compounds, two references were included, i.e. K22 [(*Z*)-*N*-[3-[4-(4-bromophenyl)-4-hydroxypiperidin-1-yl]-3-oxo-1-phenylprop-1-en-2-yl]benzamide;^16^ from ChemDiv] and GS-441524 (the nucleoside form of remdesivir; from Carbosynth). After five days incubation at 35°C, microscopy was performed to score virus-induced CPE. To next perform the colorimetric MTS cell viability assay, the reagent (CellTiter 96^®^ AQ_ueous_ MTS Reagent from Promega) was added to the wells, and 24 h later, absorbance at 490 nm was measured in a plate reader. Antiviral activity was calculated from three independent experiments and expressed as EC_50_ or concentration showing 50% efficacy in the MTS or microscopic assay (see reference^51^ for calculation details). Cytotoxicity was expressed as 50% cytotoxic concentration (CC_50_) in the MTS assay.

#### Immunofluorescence detection of viral dsRNA

Semiconfluent 16HBE cell cultures in 8-well chamber slides (Ibidi) were infected with HCoV-229E (MOI: 1000) in the presence of 12 μM of **5h** or GS-441524. After 4 h incubation at 35°C, the inoculum was removed, the compound was added again and the slides were further incubated. At 24 h p.i., the cells were subjected to immunostaining for dsRNA (all incubations at room temperature). After cell fixation with 3.7% formaldehyde in PBS for 15 min, and permeabilization with 0.2% Triton X-100 in PBS for 10 min, unspecific binding sites were blocked with 1% BSA in PBS for 30 min. Next, 1 h incubation was done with mouse monoclonal anti-dsRNA antibody (J2, SCICONS English & Scientific Consulting Kft; diluted 1:1000 in PBS with 1% BSA,), followed by 1 h incubation with goat anti-mouse AlexaFluor488 (A21131, Invitrogen; 1:1000 in PBS with 1% BSA). Cell nuclei were stained with DAPI (4′,6-diamidino-2-phenylindole, Invitrogen) in PBS for 20 min at RT. Microscopic images were acquired using the Leica TCS SP5 confocal microscope (Leica Microsystems) with a HCX PL APO 63x (NA 1.2) water immersion objective. DAPI and AlexaFluor488 were detected with excitation lines at 405 nm (blue) and 488 nm (green), and emission lines of 410-480 nm (blue) and 495–565 nm (green).

#### Time-of-addition assay

Confluent HEL cells were infected with HCoV-229E (MOI: 100) in a 96-well plate, and the compounds [**5h**, bafilomycin A_1_ (from Cayman) or K22] were added at −0.5, 0.5, 2, 4, 6 or 8 h post-infection (p.i.). At 16 h p.i., the supernatant was discarded and each well was washed twice with ice-cold PBS. The cells were lysed on ice for 10 min with 22 μL lysis mix, consisting of lysis enhancer and resuspension buffer at a 1:10 ratio (both from the CellsDirect One-Step RT-qPCR kit; Invitrogen). Next, the lysates were incubated for 10 min at 75°C and treated with DNase (Invitrogen) to remove interfering cellular DNA. The number of viral RNA copies in each sample was determined by one-step RT-qPCR. Five μL lysate was transferred to a qPCR plate containing 9.75 μL of RT-qPCR mix (CellsDirect One-Step RT-qPCR) and 0.25 μL Superscript III RT/Platinum Taq enzyme, and HCoV-229E N-gene specific primers and probe.^52^ The RT-qPCR protocol consisted of 15 min at 50°C; 2 min at 95°C; and 40 cycles of 15 s at 95°C and 45 s at 60°C. An N-gene plasmid standard was included to allow absolute quantification of viral RNA genome copies. The data from two independent experiments were expressed as the number of viral RNA copies at 16 h p.i. relative to the virus control receiving no compound.

#### Selection of resistant coronavirus mutants

HEL cells were infected with HCoV-229E virus (MOI: 25) and **5h** was added at different concentrations. After five days incubation at 35°C, the CPE was scored microscopically to estimate the EC_50_ value. From the highest compound concentration conditions showing virus-induced CPE, the supernatants plus cells were frozen at −80°C. These harvests were further passaged in HEL cells under gradually increasing compound concentrations, until a manifest increase in EC_50_ was observed. A no compound control condition was passaged in parallel. The final virus passages were submitted to RNA extraction; reverse transcription; high-fidelity PCR; and cycle sequencing on the entire viral genome using a set of 39 primers (sequences available upon request). After sequence assembly with CLC Main Workbench 7.9.1 (Qiagen), the sequences of the viruses passaged in the absence and presence of **5h** were aligned in order to identify the **5h** resistance sites in the HCoV-229E genome.

#### Virus yield assay

The virus yield assay was performed in 96-well plates with semiconfluent cultures of human bronchial epithelial 16HBE cells (a gift from P. Hoet, Leuven, Belgium) or confluent HEL cells. Serial dilutions of compound **5h** were added and the cells were infected (MOI: 100) with wild-type HCoV-229E (229E-WT), EndoU-deficient HCoV-229E (229E-H250A_nsp15_)^22^ or the mutant viruses obtained by passaging under **5h** (229E-K60R_nsp15_ or 229E-T66I_nsp15_). After 4 h incubation at 35°C, the inoculum was removed, the compound dilutions were added again and the plates were further incubated. At three days p.i., the supernatants were collected, and 2 μL of each supernatant was lysed on ice by adding 11 μL lysis mix containing lysis enhancer and resuspension buffer at a 1:10 ratio. The lysates were incubated for 10 min at 75°C and the viral RNA copy number was determined by RT-qPCR as described above for the time-of-addition assay. The data were collected in three independent experiments and expressed as the fold reduction in viral RNA compared to the virus control receiving no compound.

### Computational work

Starting from the published X-ray structure of hexameric nsp15 protein from HCoV-229E (PDB code: 4RS4), we first introduced a series of mutations, i.e. S10Q, S17G, A142T, M219I and S252L, to obtain the nsp15 protein sequence identical to that of the HCoV-229E virus used in the biological experiments. The structures of HCoV-229E nsp15 and SARS-CoV-2 nsp 15 (PDB code: 7K1O) were prepared using MOE (Chemical Computing Group, Montreal, Canada). Hydrogen addition and optimisation of protonation state and rotamers of the mutations were conducted using the AMBER-EHT force field, and identification of the potential binding sites in the multimeric complex was performed using MOE. Docking of betulonic acid derivatives was carried out by means of both MOE and GOLD with default settings, where GBVI/WSA score and Goldscore functions were used, respectively. The common top scoring solution was selected for further research.

## ANCILLARY INFORMATION

### SUPPORTING INFORMATION

^1^H NMR and ^13^C NMR spectra of the synthesized compounds.

### CURRENT AUTHOR ADDRESS

Joice Thomas, Arcus Biosciences, Hayward, CA 94545, United States.

### AUTHOR INFORMATION

The authors declare no conflict of interest.

### AUTHOR CONTRIBUTIONS

B.K. and A.S. contributed equally. B.K. and J.T. performed compound synthesis. A.S. and L.N. designed, performed and interpreted the biological experiments. B.V.L., J.V. and D.J. performed antiviral experiments. T.N. and A.V. performed and interpreted the *in silico* study. V.T. and R.D. provided materials. B.K., A.S., W.D., A.V. and L.N. co-wrote the manuscript. All authors gave approval to the final version of the manuscript.

## ACKNOWLEDGMENT

B.K. acknowledges receipt of an Erasmus Mundus Western Balkan Action 2 and doctoral fellowship from KU Leuven. B.V.L. holds an SB-PhD fellowship from the FWO Research Foundation Flanders (project: 1S66321N). Mass spectrometry was made possible by the support of the Hercules Foundation of the Flemish Government (grant: 20100225-7). We acknowledge the support of NVIDIA Corporation for donating the Titan Xp GPU used for this study. This research work was supported by funding from the KU Leuven (IDO/12/020 and project C14/19/78); the European Union’s Innovative Medicines Initiative (IMI) under Grant Agreement 101005077 [Corona Accelerated R&D in Europe (CARE) project]; and Fundació La Marató de TV3, Spain (Project No. 201832-30). We thank Amy Swinnen and Leentje Persoons for dedicated technical assistance, and Els Vanstreels for providing confocal microscopy training. We also thank Milan Urban for providing lupenone.

## ABBREVIATIONS USED

CC_50_: 50% cytotoxic concentration
CoV: coronavirus
COVID-19: coronavirus disease 2019
CPE: cytopathic effect
DAPI: 4′,6-diamidino-2-phenylindole
EndoU: endoribonuclease
FIPV: feline infectious peritonitis virus
HEL: human embryonic lung
MERS: Middle East respiratory syndrome
MHV-A59: mouse hepatitis virus A59
MTS: 3-(4,5-dimethylthiazol-2-yl)-5-(3-carboxymethoxyphenyl)-2-(4-sulfophenyl)-2H-tetrazolium
nsp: non-structural protein
p.i.: post-infection
RTC: replication-transcription complex
SARS: severe acute respiratory syndrome
SARS-CoV-2: severe acute respiratory syndrome coronavirus 2
TCID_50_: 50% tissue culture infective dose

