## Supporting Information for "Betulonic acid derivatives inhibiting coronavirus replication in cell culture via the nsp15 endoribonuclease"

**$^1$H and $^{13}$C NMR spectra** **S2-S43**

[1]**H NMR** spectrum of **5a** (400 MHz, $CDCl_3$):

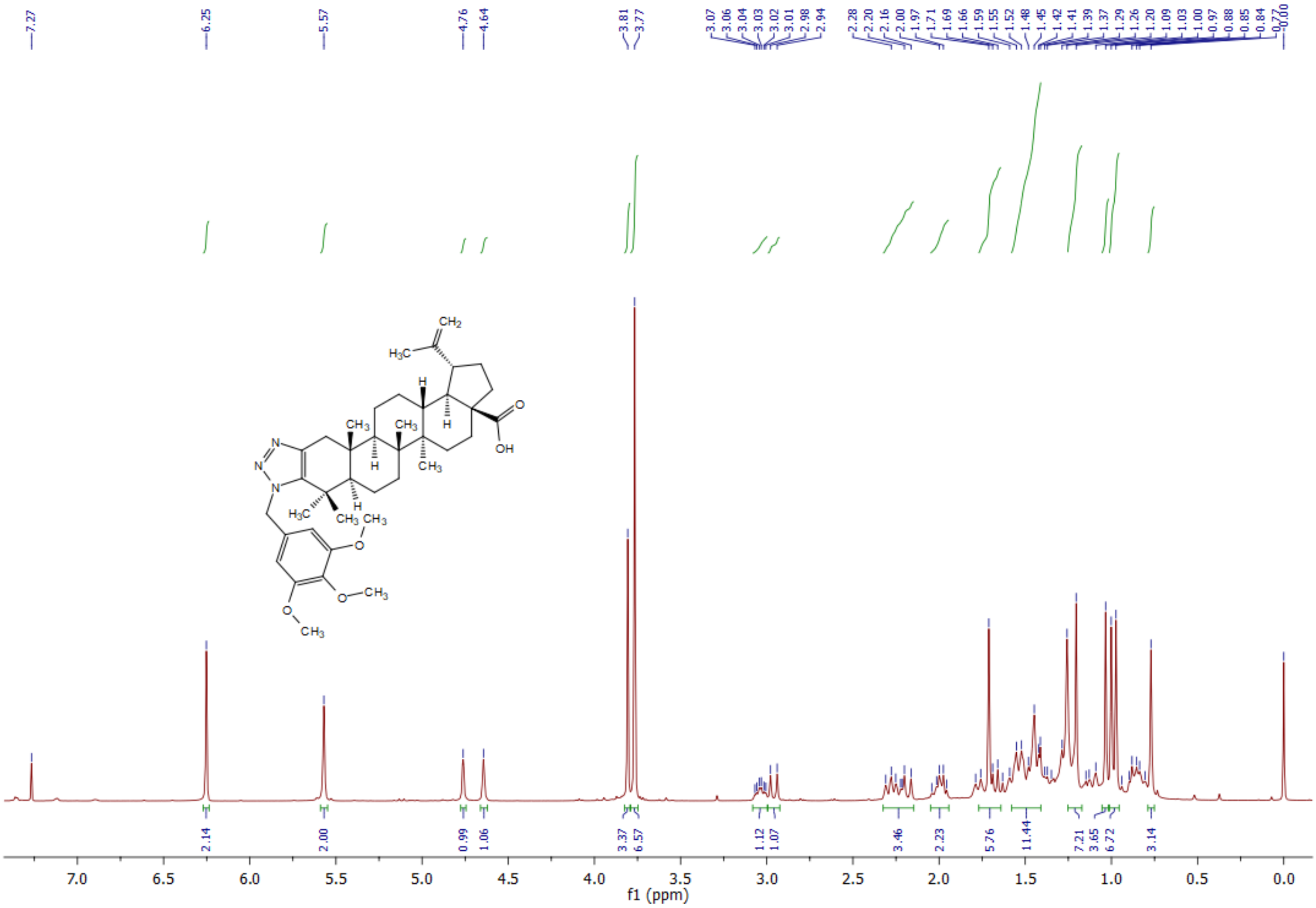

$^{13}$**C NMR** spectrum of **5a** (400 MHz, $CDCl_3$):

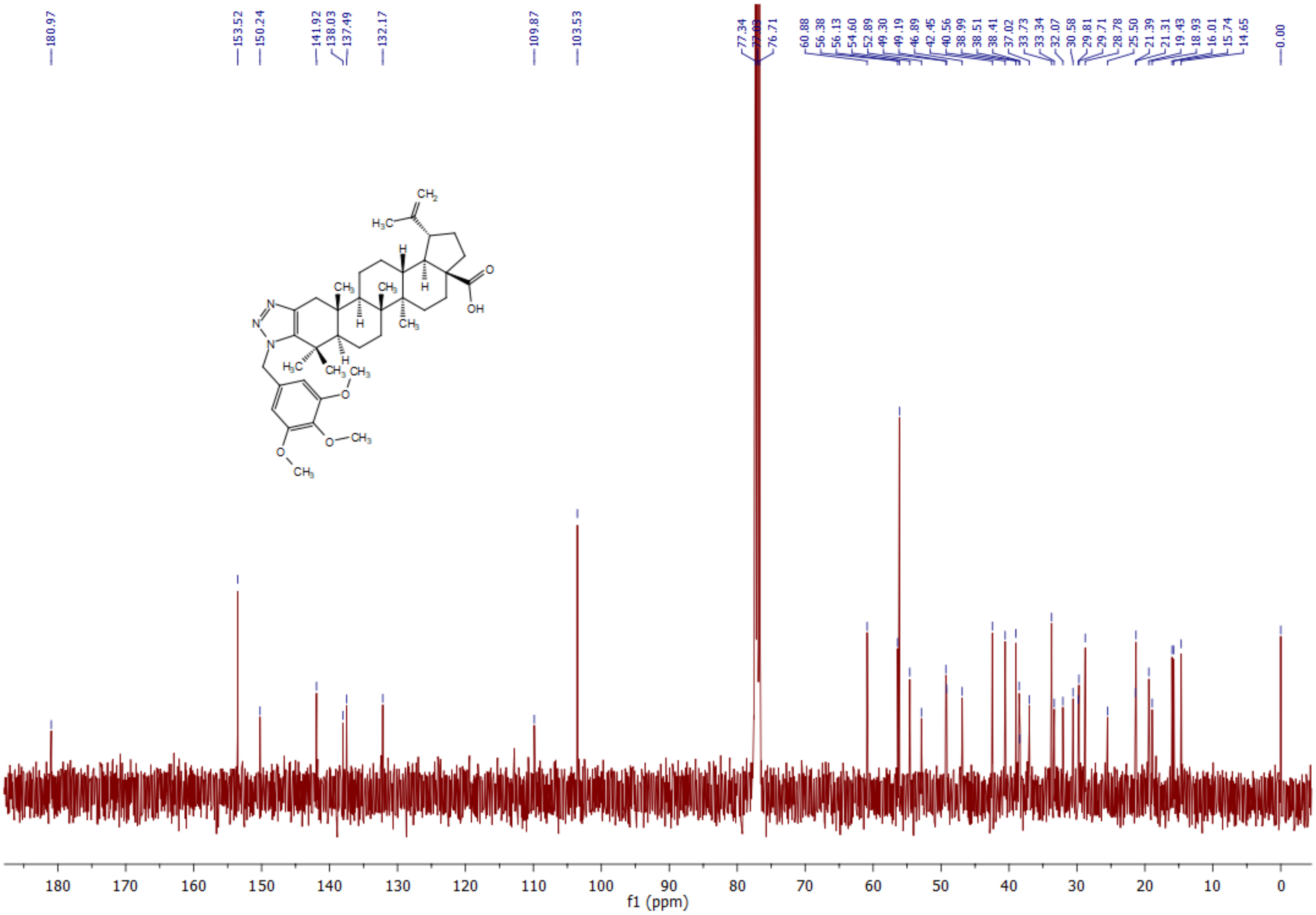

**[1]H NMR** spectrum of **5b** (400 MHz, $CDCl_3$):

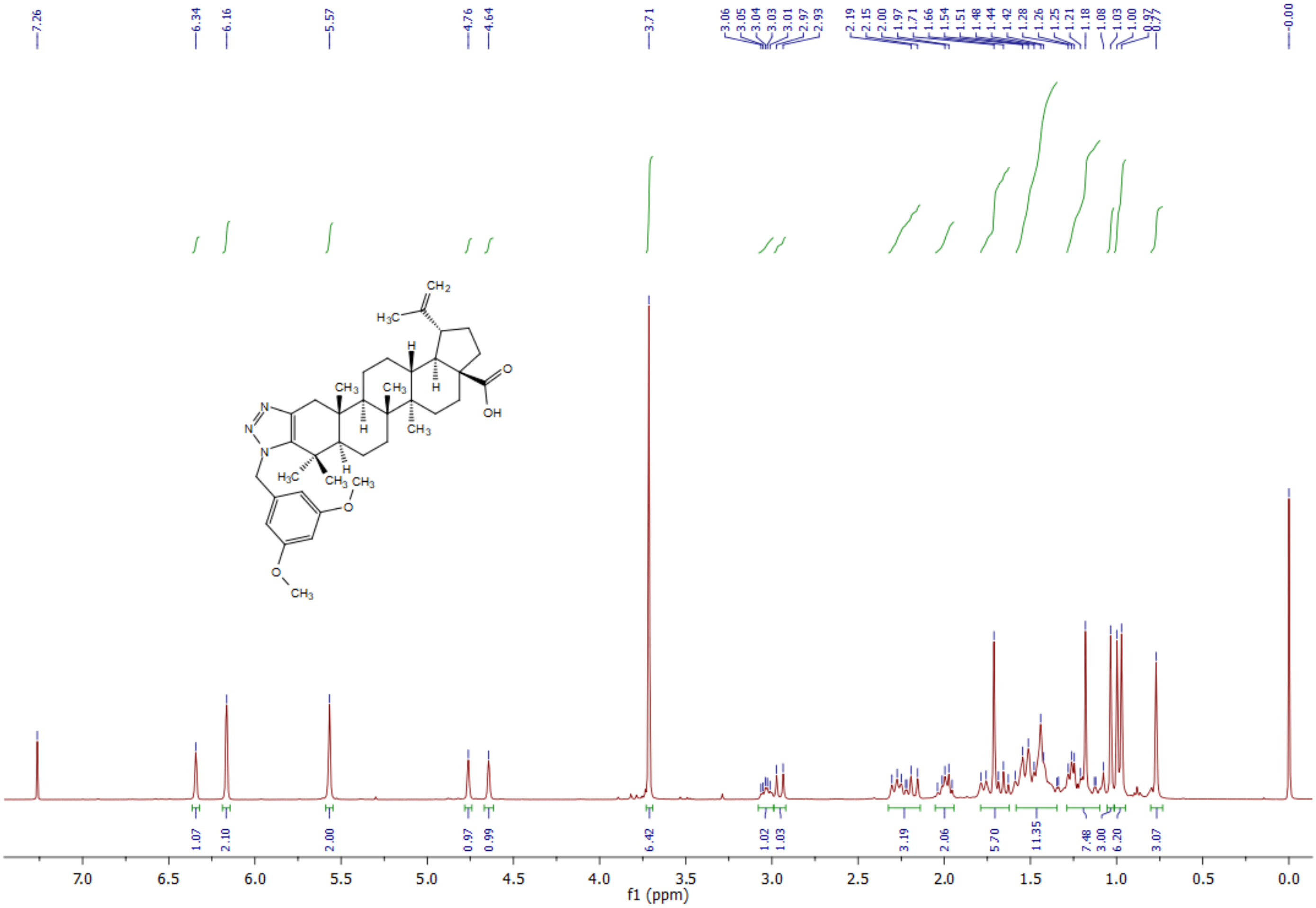

**$^{13}$C NMR** spectrum of **5b** (400 MHz, $CDCl_3$):

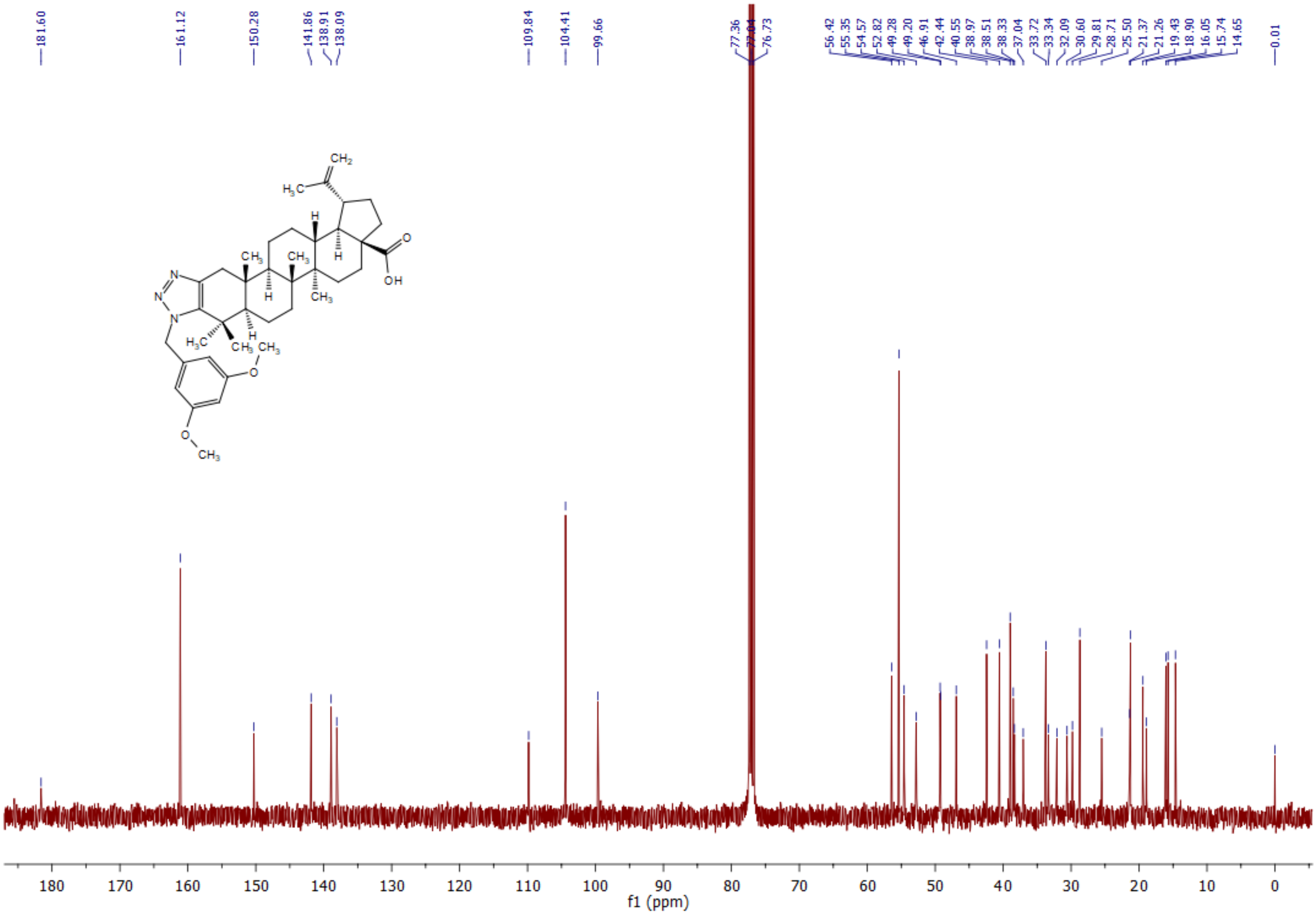

[1]**H NMR** spectrum of **5c** (400 MHz, $CDCl_3$):

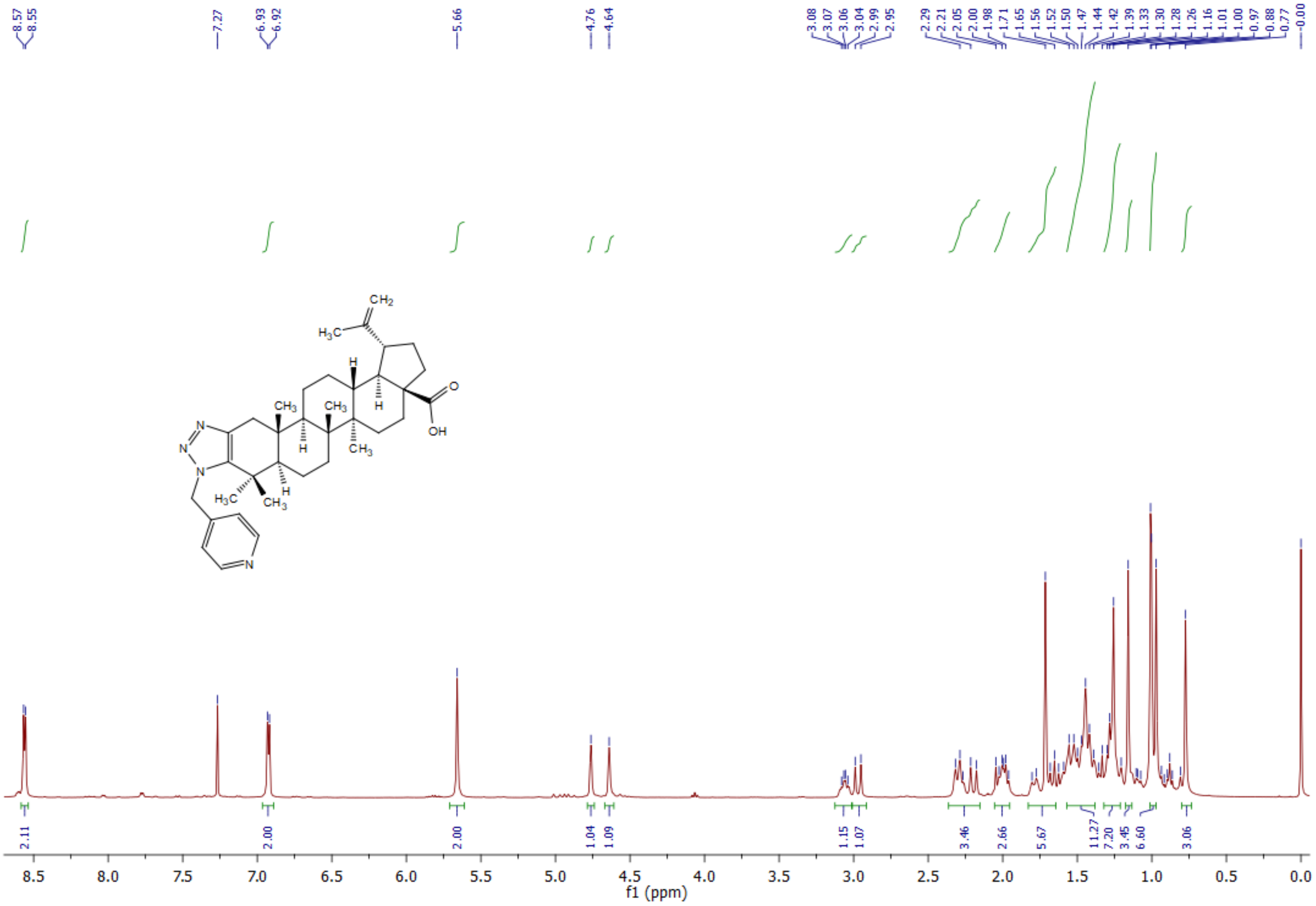

**$^{13}$C NMR** spectrum of **5c** (400 MHz, $CDCl_3$):

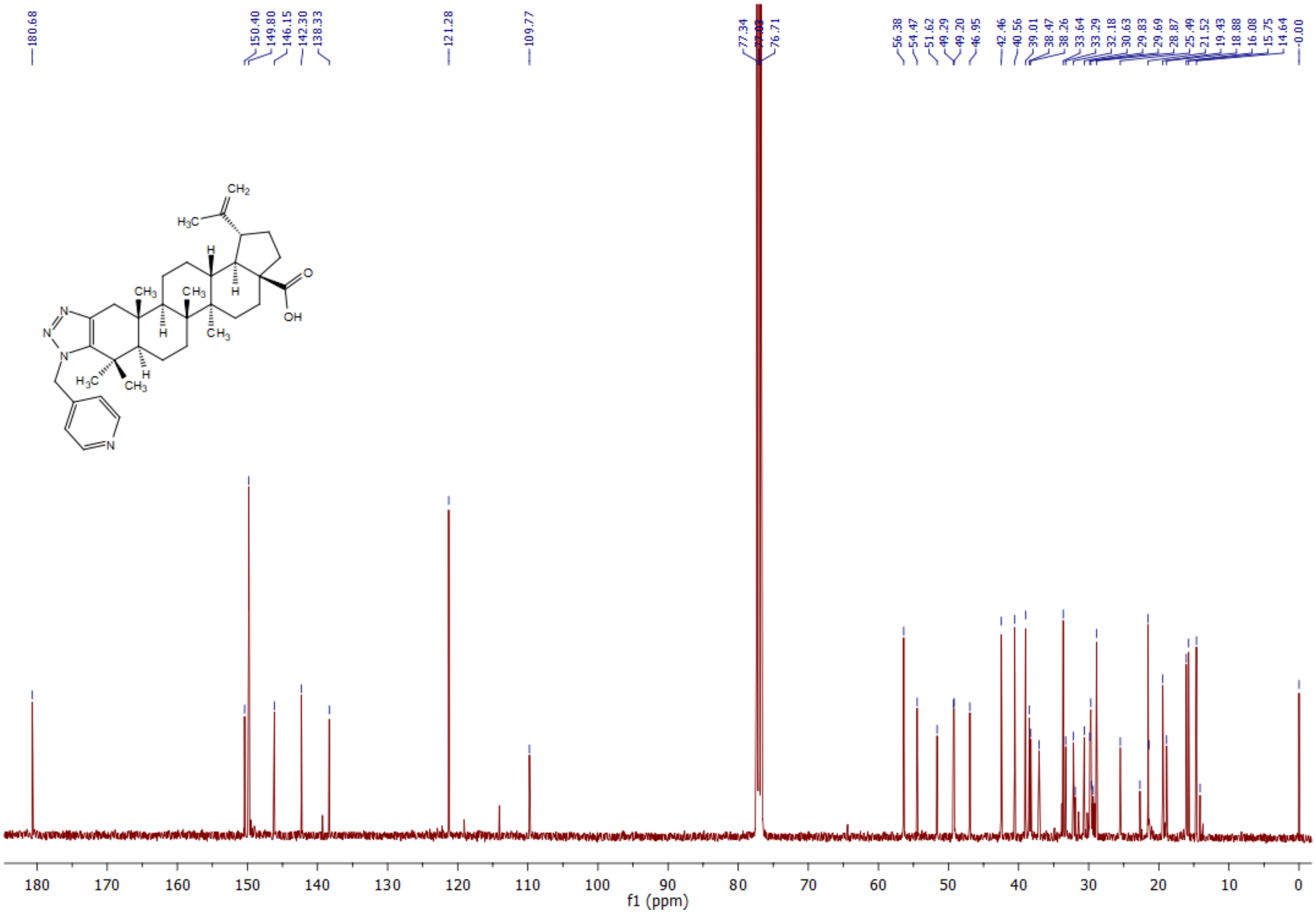

[1]H NMR spectrum of **5d** (600 MHz, $CDCl_3$):

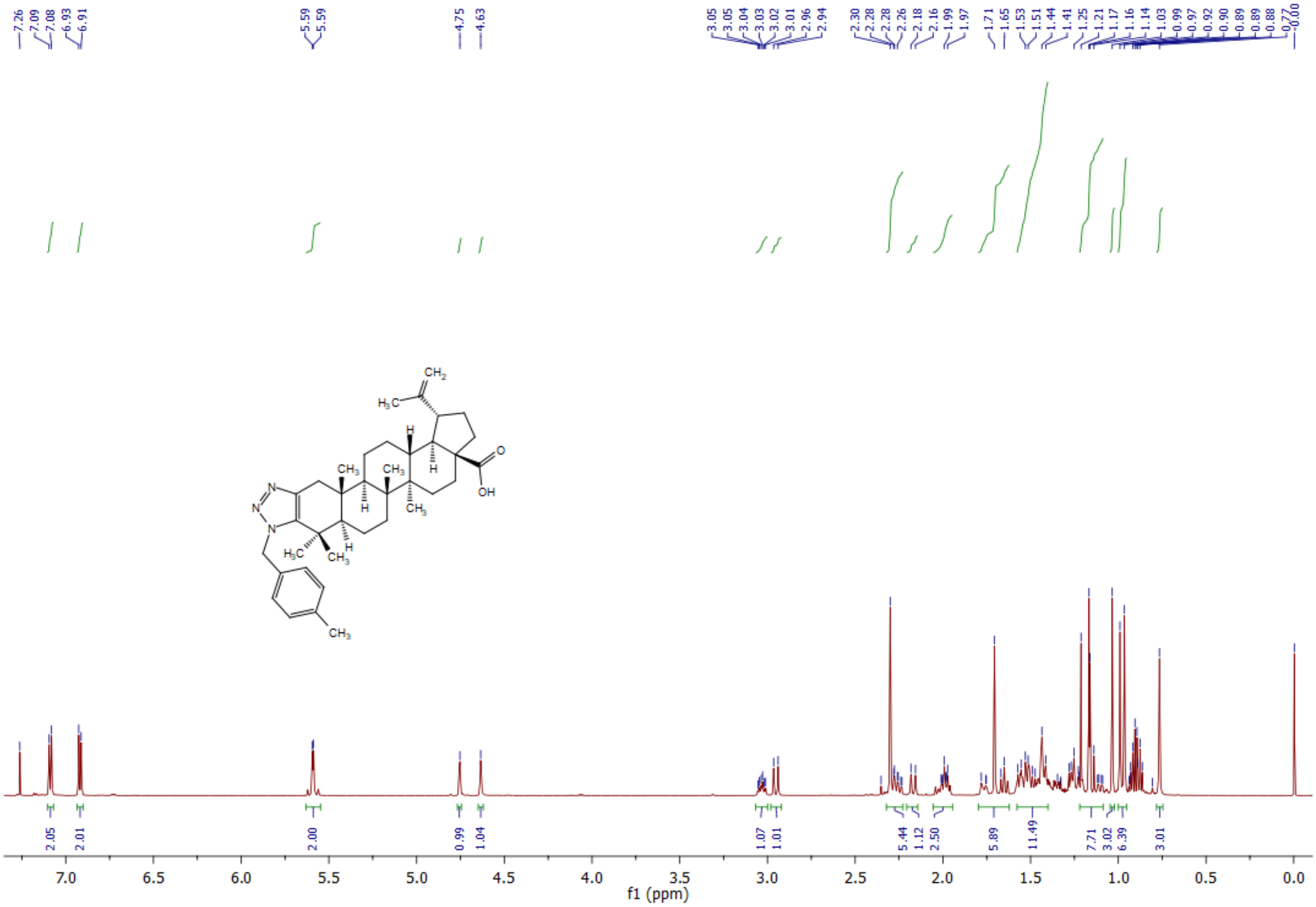

**$^{13}$C NMR** spectrum of **5d** (400 MHz, $CDCl_3$):

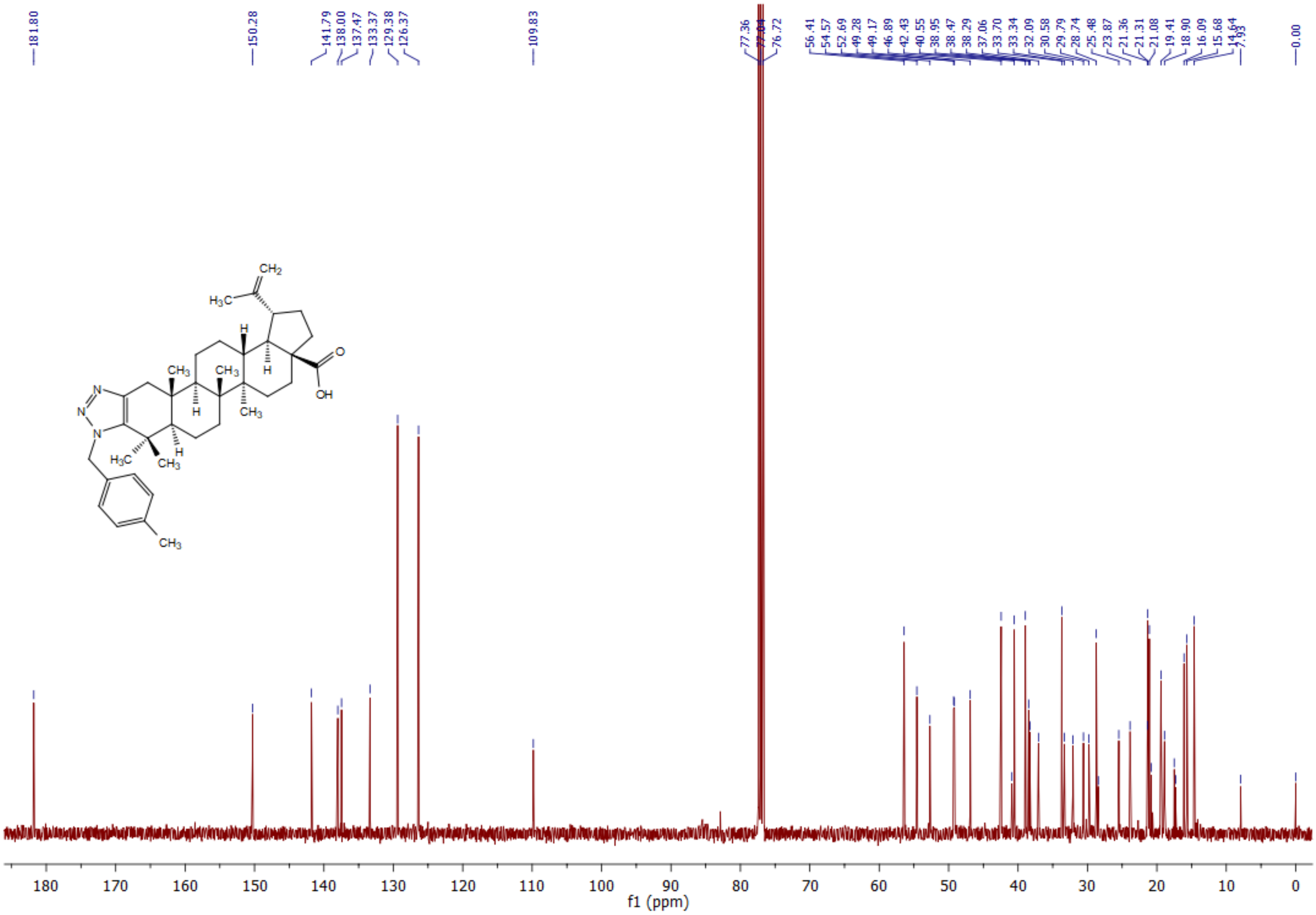

**$^1$H NMR** spectrum of **5e** (400 MHz, $CDCl_3$):

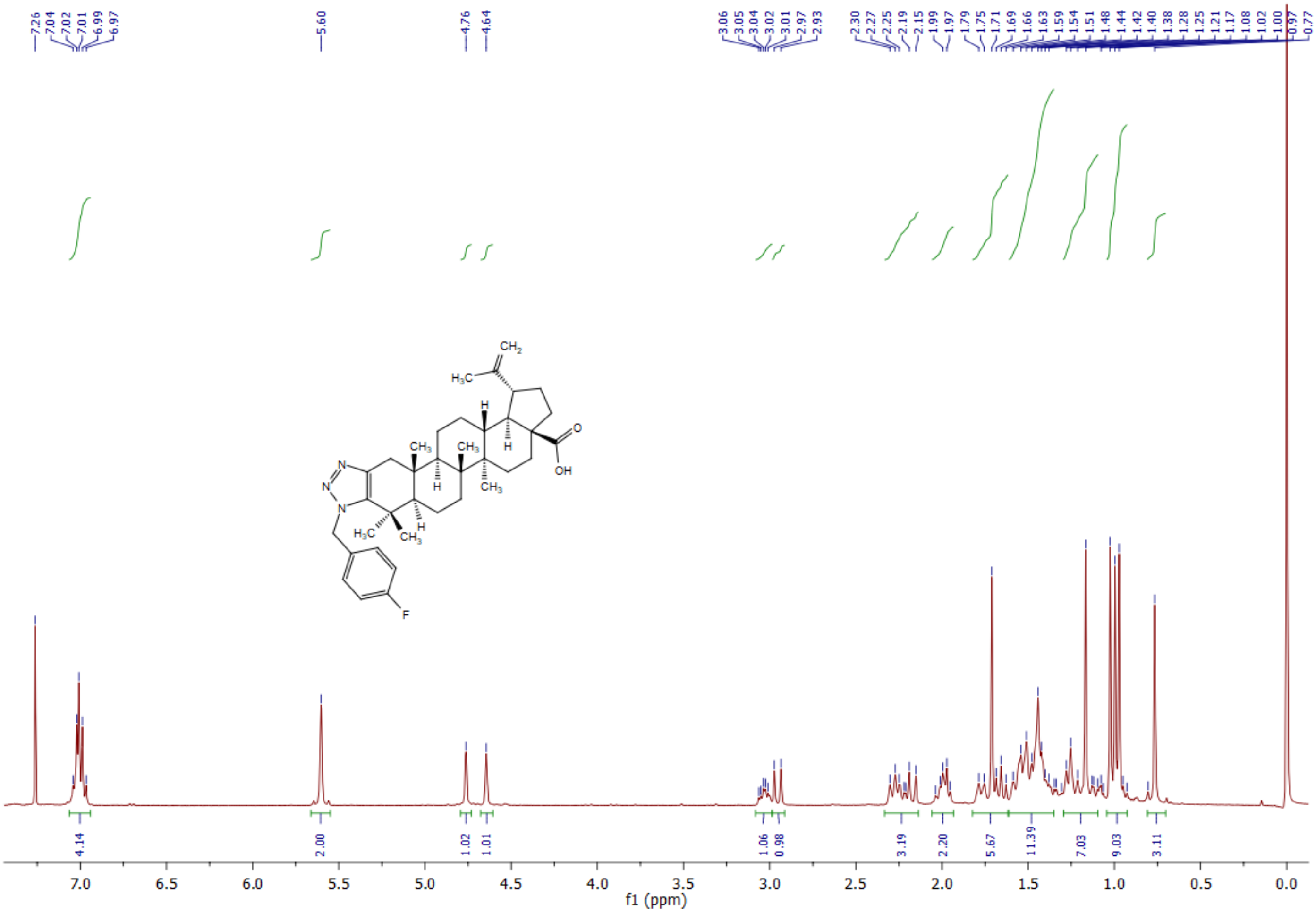

**$^{13}$C NMR** spectrum of **5e** (400 MHz, $CDCl_3$):

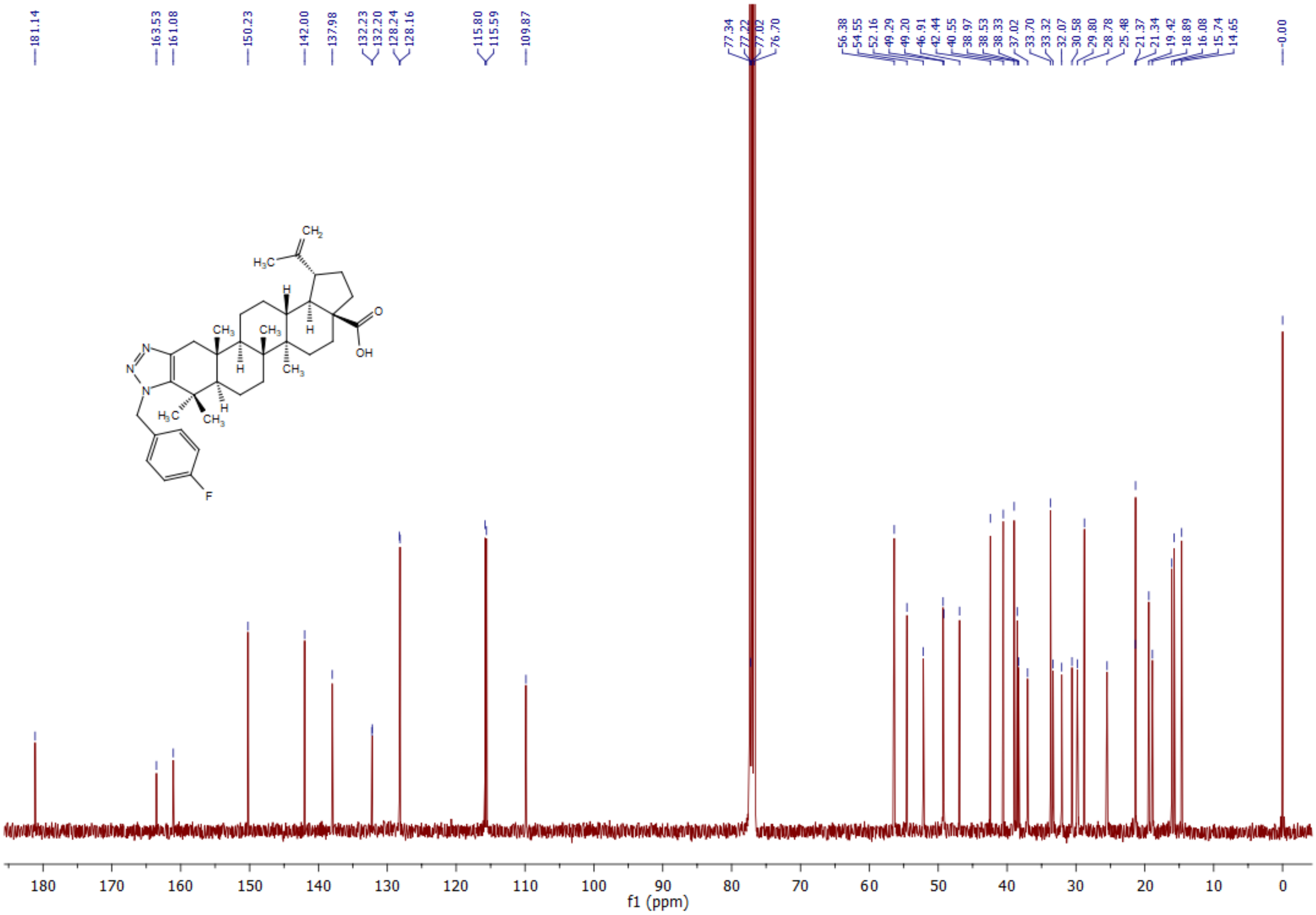

$^1$**H NMR** spectrum of **5f** (400 MHz, $CDCl_3$):

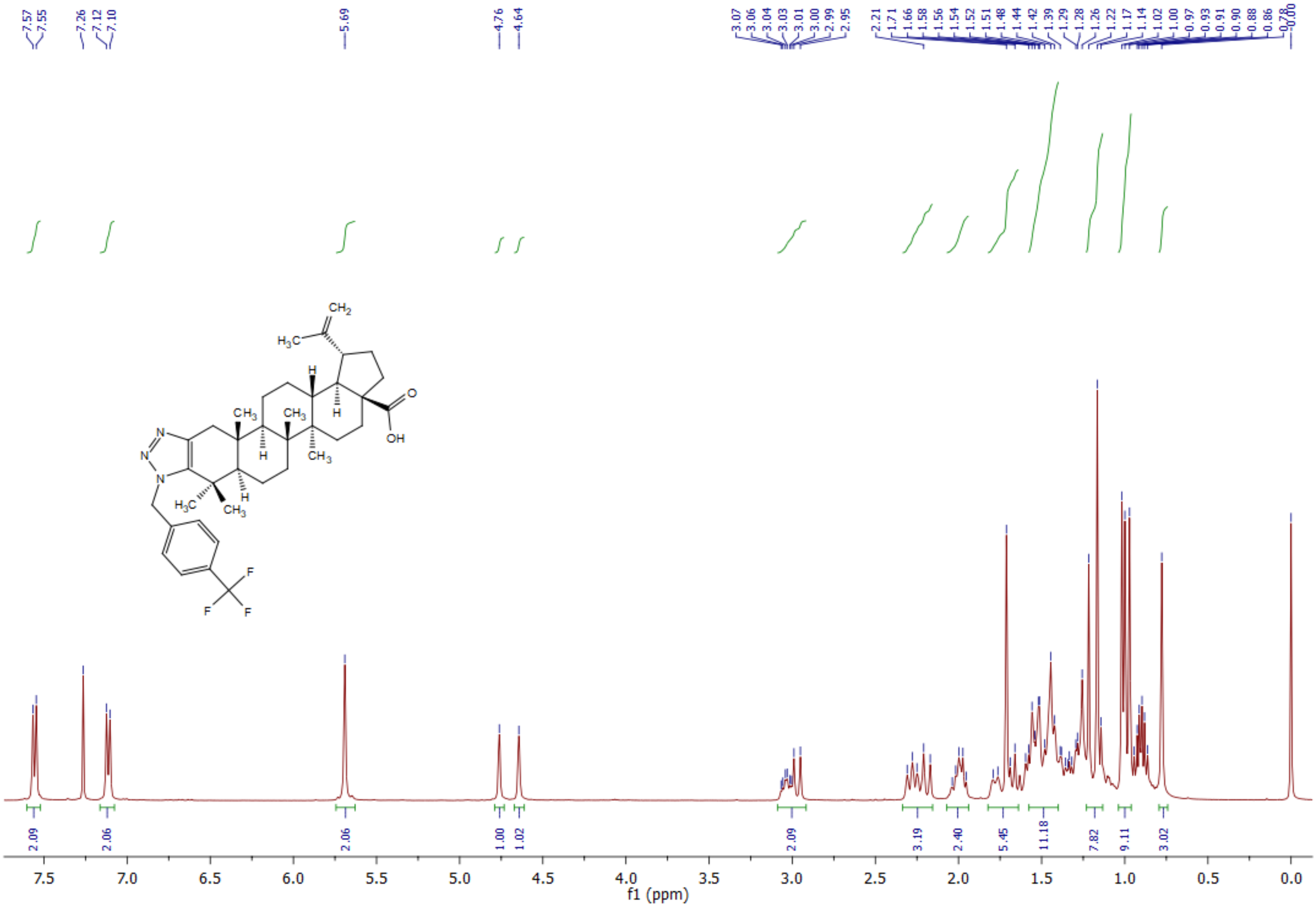

$^{13}$**C NMR** spectrum of **5f** (400 MHz, $CDCl_3$):

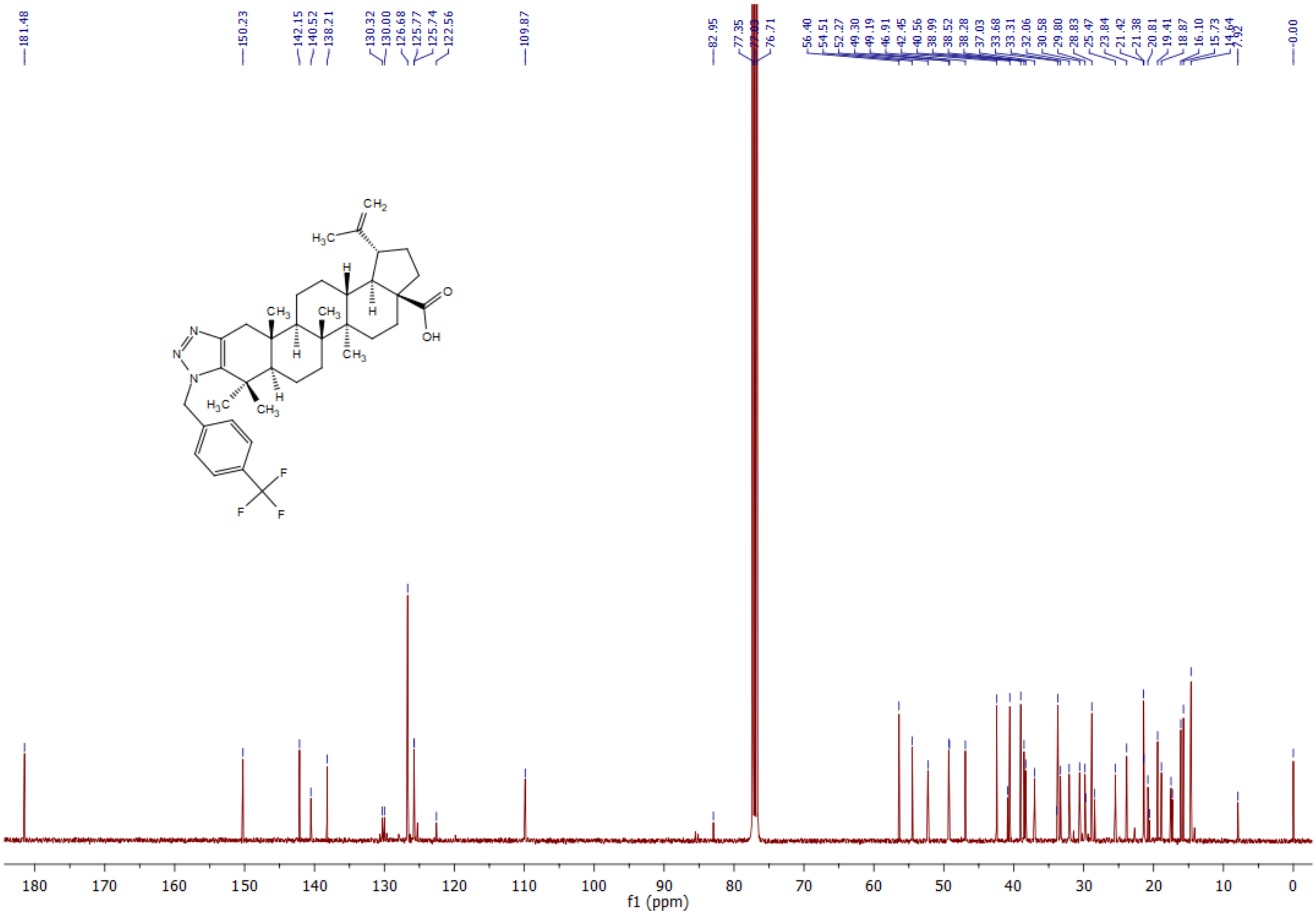

[1]**H NMR** spectrum of **5g** (400 MHz, $CDCl_3$):

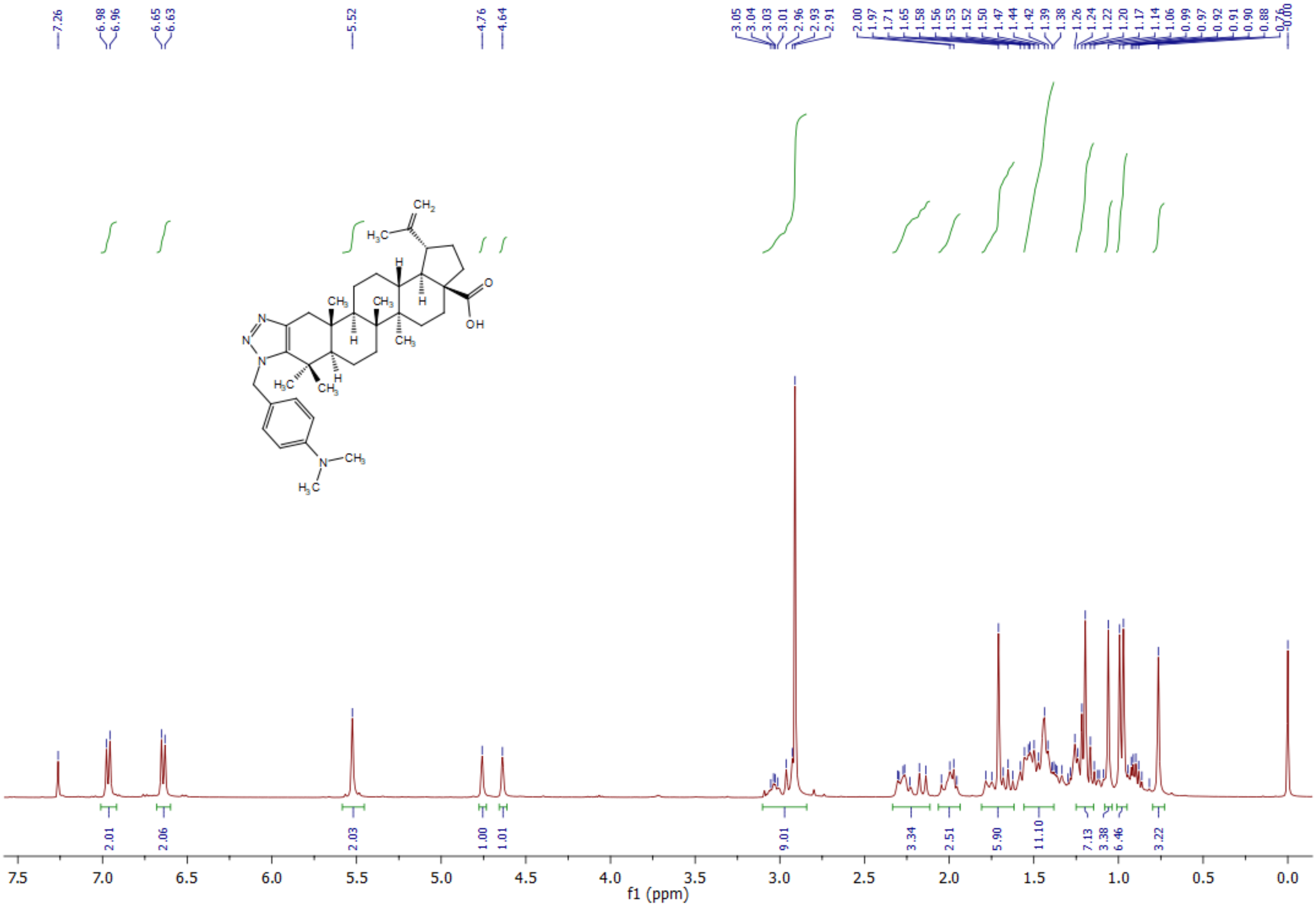

**$^{13}C$ NMR** spectrum of **5g** (400 MHz, $CDCl_3$):

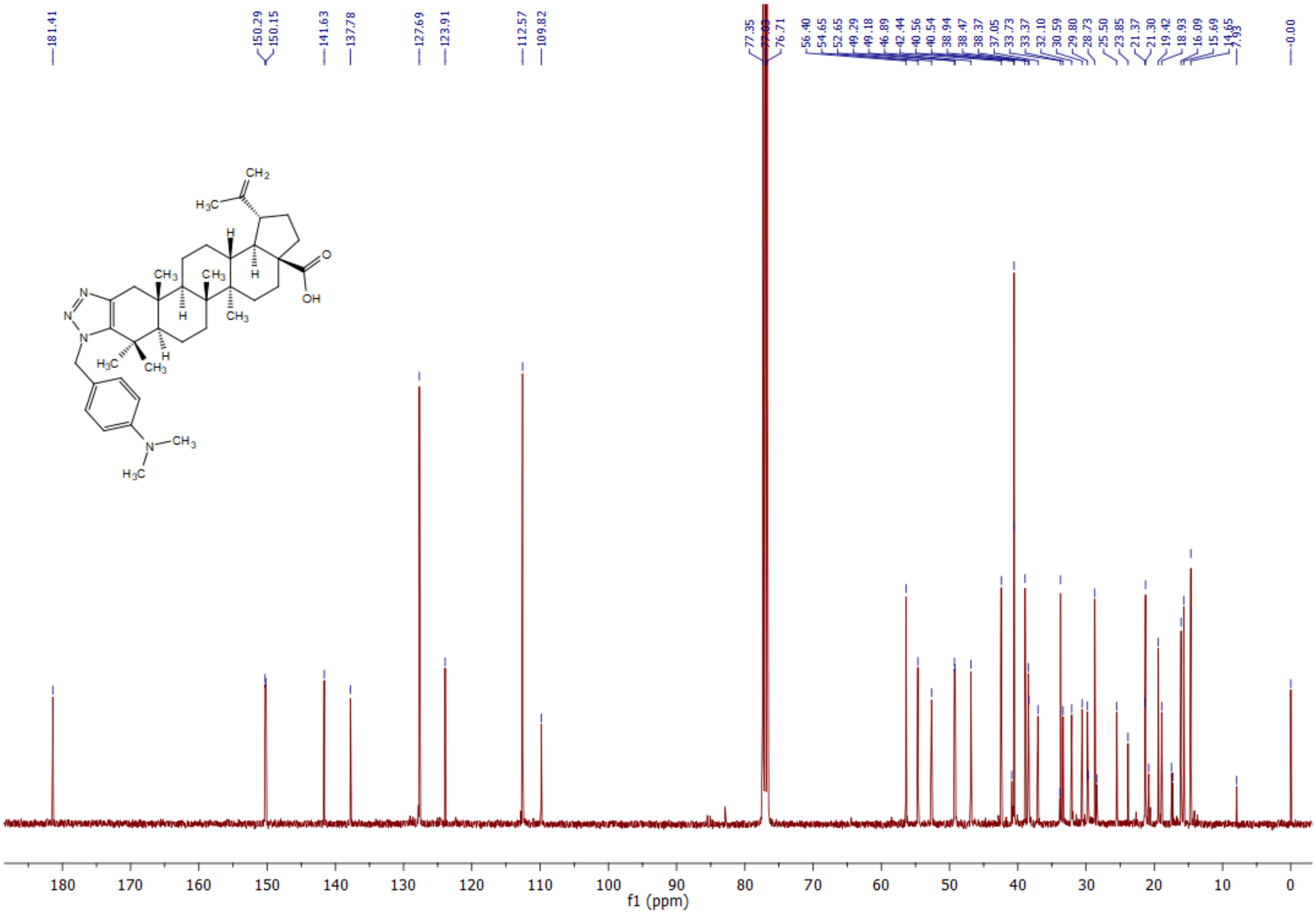

[1]H NMR spectrum of 5h (400 MHz, $CDCl_3$):

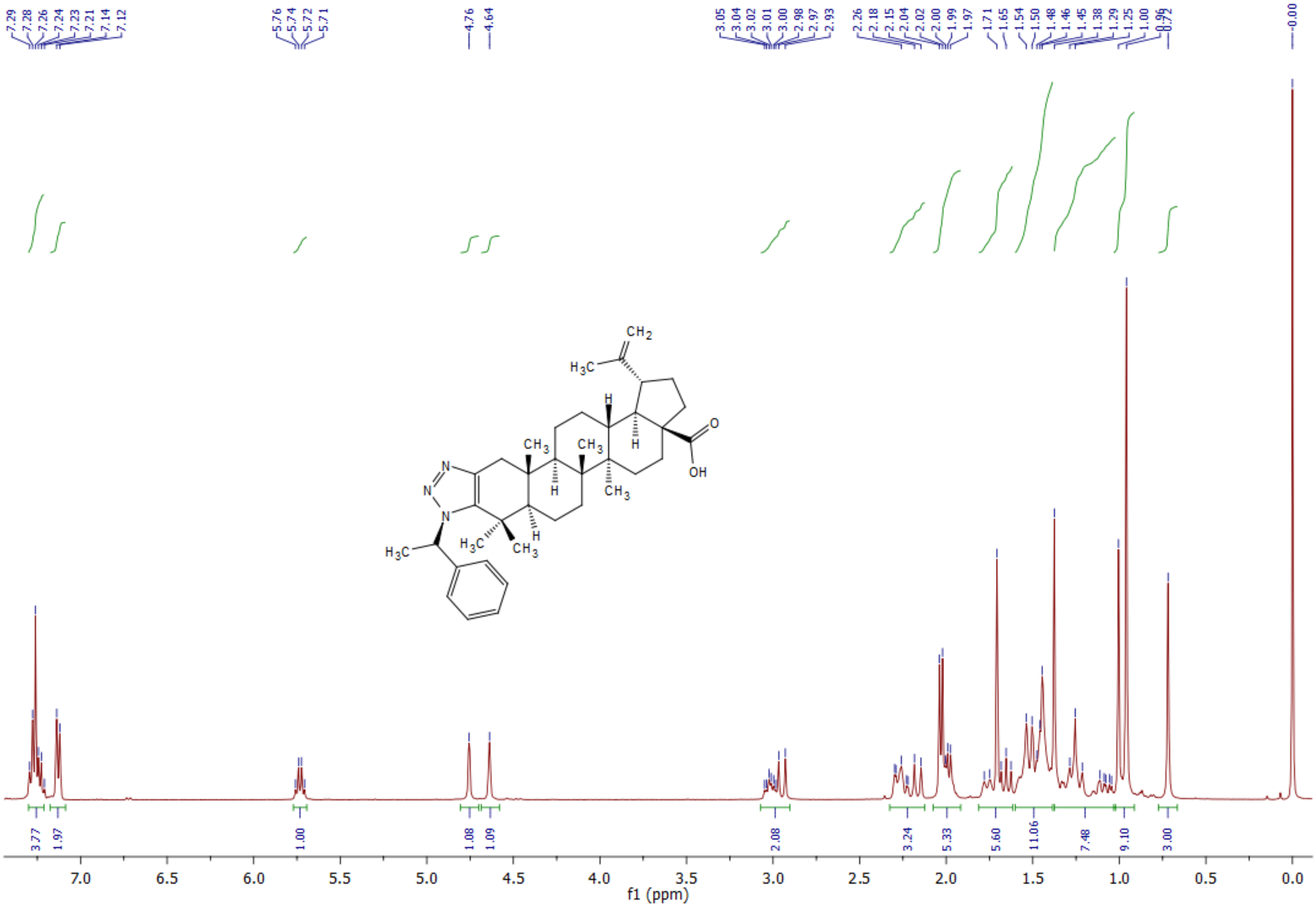

**$^{13}$C NMR** spectrum of **5h** (400 MHz, $CDCl_3$):

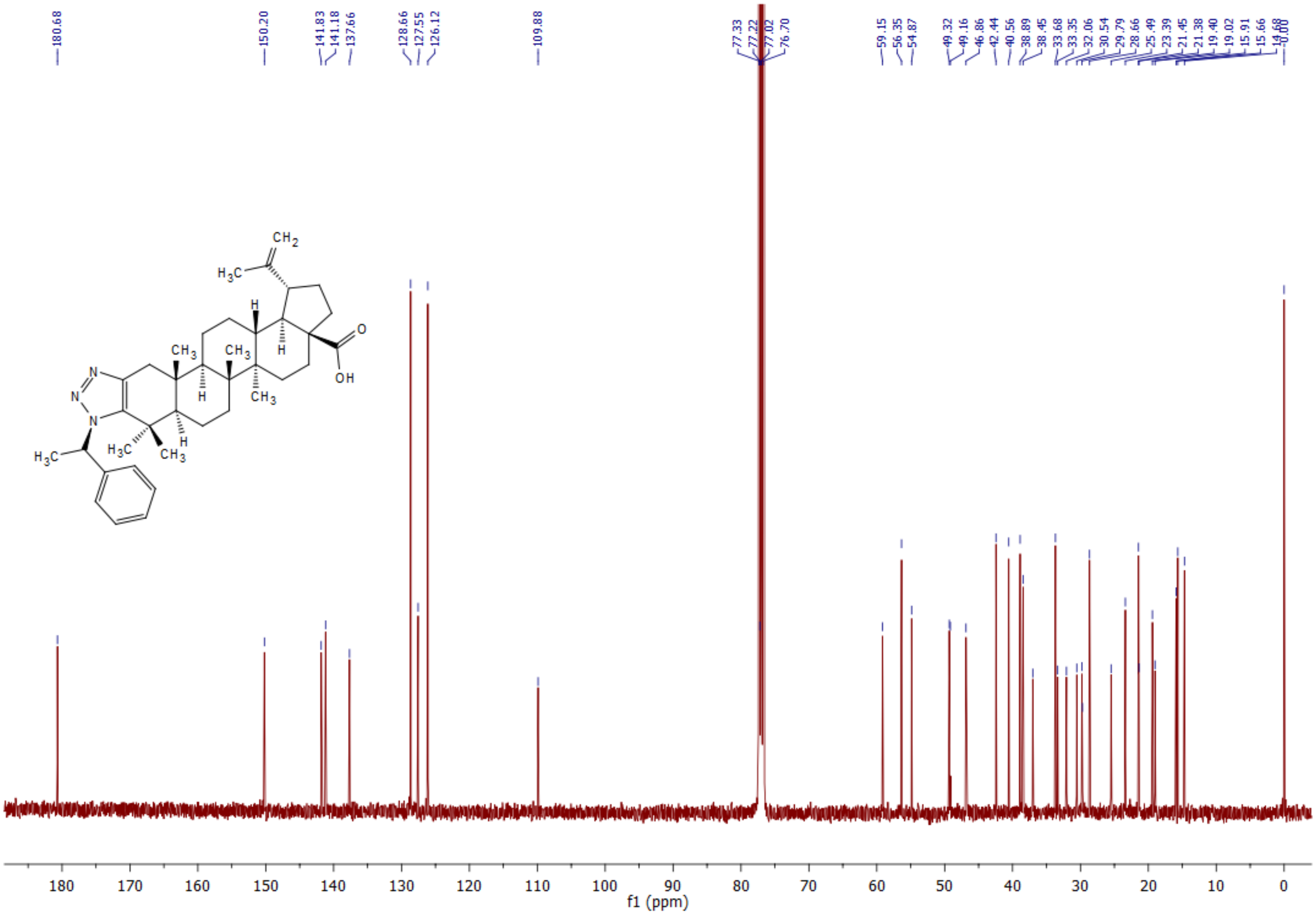

[1]H NMR spectrum of **5i** (600 MHz, $CDCl_3$):

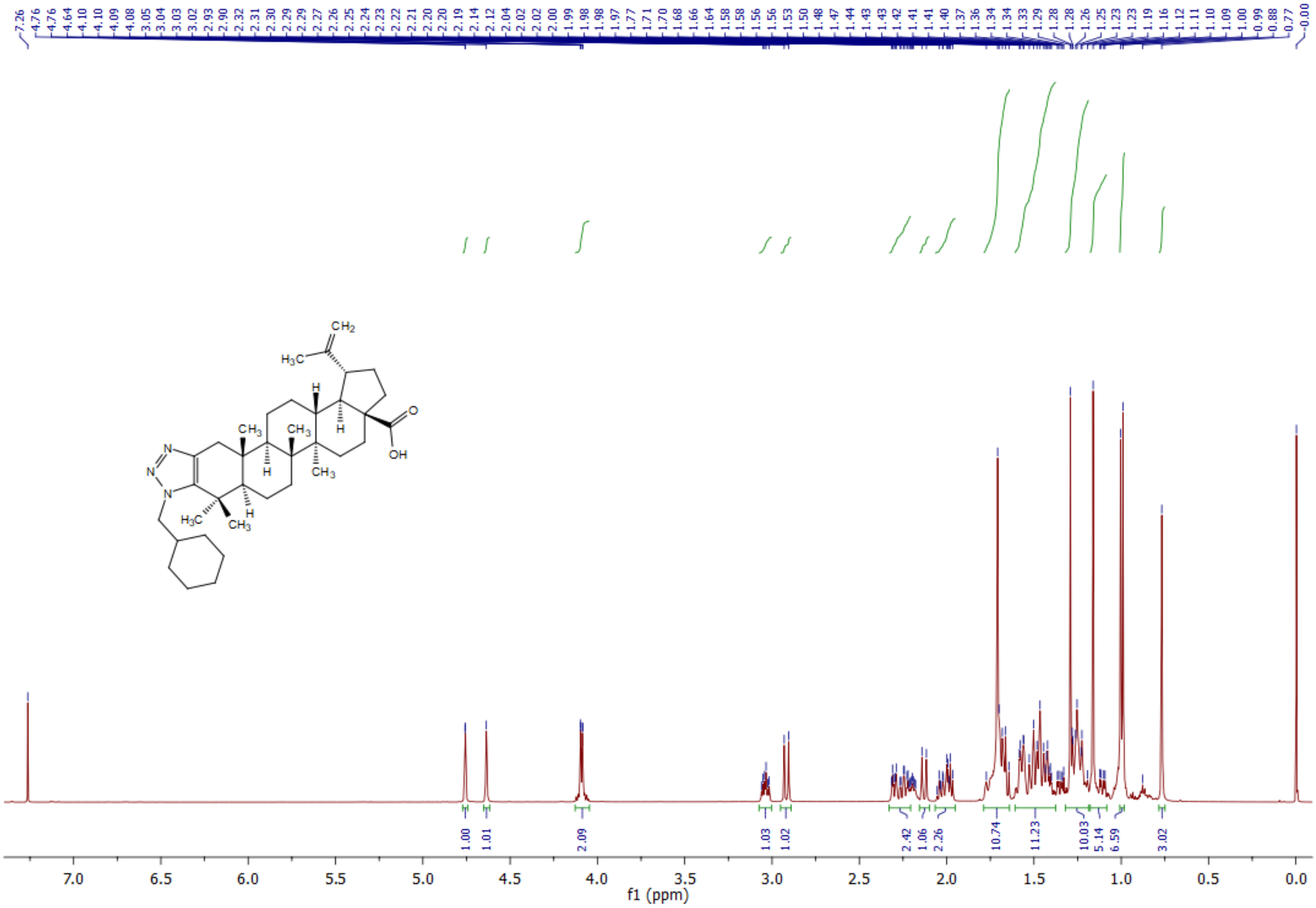

**$^{13}$C NMR** spectrum of **5i** (400 MHz, $CDCl_3$):

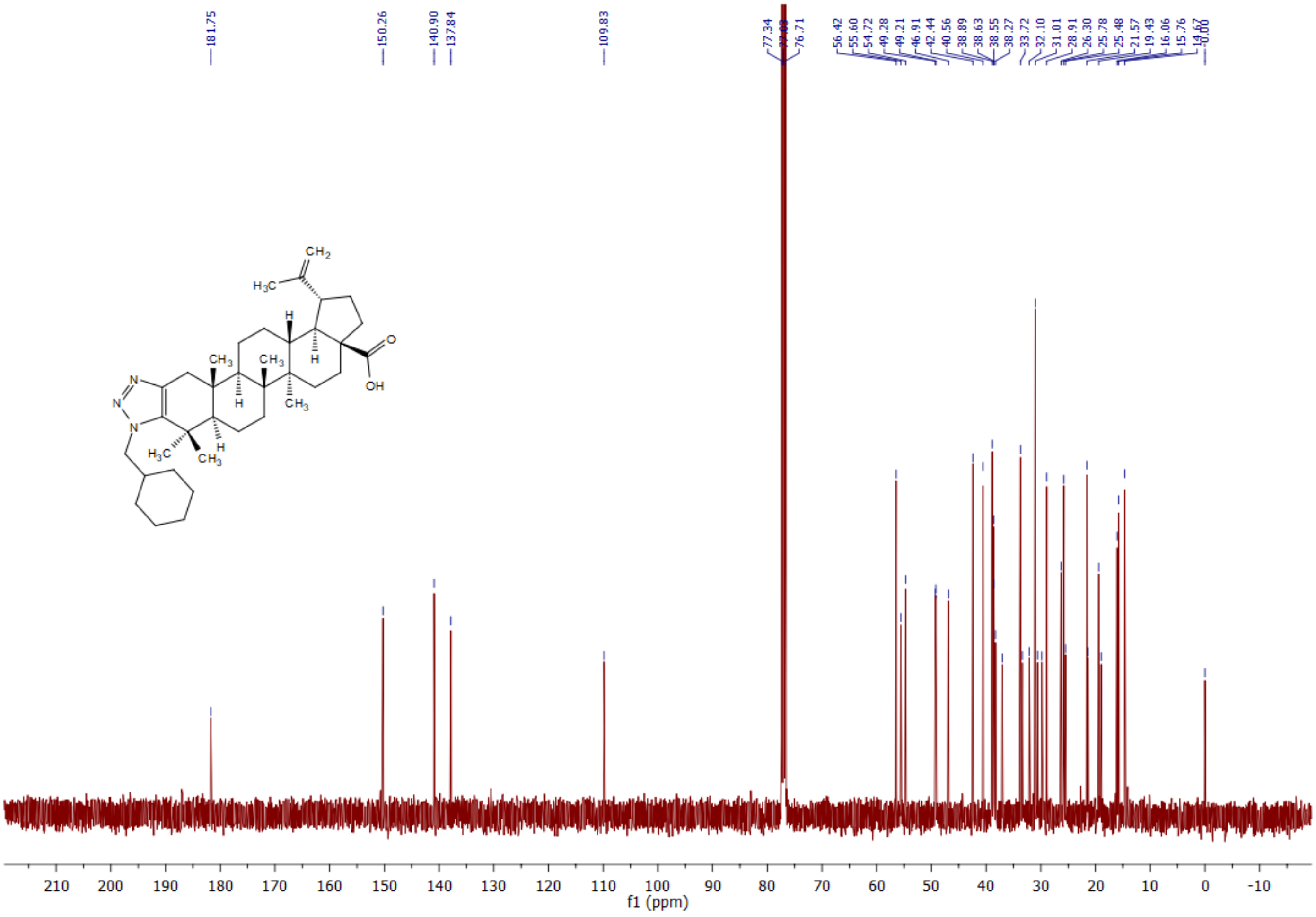

$^1$**H NMR** spectrum of **5j** (300 MHz, $CDCl_3$):

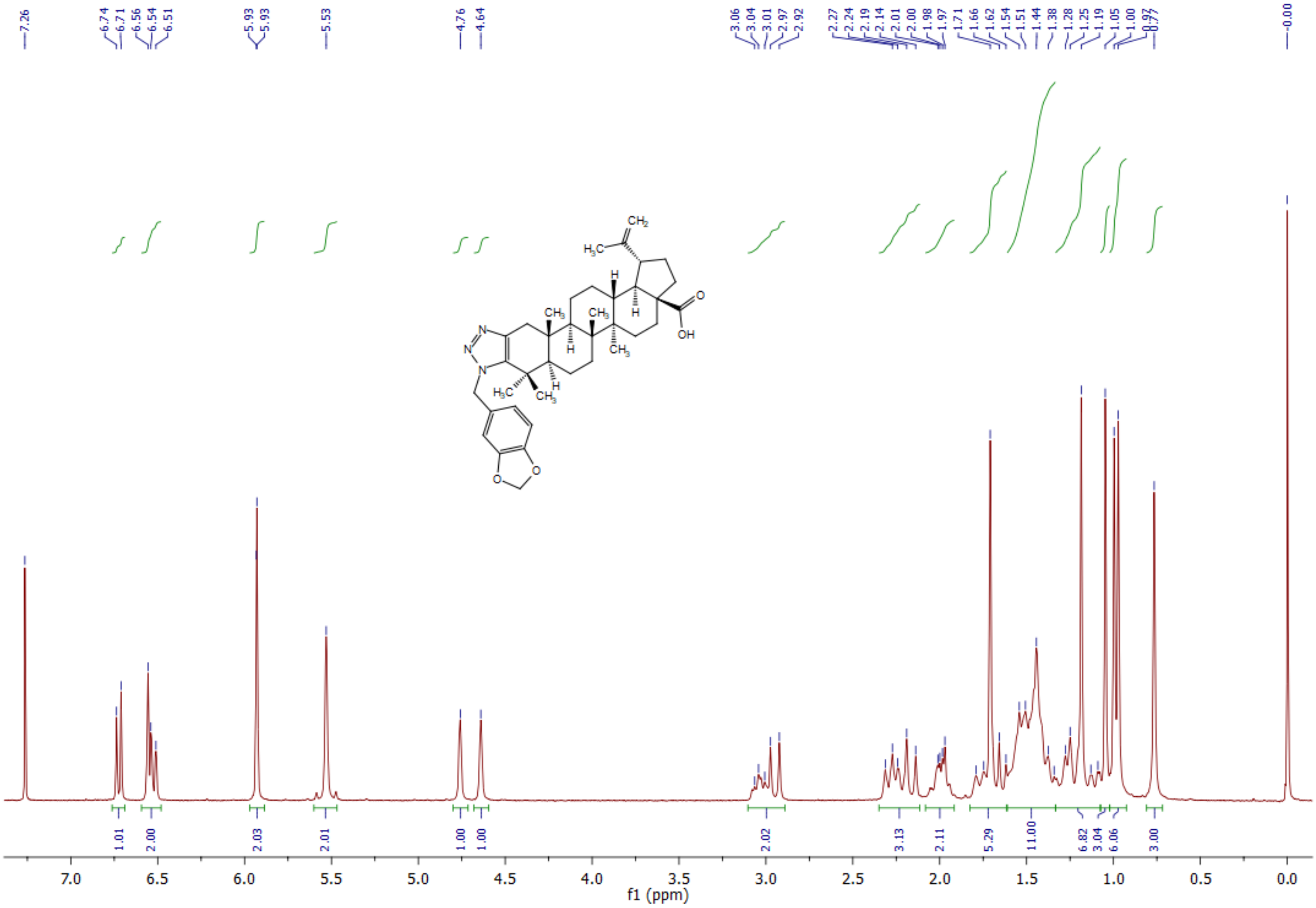

**$^{13}$C NMR** spectrum of **5j** (400 MHz, $CDCl_3$):

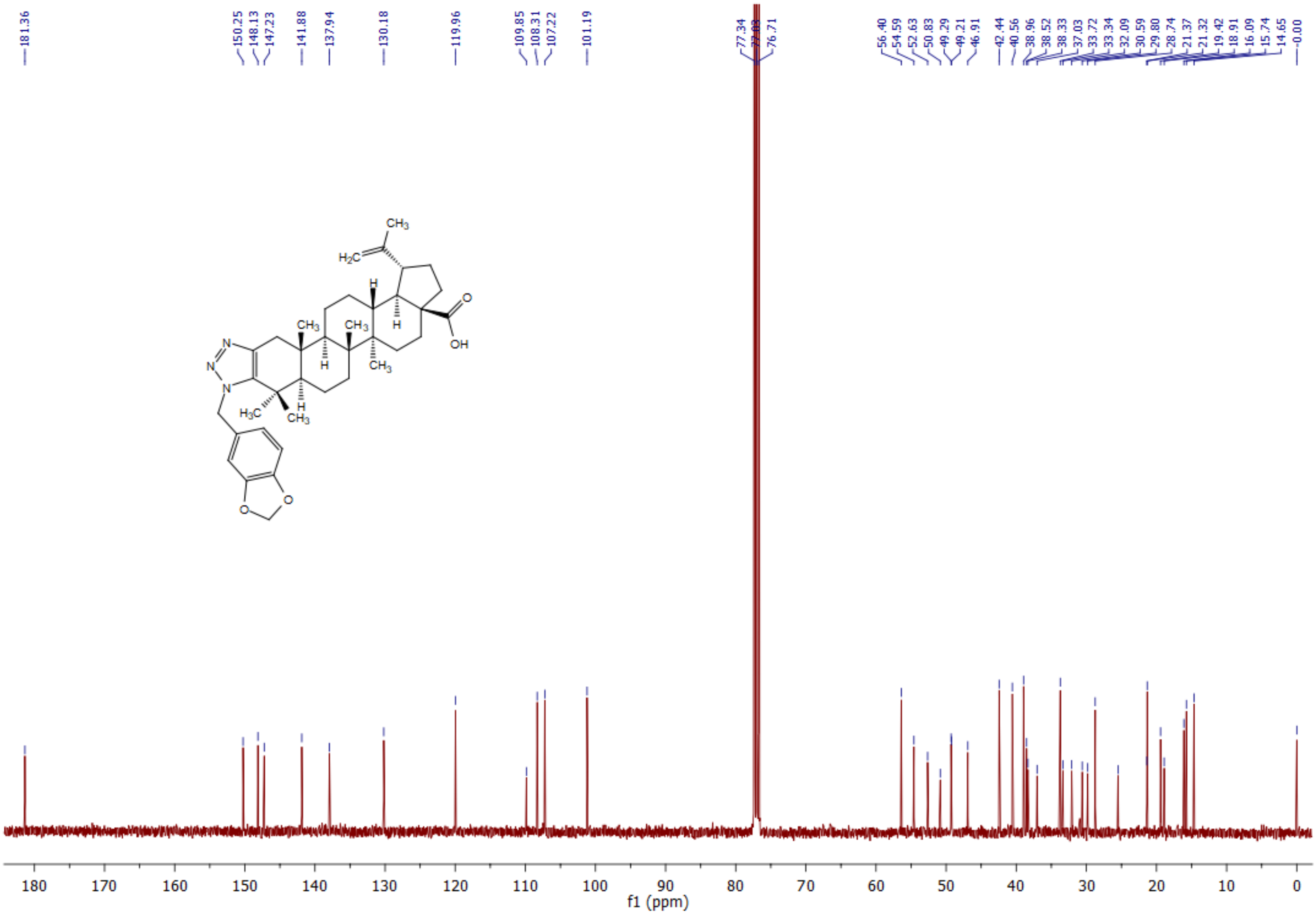

**[1]H NMR** spectrum of **5k** (400 MHz, $CDCl_3$):

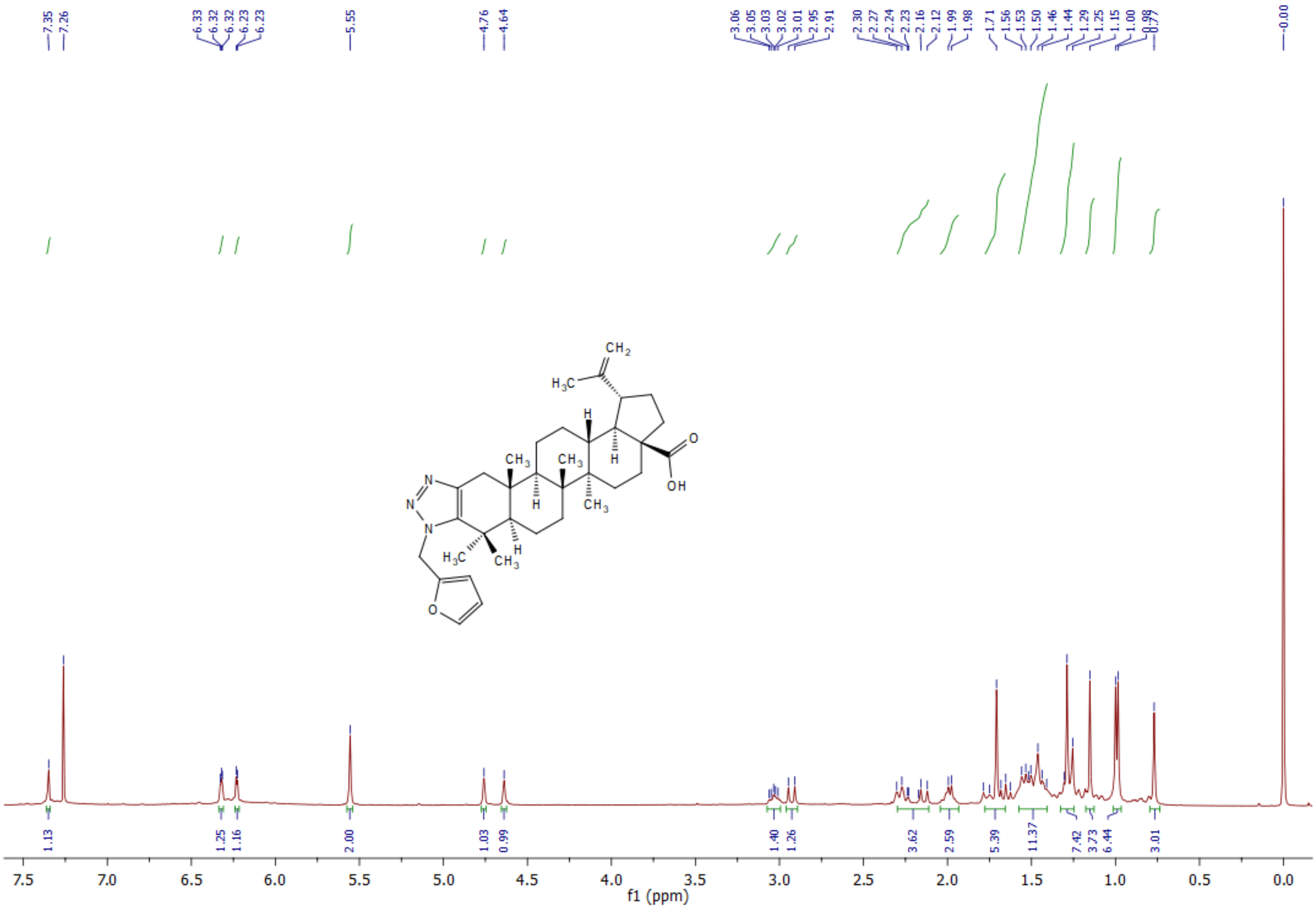

**¹³C NMR** spectrum of **5k** (400 MHz, $CDCl_3$):

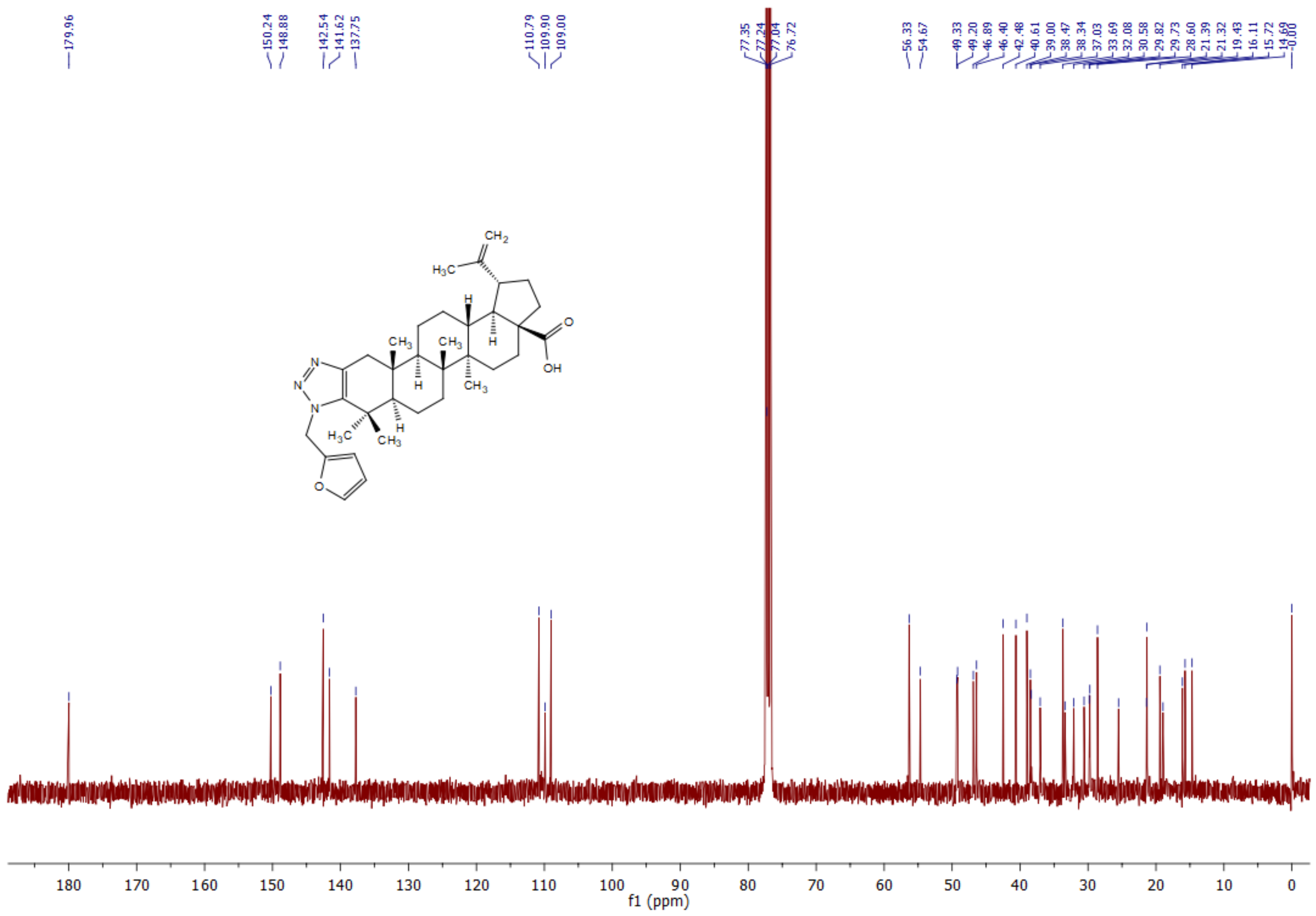

[1]**H NMR** spectrum of **5l** (400 MHz, DMSO):

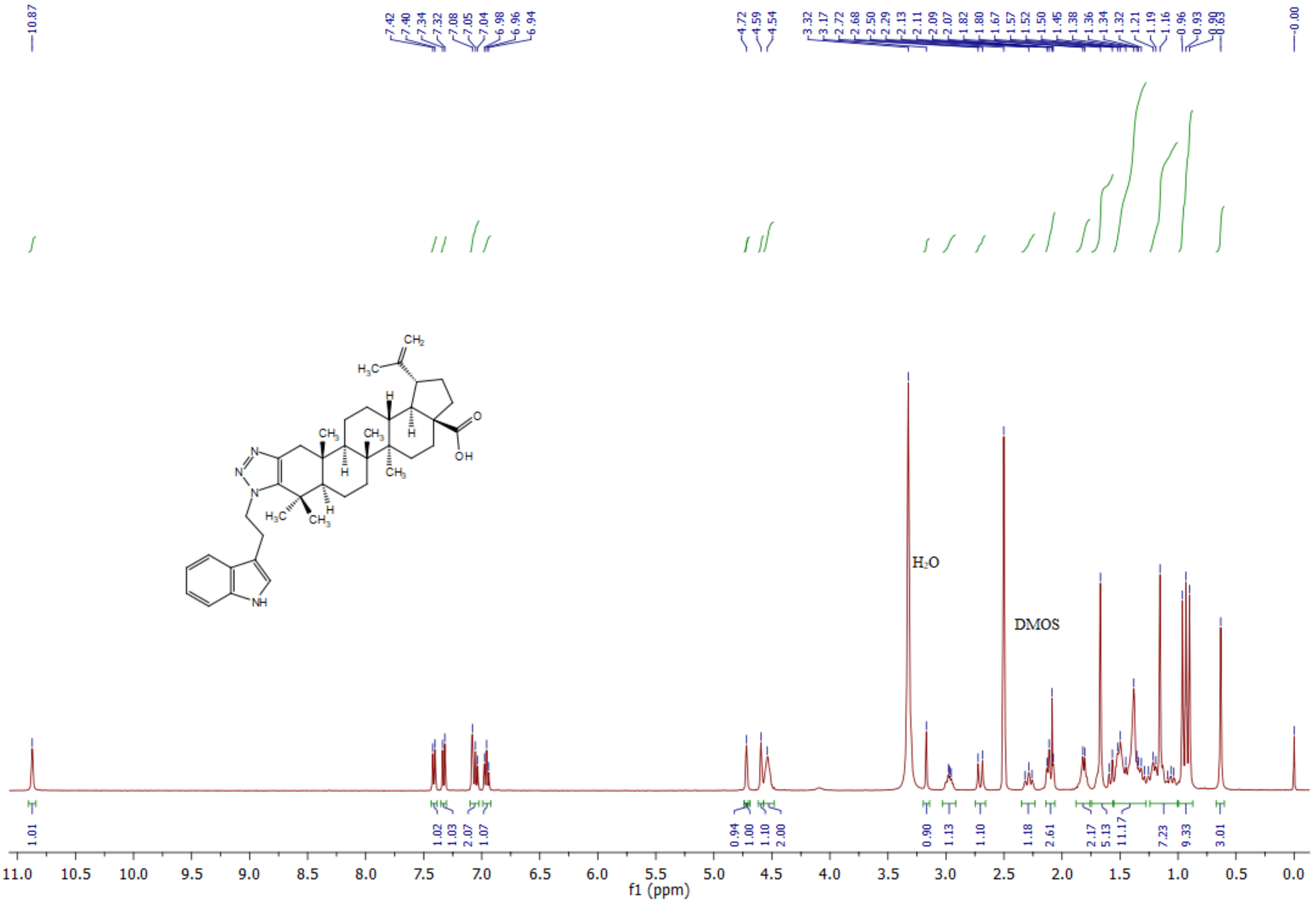

**$^{13}$C NMR** spectrum of **5l** (400 MHz, DMSO):

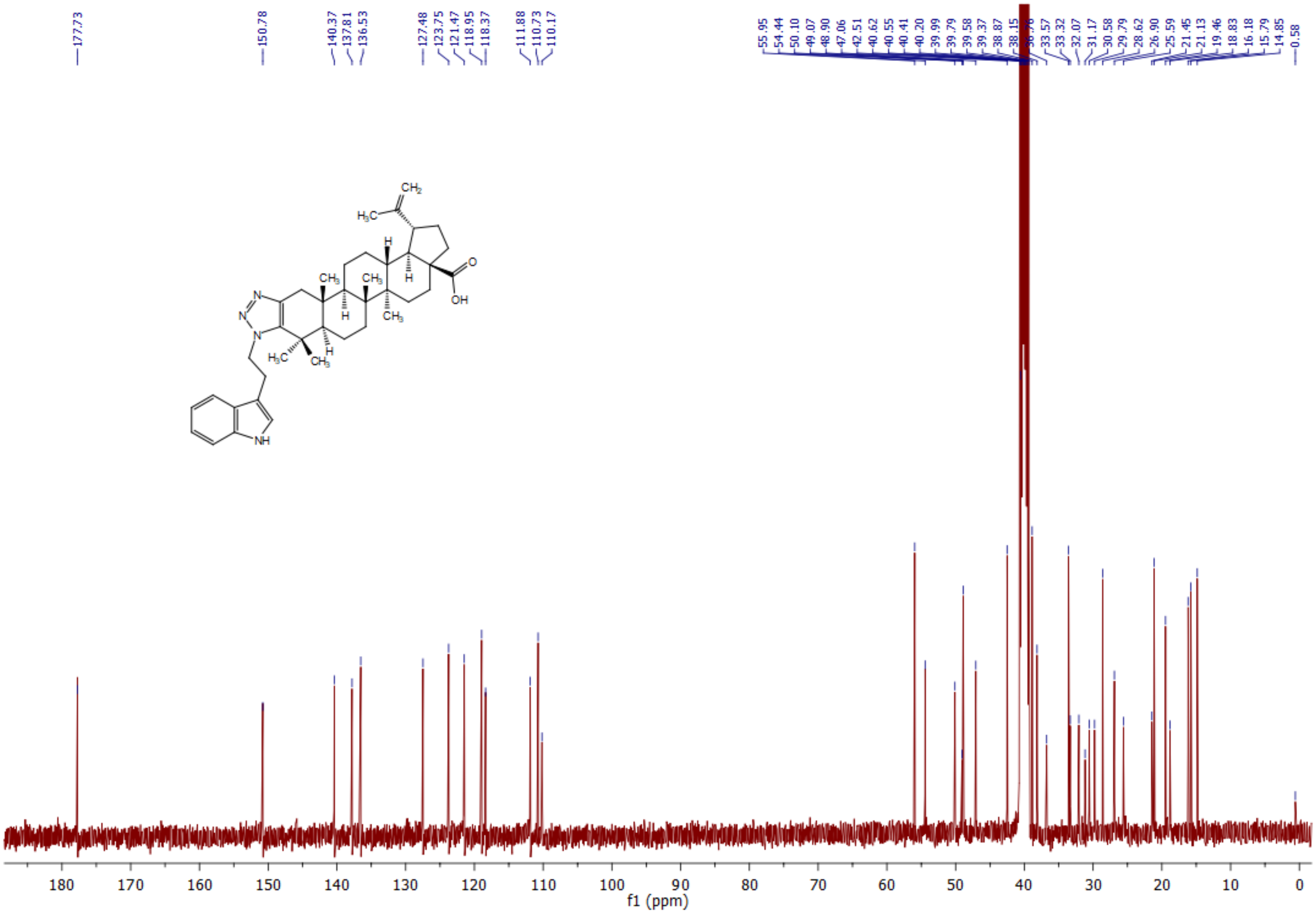

[1]H NMR spectrum of **5m** (400 MHz, DMSO):

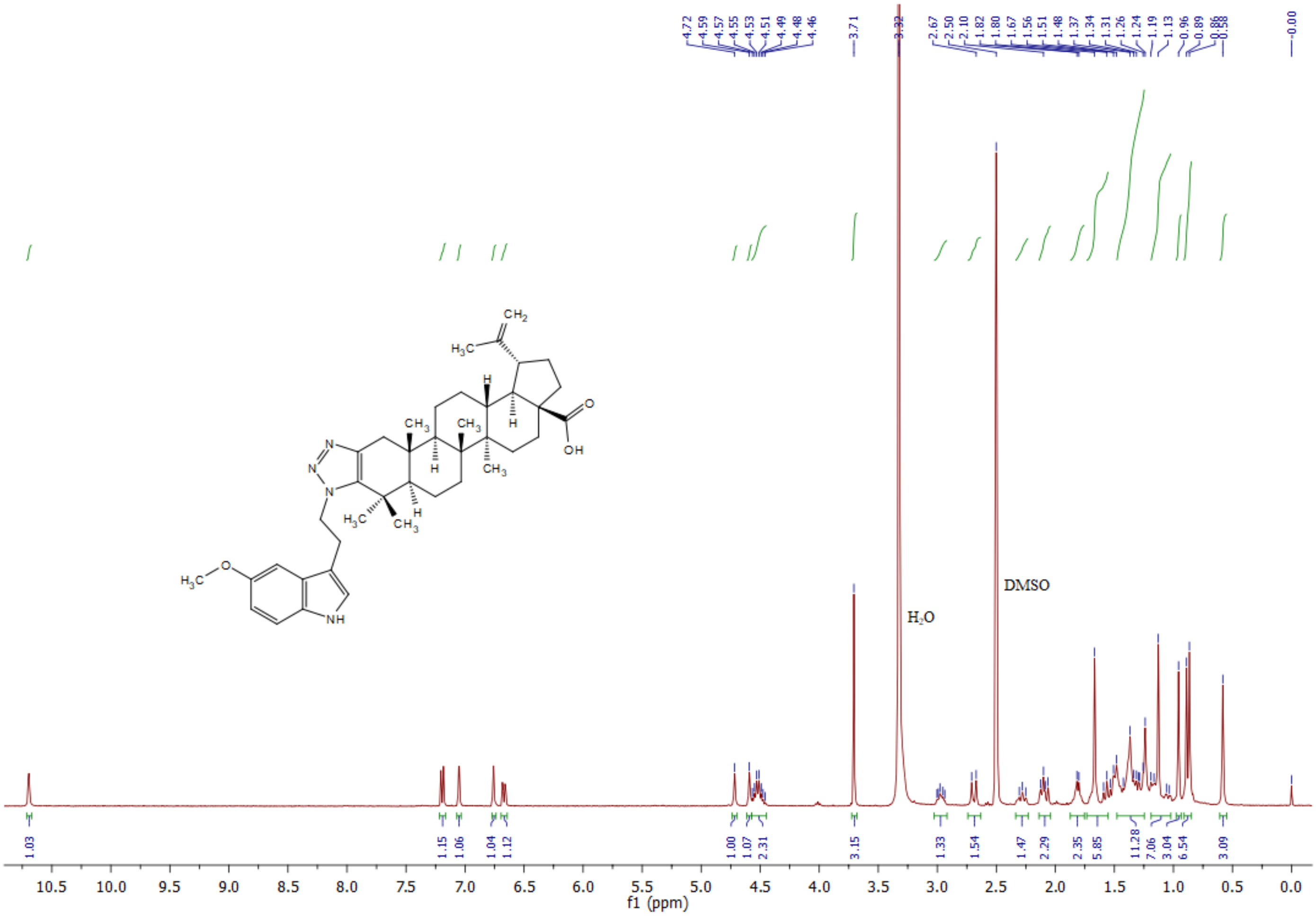

**$^{13}$C NMR** spectrum of **5m** (400 MHz, DMSO):

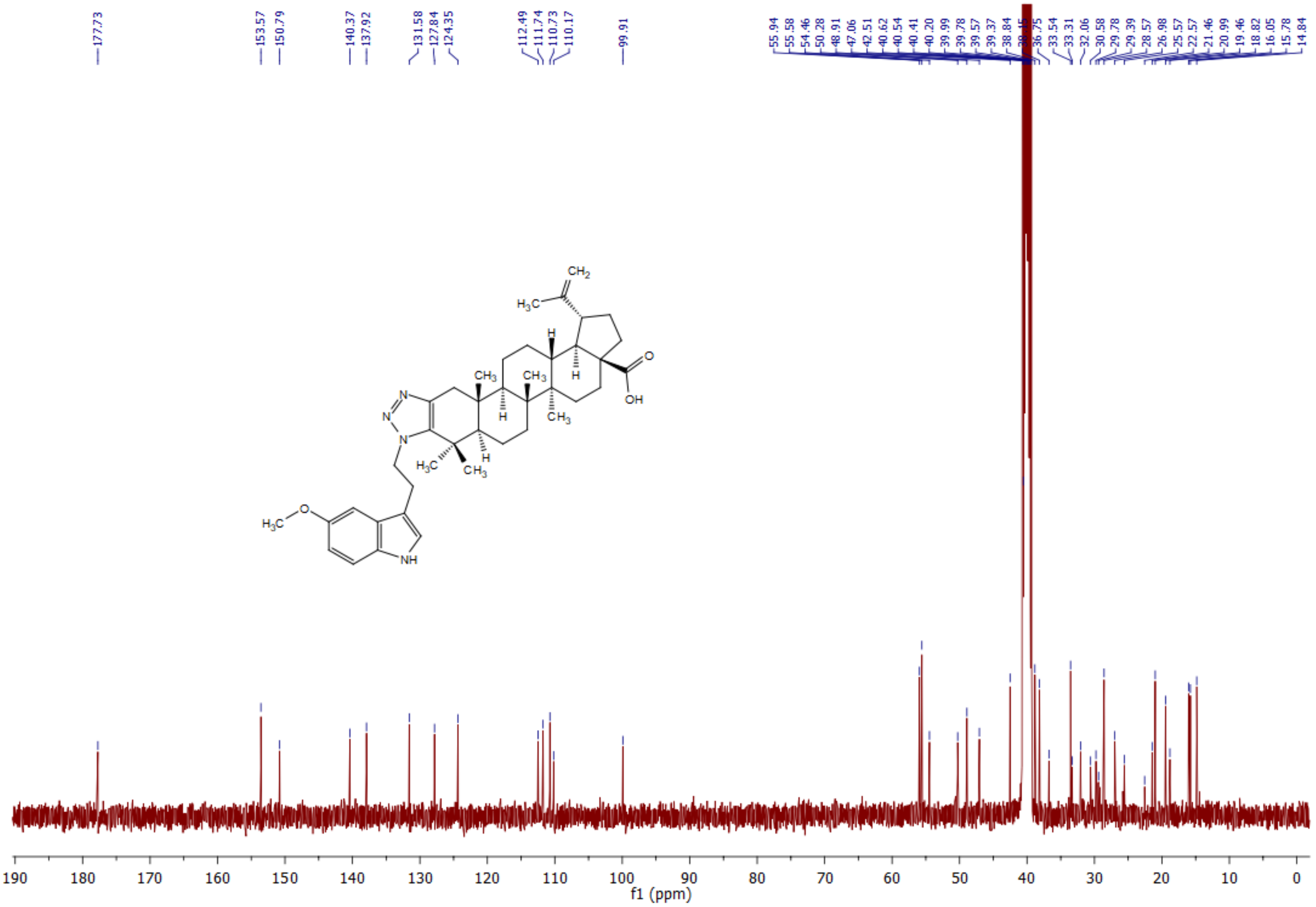

**[1]H NMR** spectrum of **5n** (400 MHz, $CDCl_3$):

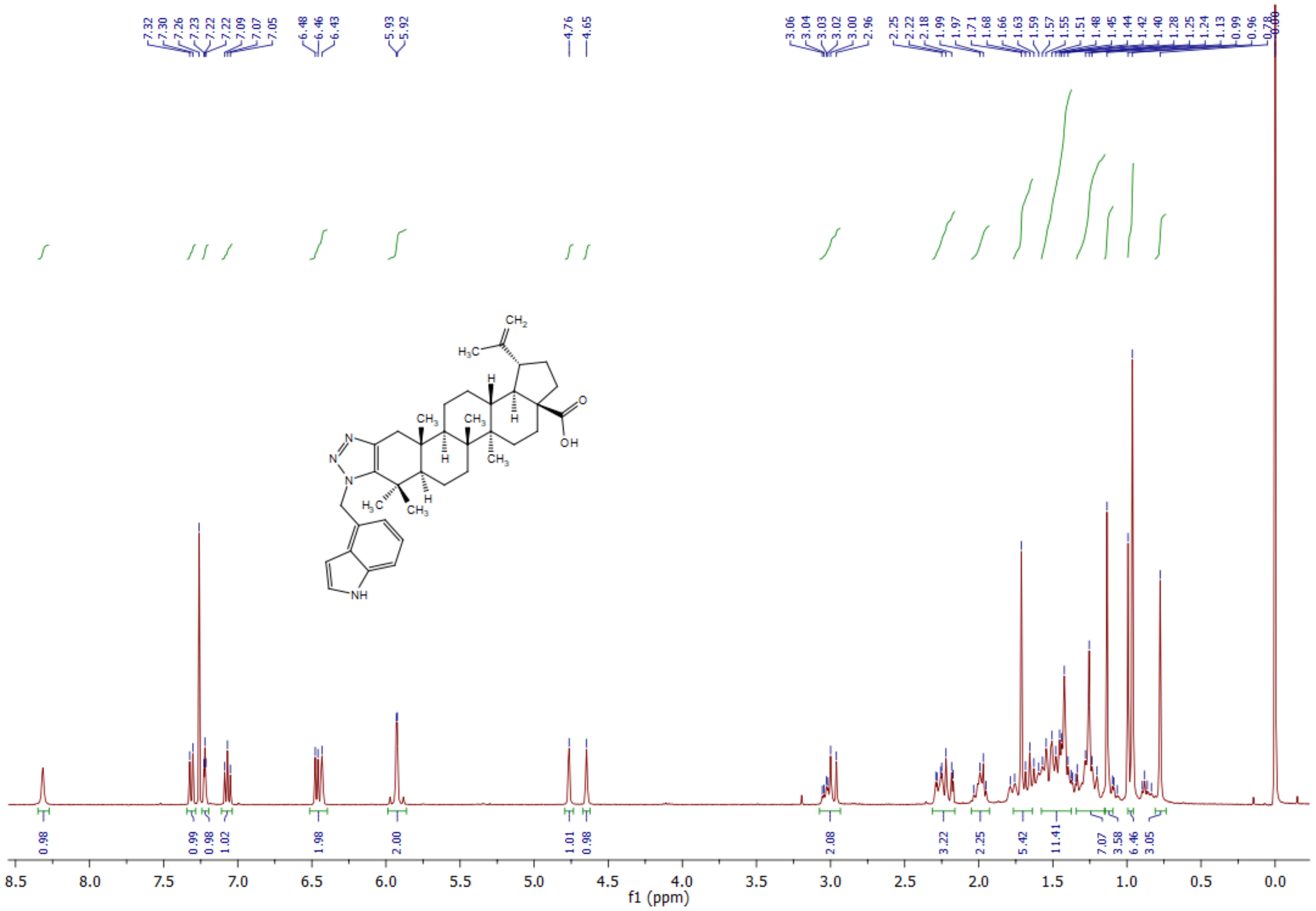

**$^{13}$C NMR** spectrum of **5n** (400 MHz, $CDCl_3$):

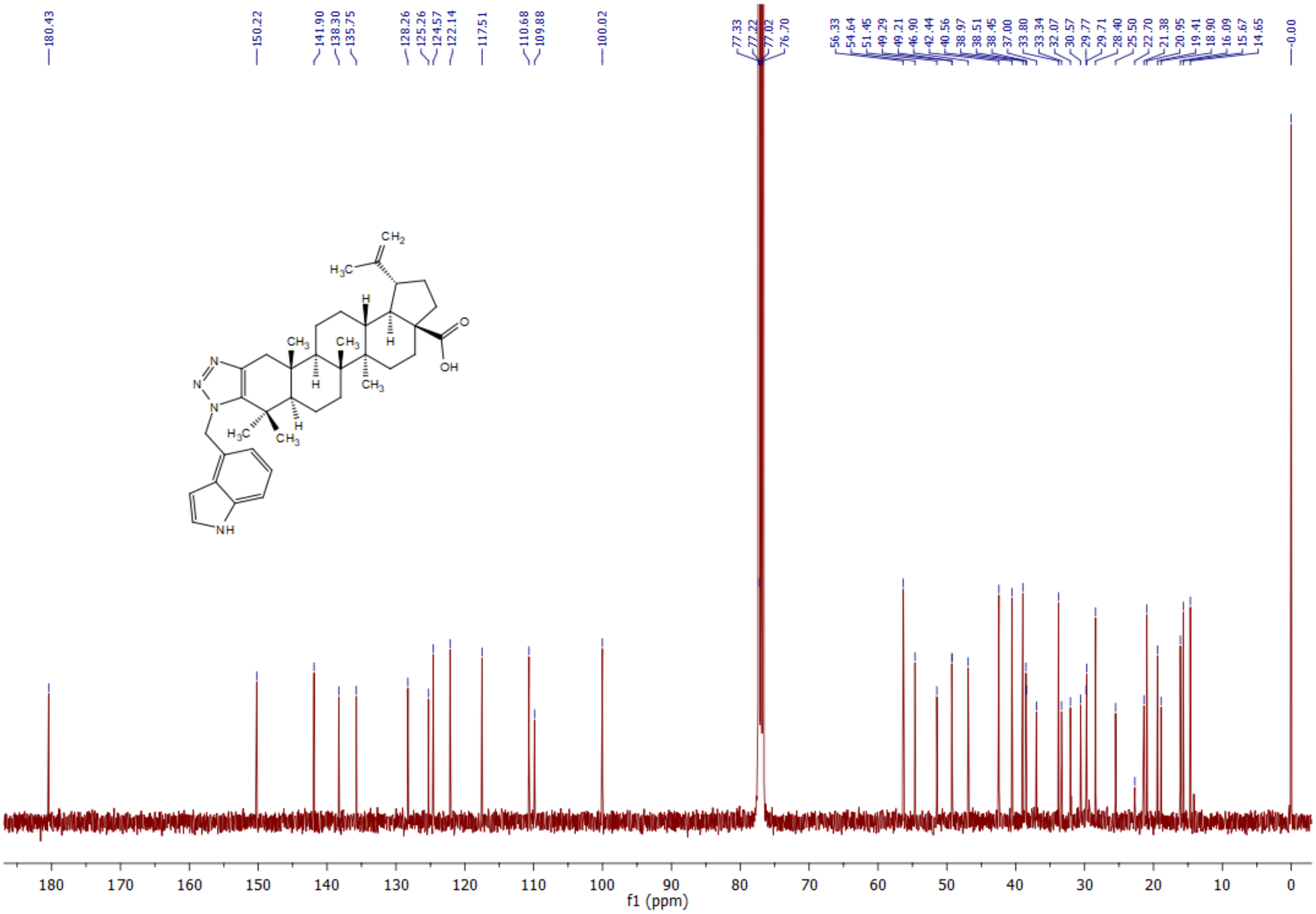

**$^{1}$H NMR** spectrum of **5o** (400 MHz, DMSO):

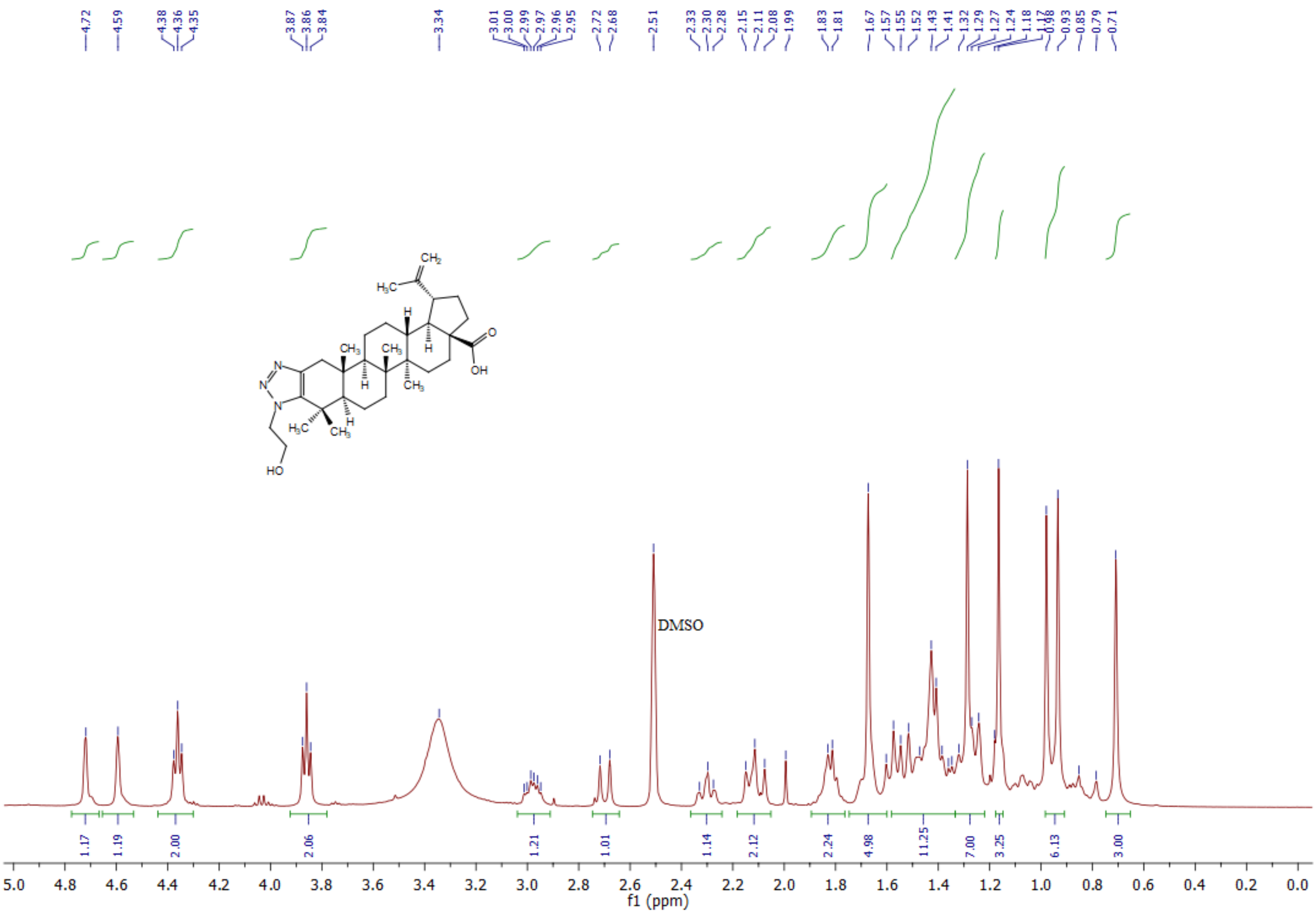

[13]C NMR spectrum of **5o** (400 MHz, DMSO):

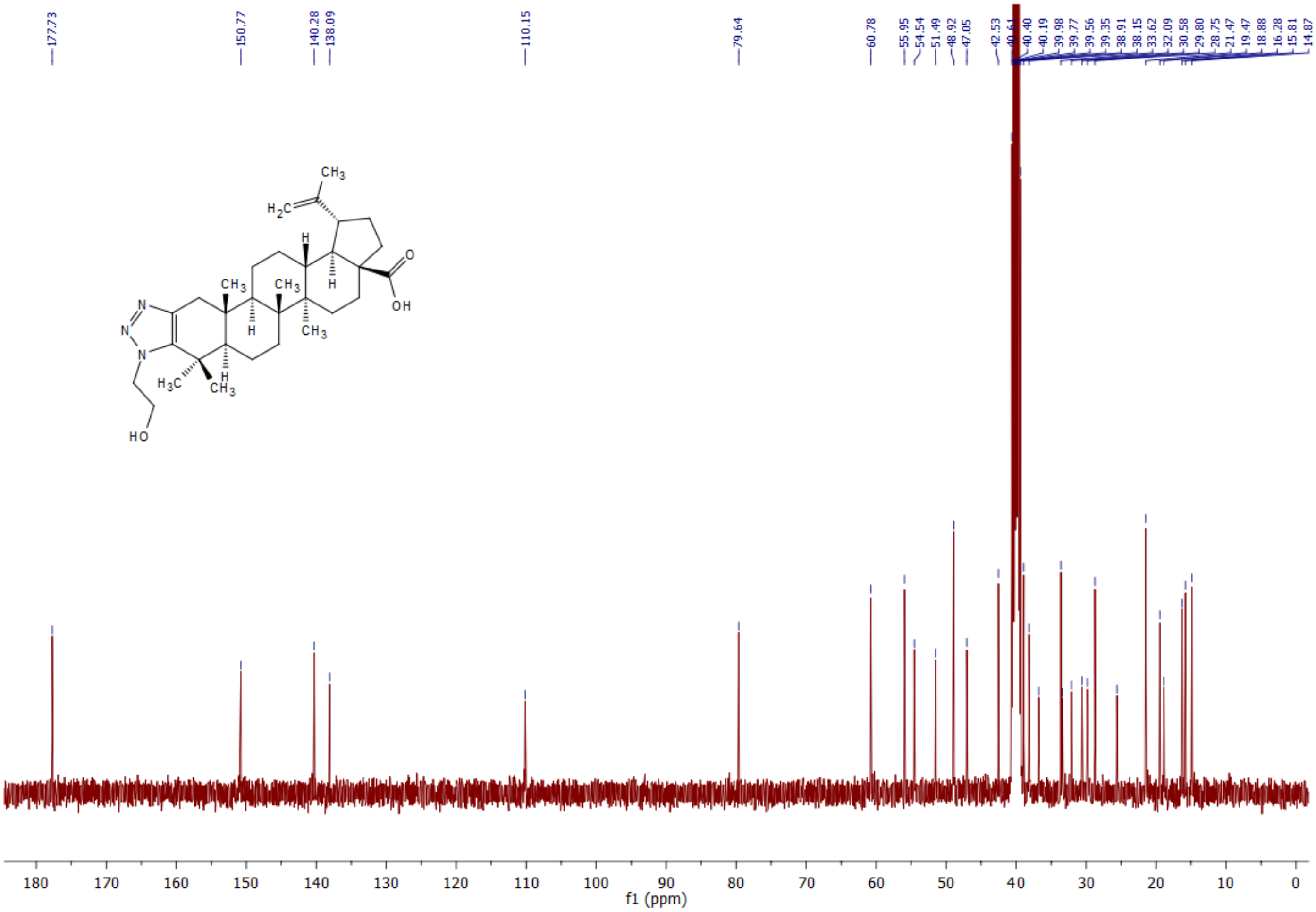

**[1]H NMR** spectrum of **5p** (600 MHz, $CDCl_3$):

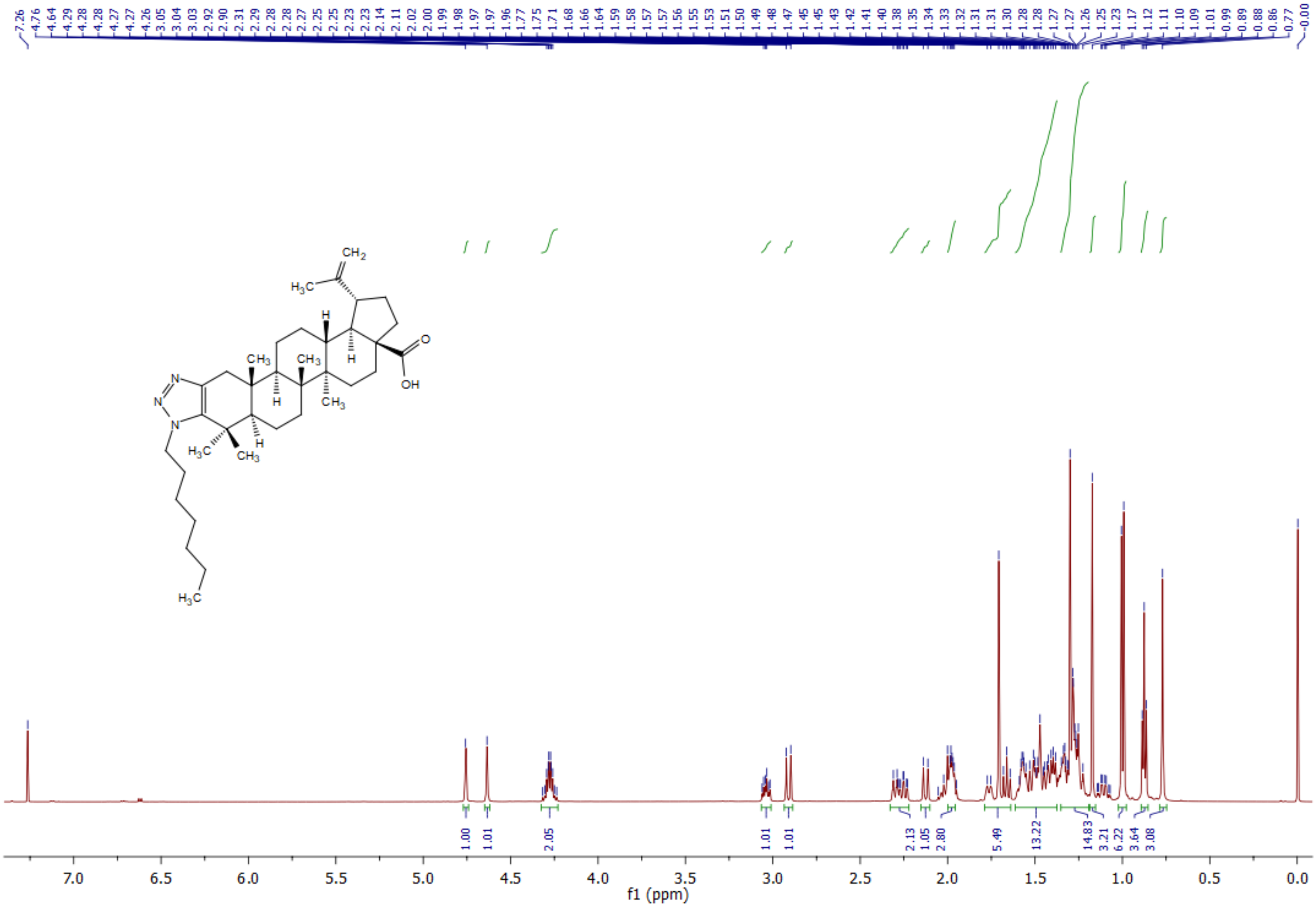

**$^{13}C$ NMR** spectrum of **5p** (400 MHz, $CDCl_3$):

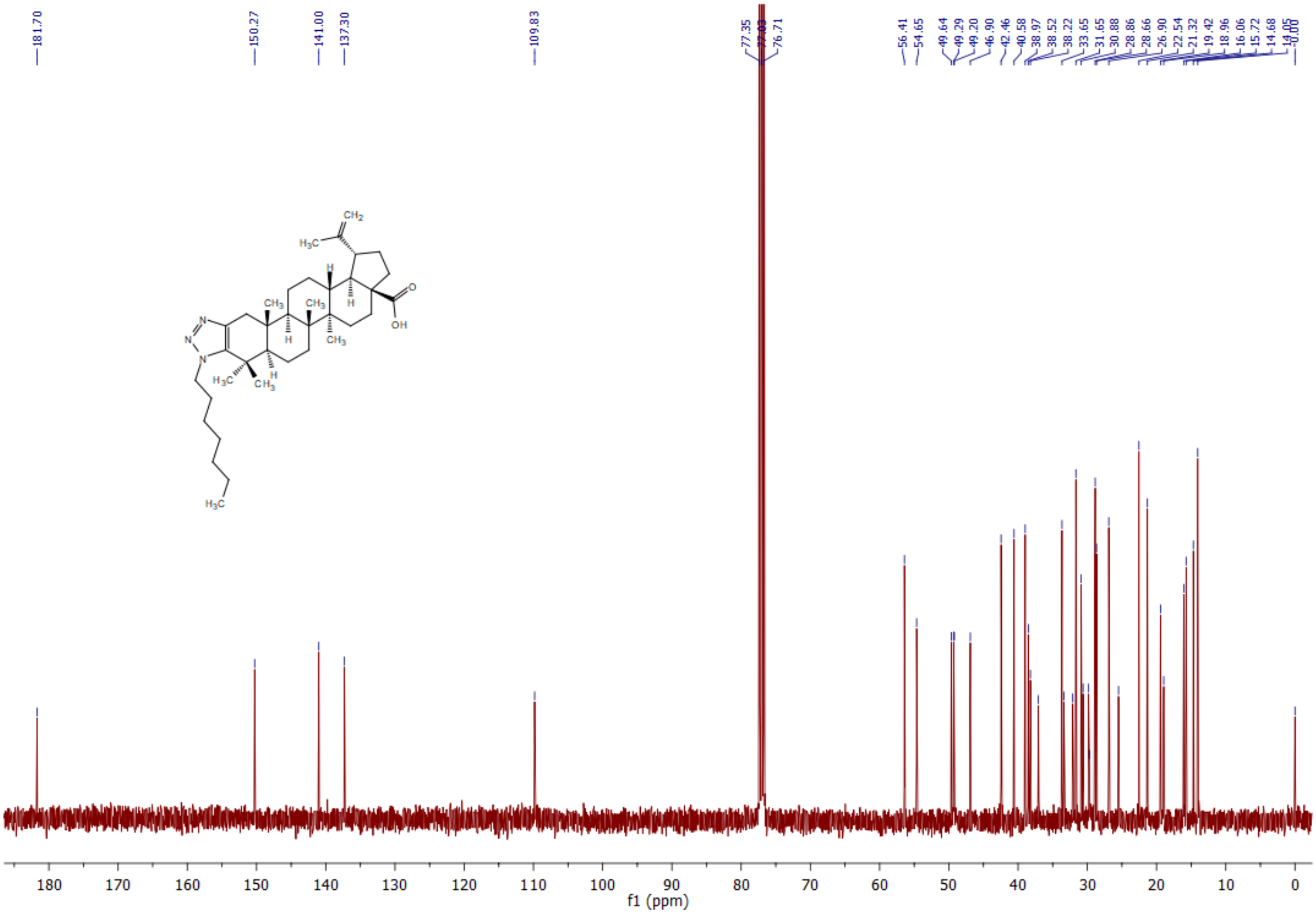

**$^1$H NMR** spectrum of **5q** (400 MHz, $CDCl_3$):

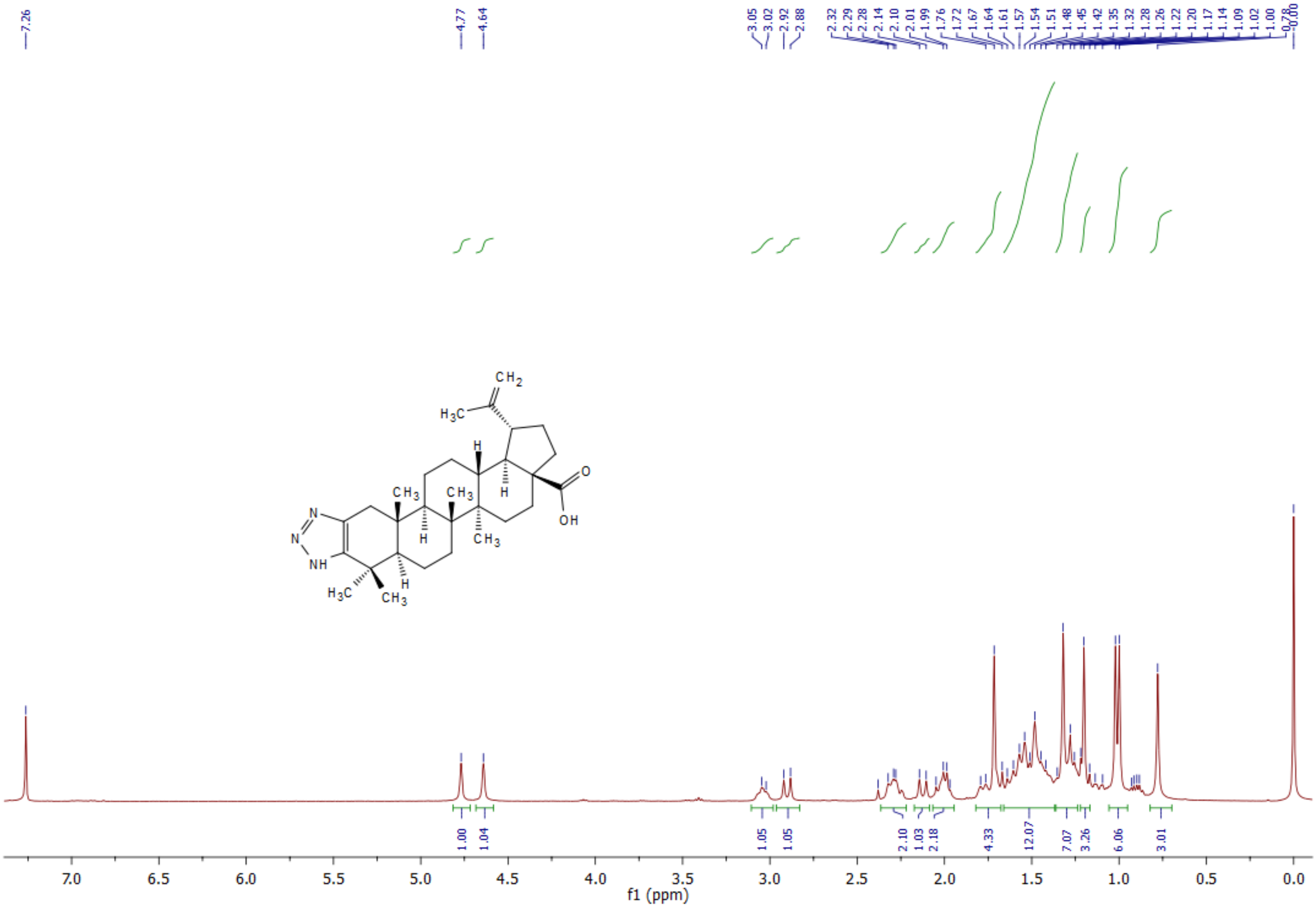

**$^{13}$C NMR** spectrum of **5q** (400 MHz, $CDCl_3$):

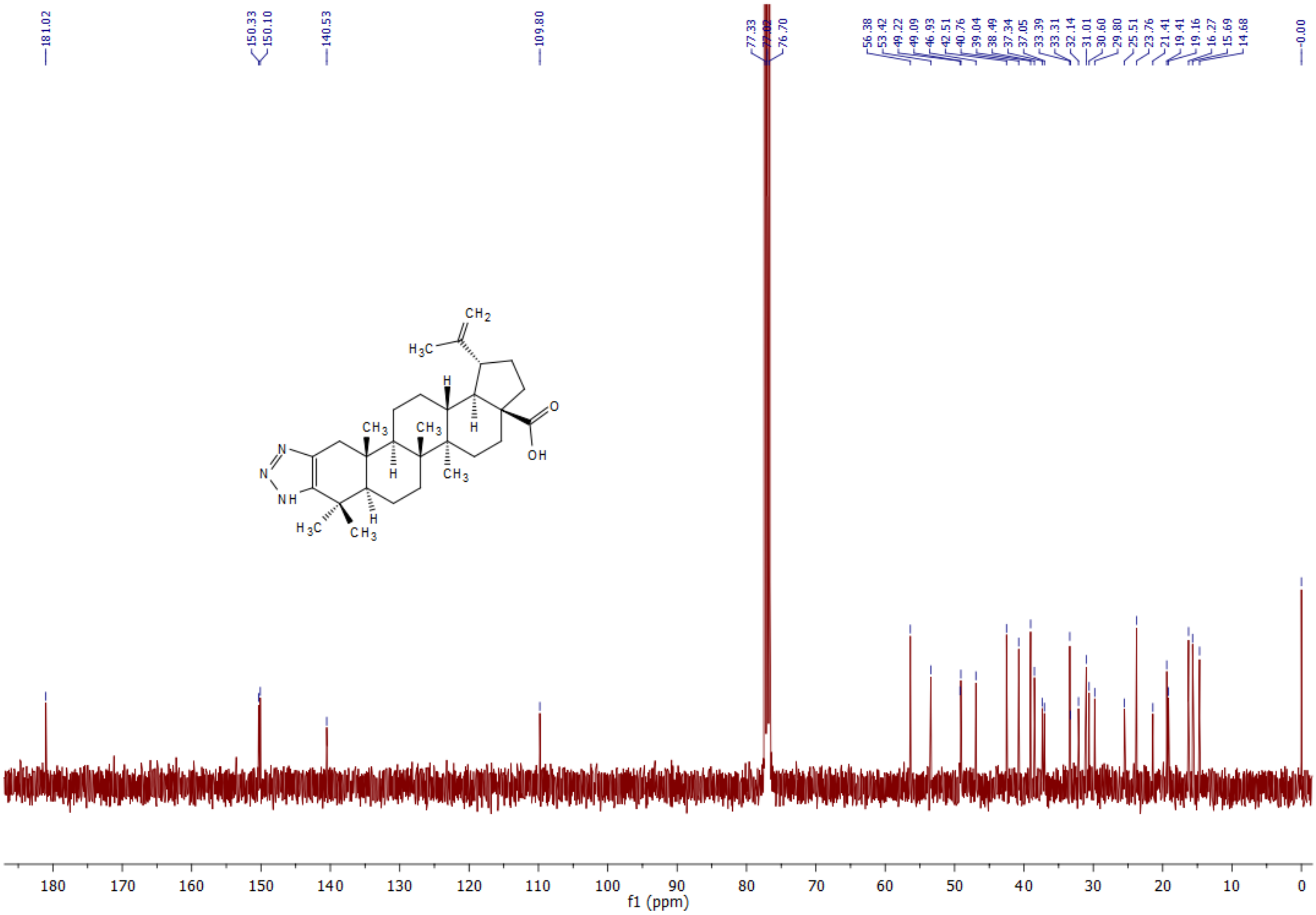

$^{1}$**H NMR** spectrum of **5r** (400 MHz, $CDCl_3$):

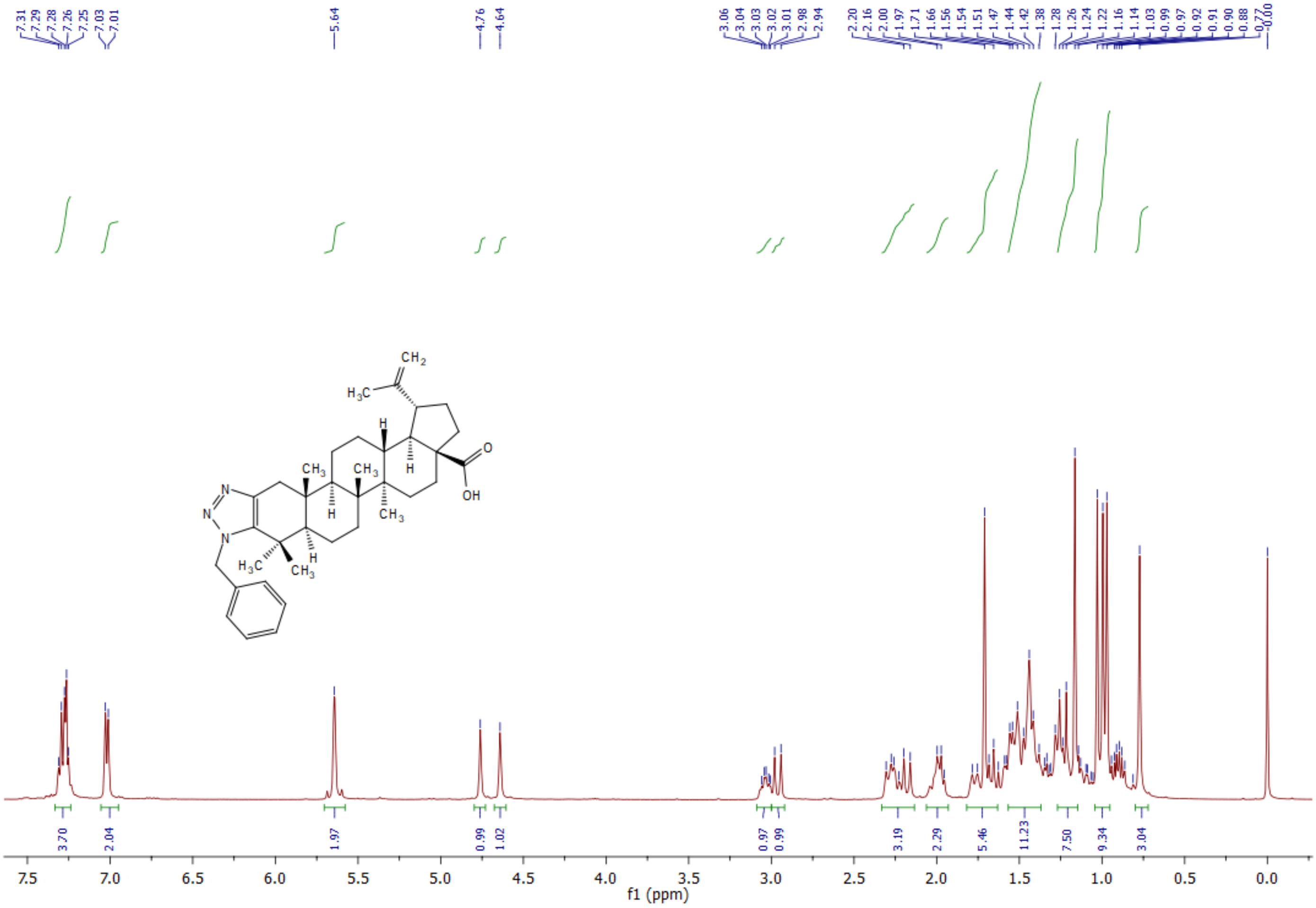

**[13]C NMR** spectrum of **5r** (400 MHz, $CDCl_3$):

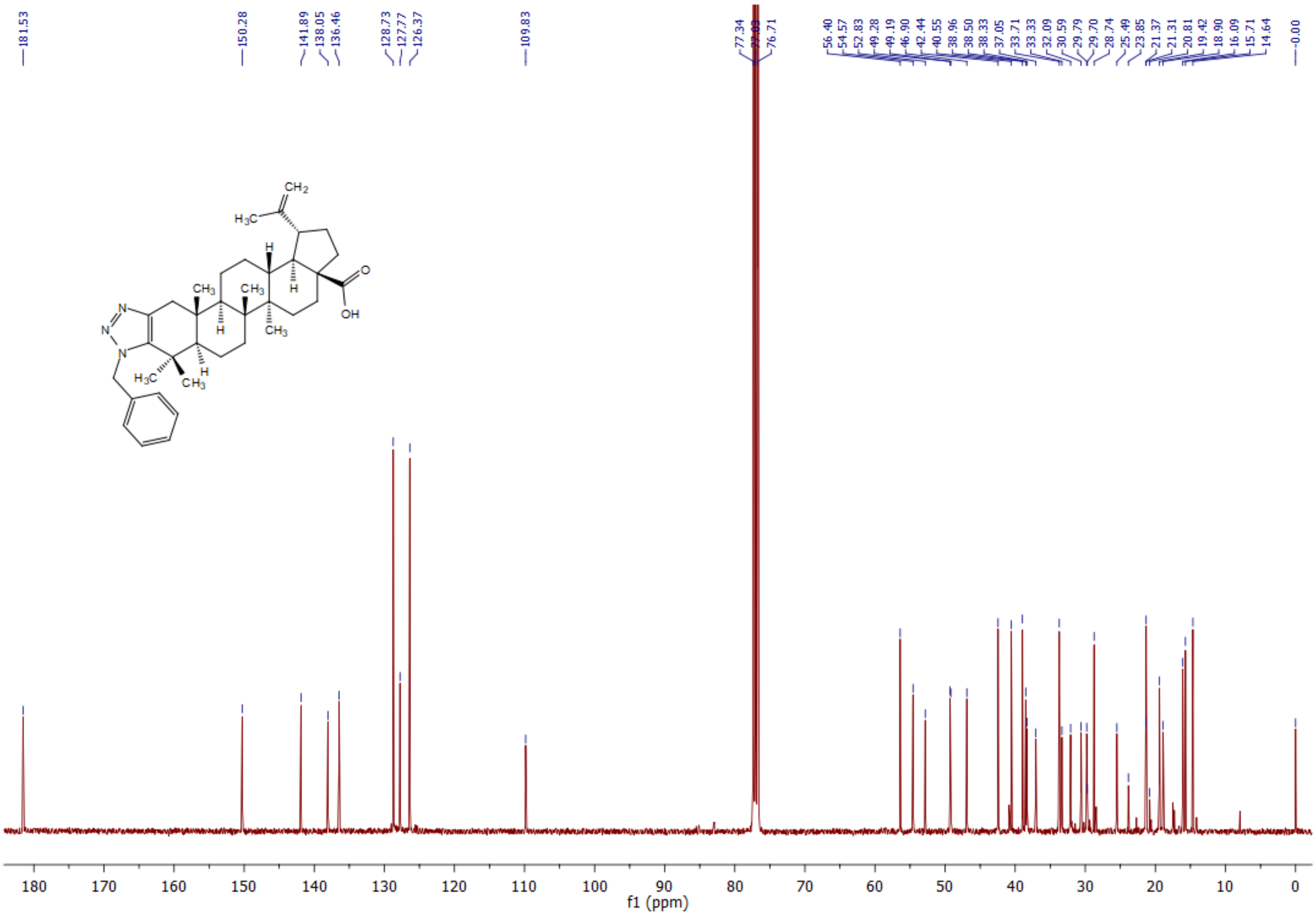

[1]H NMR spectrum of **5s** (600 MHz, $CDCl_3$):

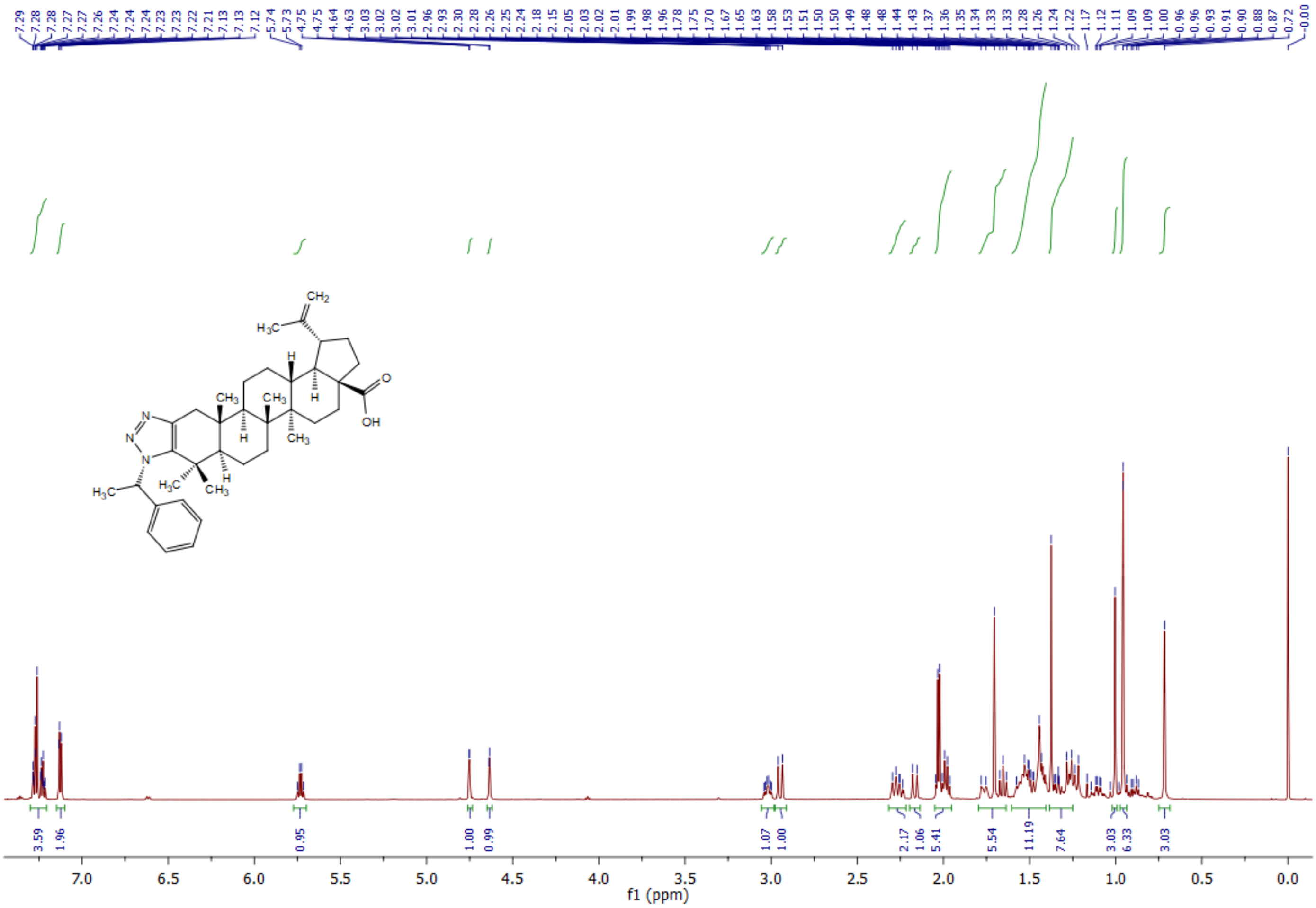

**$^{13}$C NMR** spectrum of **5s** (400 MHz, $CDCl_3$):

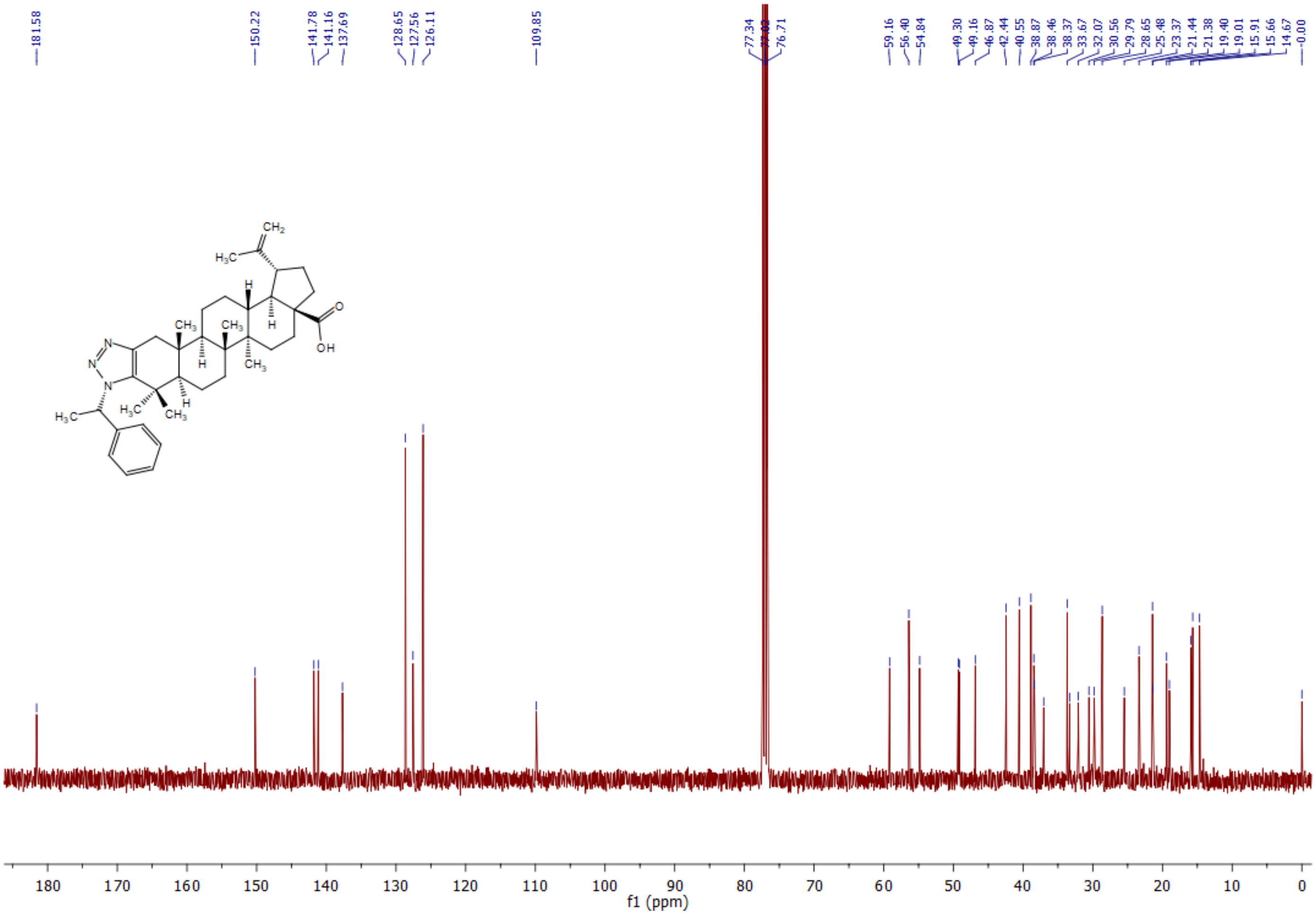

**$^1$H NMR** spectrum of **5t** (400 MHz, $CDCl_3$):

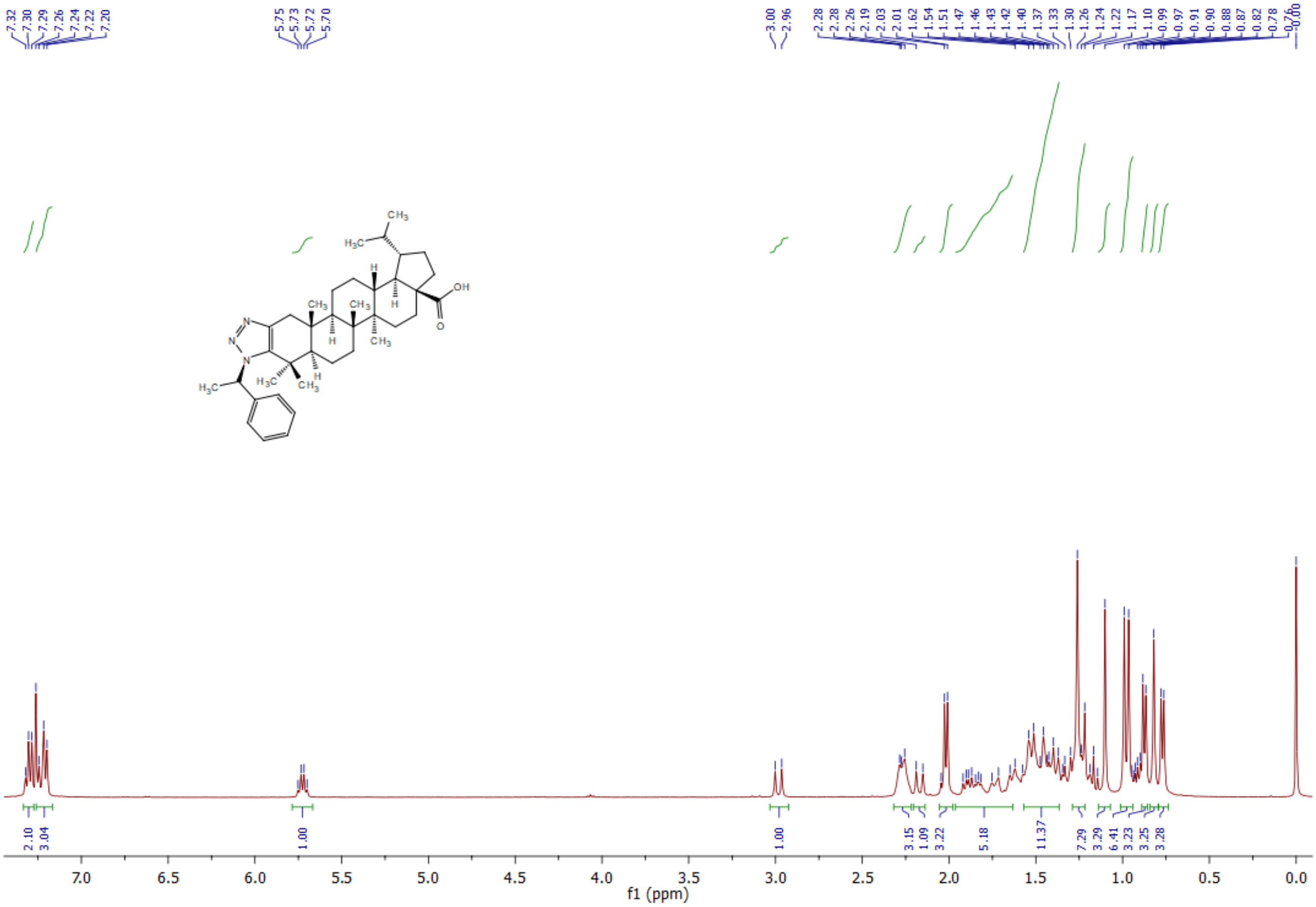

$^{13}$**C NMR** spectrum of **5t** (400 MHz, $CDCl_3$):

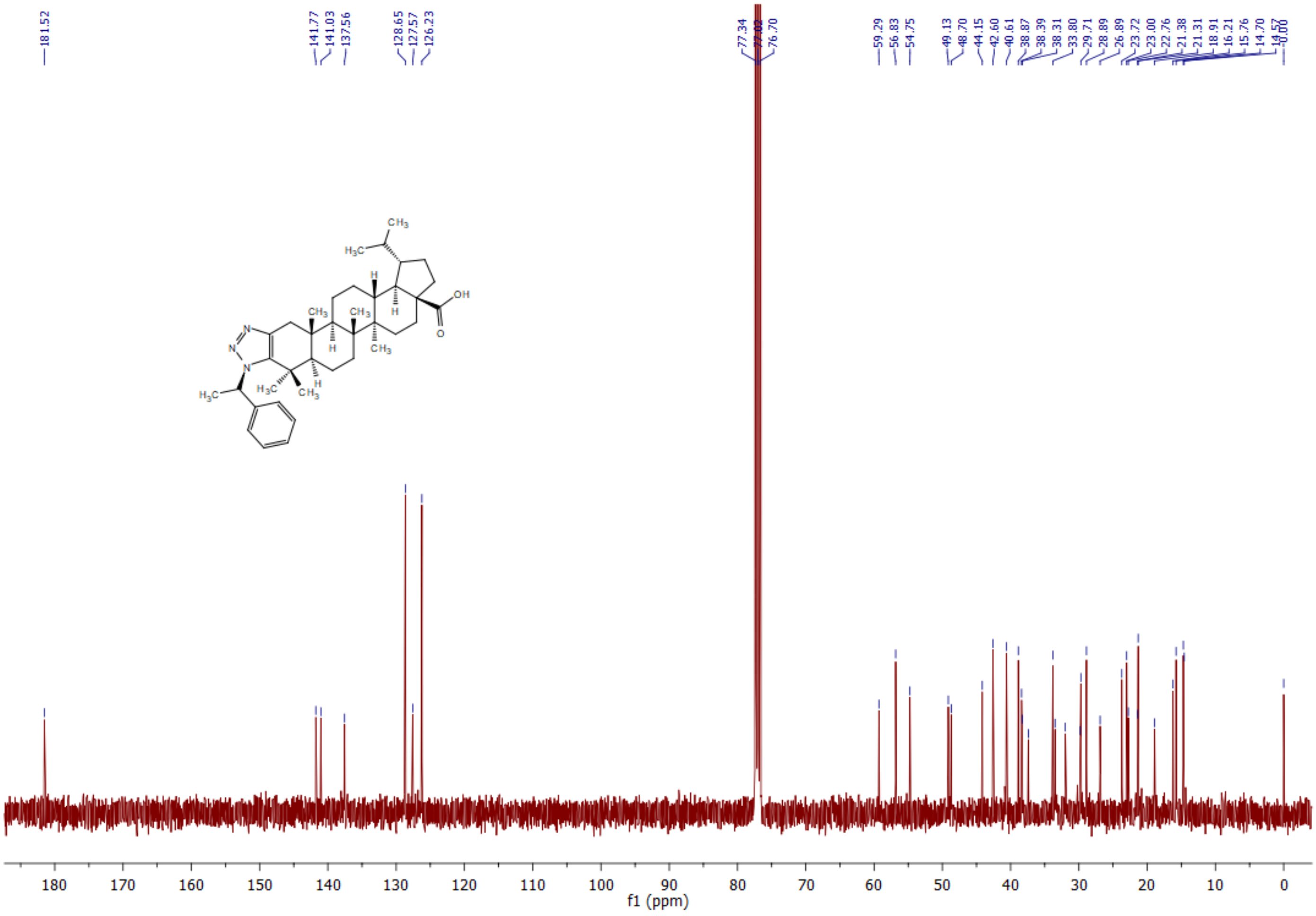

**$^1$H NMR** spectrum of **5u** (600 MHz, $CDCl_3$):

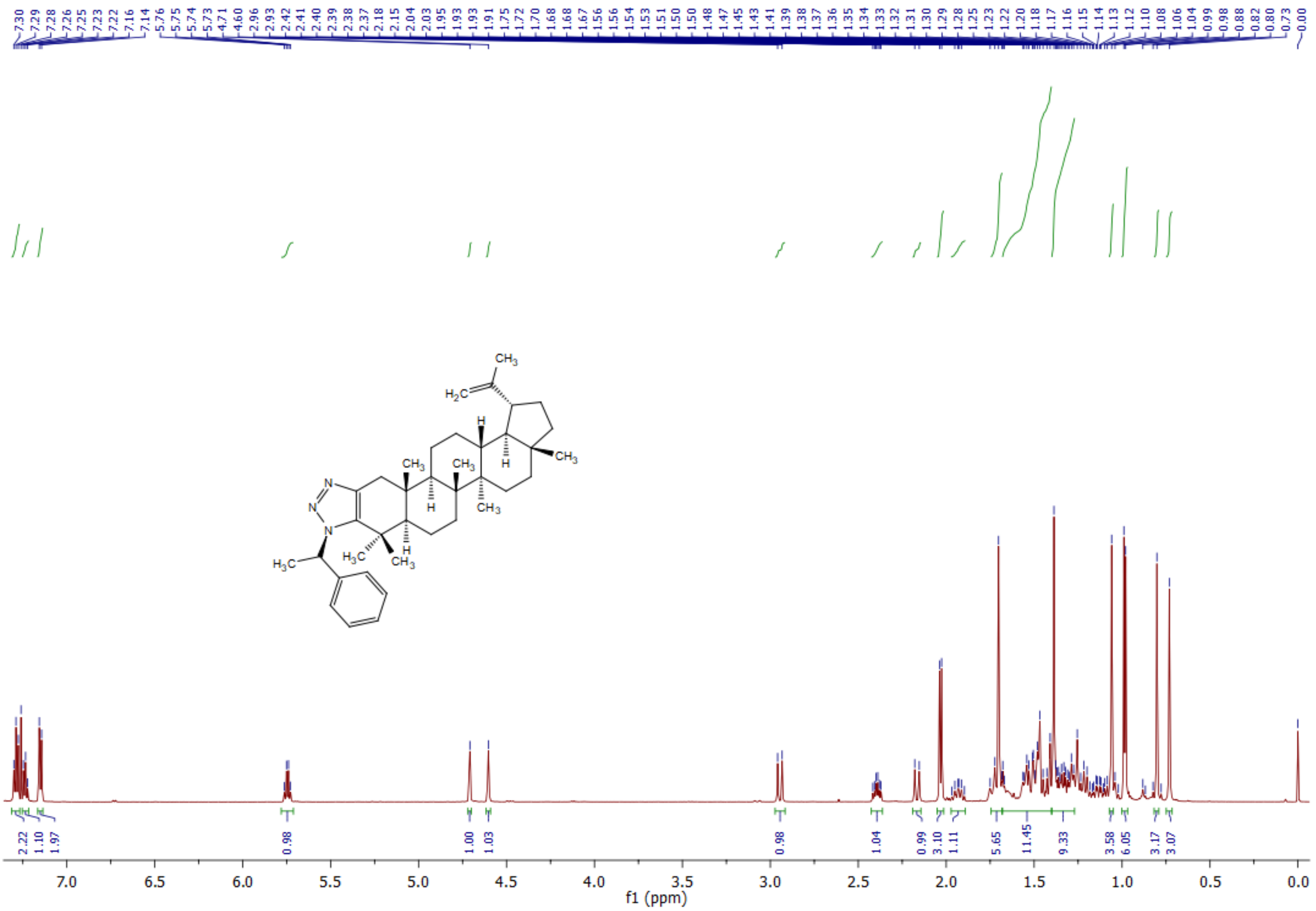

**$^{13}$C NMR** spectrum of **5u** (600 MHz, $CDCl_3$):

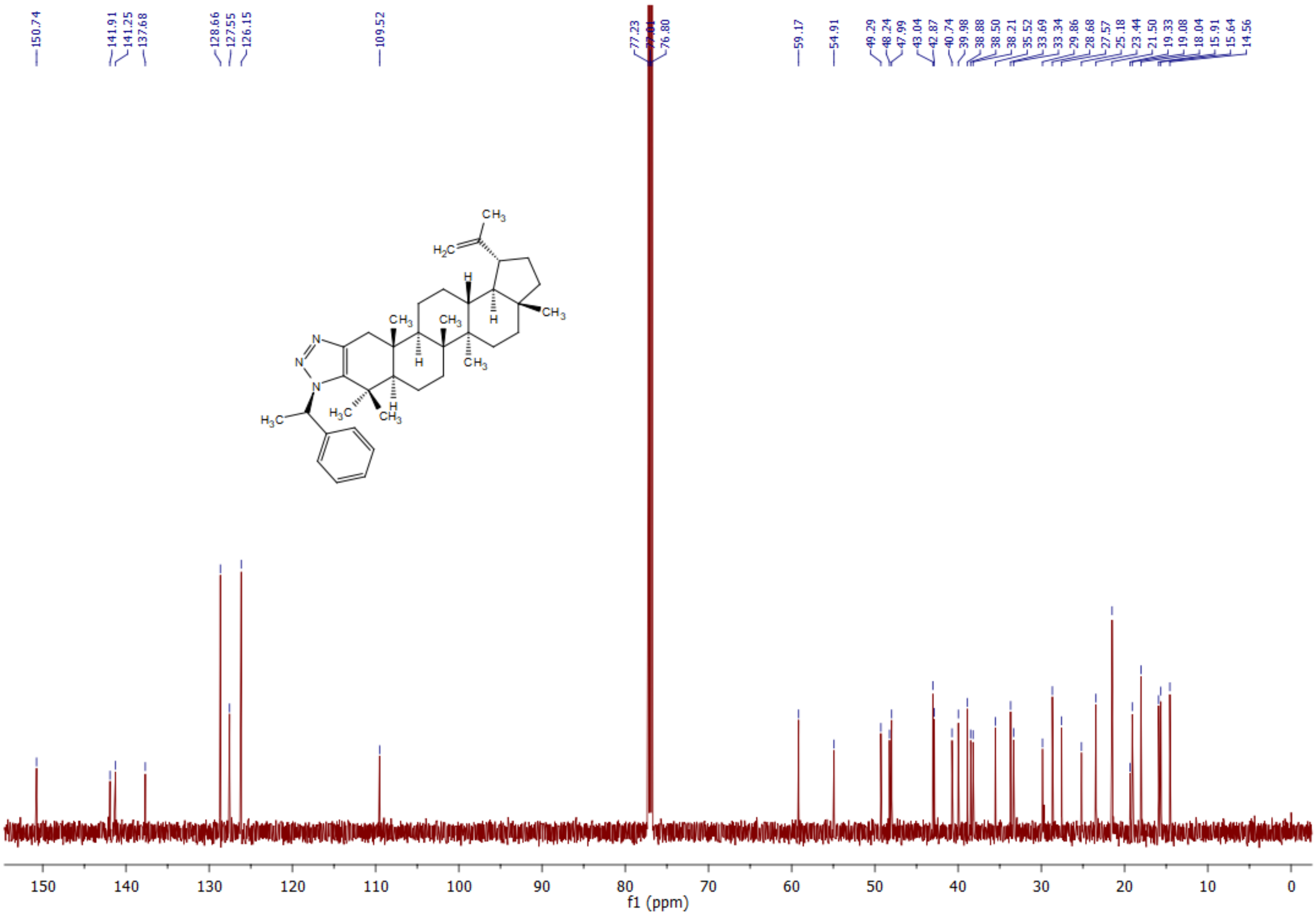
